# Extensive Benchmark Study of the Resonance Raman Spectra of Lumiflavin

**DOI:** 10.1101/2025.03.25.642206

**Authors:** Prokopis C. Andrikopoulos, Heba Halimeh

**Affiliations:** Institute of Biotechnology of the Czech Academy of Sciences, BIOCEV, Průmyslová 595, CZ-252 50 Vestec, Czechia; Charles University, 1^st^ Faculty of Medicine, BIOCEV, Průmyslová 595, CZ-252 50 Vestec, Czechia; Rhine-Waal University of Applied Sciences, Marie-Curie-Str.1 D-47533 Kleve, Germany

**Keywords:** Flavins, DFT, Density Functional Theory, Raman, QM, Resonance, Lumiflavin, LOV domains, BLUF, Benchmark, FSRS

## Abstract

An extensive computational TDDFT resonance Raman study is presented here including forty-two different DFT functionals. The functionals were checked against the experimental FSRS Evolution Associated Spectrum of the equilibrated S_1_ state of FMN published earlier. Off-resonance spectra were computed first and yielded adequate agreement with the experimental spectrum. Fine-tuning of the correlation was achieved with the inclusion of specific scaling factors for each DFT functional, aimed to align the highest computed peak (symmetric C=O stretch) to the corresponding experimental peak in the fingerprint region. Subsequently, resonance Raman intensities were calculated with a careful choice of the resonant states. The experimental Evolution Associated Spectrum utilized for the comparisons was taken under resonance conditions, hence the inclusion of resonance enhancements in the calculations improved the agreement in most of the DFT functionals. For six particular DFT functionals, namely HCTH/407, OLYP, OPBE, O3LYP, tHCTHhyb and TPSSh, the theoretical/experimental correlation was particularly facile, and their agreement was predicted superior to the other functionals. The narrowing down to the above set was achieved by the evaluation of all DFT functionals according to five criteria ranging from the percent error of the main flavin excitations to the experimental values, to visual inspection of the spectra, and the determination of whether the inclusion of resonance in the calculation improved the agreement with experiment. Owing to the extent of the data set, valuable insights were gained to assist similar studies.

## Introduction

Flavins are a family of isoalloxazine-containing chromophores that are present as co-factors in many photosensitive proteins.^1-3^ Members of the family include lumiflavin, riboflavin, flavin mononucleotide (FMN) and Flavin adenine dinucleotide (FAD). The difference between the members depends on the substitution of the N_10_ atom of the isoalloxazine ring, from methyl in lumiflavin, up to a combination of a sugar chain, phosphate group and nucleotide base in FAD (**Scheme 1**). Lumiflavin in particular, is broadly used as a computational analogue for the more substituted members of the family in many photochemical,^4-7^ as well as in benchmark studies.^8^

To study experimentally chromophore systems in solution, or embedded in photoproteins, one of the techniques of choice is time-resolved vibrational spectroscopy. Transient Absorption (TA), ultrafast transient Infrared Spectroscopy (TRIR), resonance Raman (rR) and other time-resolved spectroscopic techniques offer a wealth of information on the light response of photochemical and photobiological systems.^9-12^ Femtosecond Stimulated Raman Spectroscopy in particular (FSRS),^13^ has unique advantages owed to its ultrafast resolution and tunability, which entails the targeting of specific electronic transitions of the target chromophore, under resonance conditions. When these conditions are met, the obtained spectra exhibit fewer features than their off-Resonance counterparts, with enhanced signal strength.^14-16^ Notwithstanding the above advantages, spectra obtained under resonance pose an additional challenge in their interpretation. In broad terms, the normal modes pertaining to the electronic transitions in tune with the incident light are promoted in their intensities, while other signals are supressed, and this has to be taken to account in their analysis.

Theoretical calculations have proven to be indispensable, not only in the interpretation of time-resolved spectroscopy,^10, 17-19^ but also in aiding the design of photobiological molecules with specific properties.^6, 20^ For resonance Raman spectroscopy in particular, the time dependent theory of resonance Raman spectroscopy (TD-RR) is currently implemented in computational chemistry codes, including Herzberg-Teller contributions and solvent and anharmonic effects.^21^ To compute the spectra, two different approaches exist, either requiring the optimization of the resonant state, or utilizing only the gradients of the excitations in the geometry of the reference state, yielding similar results.^22^ Several computational studies have appeared recently in the literature, tackling resonance Raman calculations of various systems ranging from thiophene derivatives to fluorescent protein chromophores, including often a few DFT functional in their studies.^22-25^ The popular density functional theory (DFT) functional B3LYP^26-27^ is used extensively for the computation of flavin-containing systems, due to the accurate prediction of the excitation energies.^9, 17^-^18, 28-29^ However, as Green *et al* demonstrated with Flavin Mononucleotide (FMN, Fig.2A),^30^ the calculated B3LYP off-Resonance spectra of lumiflavin correlate far better with the experimental FSRS – obtained under resonance conditions - than the calculated resonance Raman spectra.

In this study, we present a comprehensive approach to evaluate the off-Resonance (offR) and resonance Raman spectra (rR) of a plethora of DFT functionals against the FSRS experimental spectrum of FMN. We included a total of forty-two DFT functionals, with lumiflavin as the target compound, benchmarked against the experimental Evolution Associated Spectrum of 1FMN*,^16^ obtained under resonance conditions, and assigned to the equilibrated first excited singlet state (S_1_). The choice of a modest double-ζ basis set for the majority of the calculations allows the findings presented here to be applicable for larger cluster or QM/MM studies of lumiflavin embedded in a protein environment, where a higher basis set would not be affordable. The functionals were scored against: (i) the predicted excitation energies, (ii) the percent error of the correlation between experimental and rR-calculated spectra, (iii) the difference in percent errors between the offR and rR correlations - indicating whether the agreement improved or deteriorated with the addition of resonance, (iv) the predicted rR intensity of the strongest experimental peak in the fingerprint region at 1498 cm^-1^ and finally, (v) the visual inspection of the spectra, to determine whether the experimental/theoretical spectral profiles are compatible, facilitating the assignment.

The aim of this benchmarking effort is not to delve deep into the particulars of each DFT functional, but instead to narrow down the chosen large set of DFT functionals to a smaller set that can be used to describe more faithfully flavin-related systems with resonance Raman calculations. The insights gained from this benchmark study can hopefully be transferable to other chromophore/photoprotein systems.

### Computational Details

All calculations were performed with the Gaussian program (G16 Rev. C.01).^31^ A total of forty-two DFT functionals were utilized, as shown in **Table S1** together with the percentage of Hartree Fock exchange, short description and references to original articles. The empirical dispersion correction parameters employed in this study are shown in **Table S2**.^32-33^ Empirical corrections were included for all but three of the functionals, namely SOGGA11, VSXC and LSDA(SWVN). Regarding the frequency scaling factors, no comprehensive study exists in the literature for excited states. Consequently, the available scaling factors for ground state calculations were utilized, as described in **Table S3**.^34-36^ For the functionals that scaling factors were unavailable, the FREQ program by Truhlar and co-workers was used to create them for this study, using the full scale factor optimization model.^37-39^ Both literature and scaling factors derived from FREQ will be denoted as **Sc**_**L**_ further in the text. An additional scaling factor was devised called Specific Scaling Factor (**Sc**_**S**_, see **Table S3**), which aligns the computed S_1_ state symmetric C=O stretch of each DFT functional (v_75_) with the experimental FMN peak at 1626 cm^-1^.

Due to the extended benchmarking, all calculations employed either the modest double-ζ basis set cc-pVDZ or its expansion including diffuse functions, aug-cc-pVDZ.^40-41^ The latter always produced excited states within the experimental resonance window, set at the wide range of 750-800±100 nm,^16-17, 30^ which was not always the case for the modest basis set. Specifically, for B3LYP, a functional that has been associated with numerous computational studies of flavins,^17-18, 30^ a basis set dependence/convergence study is included from up to the quadruple-zeta cc-pVQZ,^40-43^ together with the equivalent augmented sets (up to triple-ζ), ranging from 326 up to 1405 basis functions. The Polarizable Continuum Model (PCM) was used as the solvation method in all calculations with water as the solvent. Equilibrium solvation was used for both ground state^44-46^ and excited state optimizations.^47^

Vertical excitations, simulations of ground-state UV–vis spectra, and excited state optimisations, following the geometric approach for the resonance Raman calculations, were carried out with the TDDFT formalism^47-50^ and were solved for a total of forty states for each combination of functional/basis set. For the resonance Raman frequency calculations, the Franck-Condon method was used.^51-54^ The path-integral approach was utilized with explicit definition of the transition dipole moments. 2^12^ steps were used for integration with the correlation function computed at each step and a time interval of 2^12^×10^-17^ sec. All resonance spectra were computed at the 0-0 transition between the reference (S_1_) and the resonant (r_n_) state. Test calculations employing the experimental incident light frequency produced no discernible differences. A HWHM homogeneous broadening of 20 cm^-1^ was applied to the spectra. For the evaluation of the calculated resonance Raman intensities, the dipole-dipole interactions on the XY plane were inspected at each vibration (lumiflavin is almost planar), as well as their corresponding Huang-Rhys factors^55^ – which give identical results. For the Raman spectra calculated at optimized structures on the S_1_ manifold (off-Resonance, offR) a HWHM value of 10 cm^-1^ was used to match the experimental line curves. For comparison with the calculated spectra, the experimental FSRS third Evolution Associated spectrum (EAS) was used with a life time of τ = 2.9 ns assigned to the equilibrated 1FMN* state.^16^

Simple statistical analysis was performed, due to the vastness of the data set, including the simple deviation σ = |V_T_-V_E_|, the percent error δand their averages μ_σ_ and μ_δ_. The percent error is given by the equation 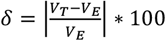. Since the experiments are used as a reference, V_T_ and V_E_ in the equations are defined as the theoretical and experimental values, respectively.

## Results and Discussion

The study proceeded with the optimization, excitation analysis for 40 states, and Raman frequency calculation of the ground state of all the selected DFT functionals (**Scheme 2**, steps 1 and 2). The two major excitations of lumiflavin (S_0_→S_1_ and S_0_→S_2_) were compared to the experimental values for FMN of 445 and 372 nm respectively.^17^ The average percent error **μ**_**δ**_of the two values for each DFT functional is included in the chart of **Figure 1**, sorted from larger to smallest error. B3LYP predicts excitations close to the experimental values -which justifies its popularity among DFT functionals in the study of flavin systems -^9, 28-29^ with 6.3% and 4.0% percent error for the cc-pVDZ and augmented equivalent, respectively. Long range corrected functionals such as LC-OPBE and CAM-B3LYP fare much worse, with over 20% error, similarly to the BHandHLYP functional (50% HF exchange), irrespective of the basis set. Among the most accurate predictions are given by the Minesota meta-GGA functionals M06L and M11L with errors lower than ∼3%. A similar graph to **Figure 1** is included in the **ESI** (**Figure S1**) that plots the average **μ**_σ_ and the separate deviation values **σ**_**S0→S1**_ and **σ**_**S0→S2**_ for the two excitations. All separate and averaged deviations and percent errors for the two excitations are tabulated in **Table S4**.

**Figure 1.**
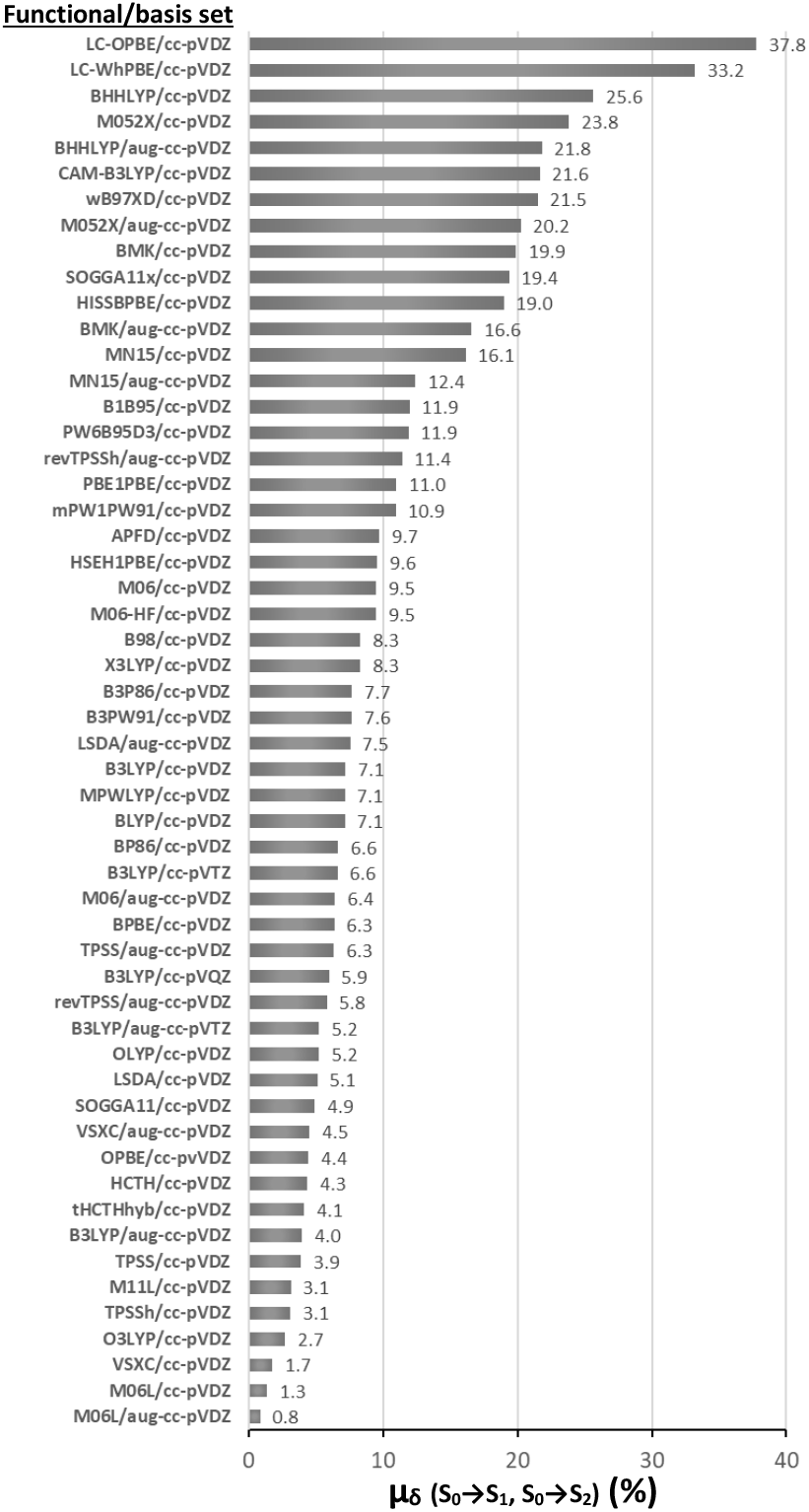
Average Percent error (**μ**_**δ**_, %) of the two major excitations of lumiflavin with each of the tested DFT functionals with respect to the experimental absorption values of FMN.

Subsequently, the first excited singlet state S_1_ was optimized (**Scheme 2**, Step 3). Most of the functionals predicted the first root (r_1_), while some of the GGA functionals, particularly at the smaller basis set, predicted r_2_ as the lowest singlet excited state. The optimized S_1_ state was identified as the lowest energy ππ* transition, before and after optimization (**Scheme 2**, Step 3.1). For LSDA/aug-cc-pVDZ in particular, the r_1_ state was identified as nπ*, thus further analysis was carried only on with the cc-pVDZ basis set (see **Table S7** in the **ESI** for hole/electron information on the S_1_ ππ* states). Then, the off-Resonance Raman spectra of all verified S_1_ states were computed and correlated with the experimental spectrum (**Scheme 2**, Step 3.2). The spectra were correlated with the 3^rd^ Evolution Associated Spectrum (EAS) of the FSRS spectrum of FMN, assigned as the equilibrated 1FMN* state (see **Table S9** in the **ESI**).^16^ Eight prominent peaks are featured in the fingerprint region of the EAS (**Figure 2**, blue line), and these were correlated to the calculated peaks in the 1000-1900 cm^-1^ region. Due to the abundance of calculated vibrations in the fingerprint region, multiple vibrations can be correlated to each experimental peak: for the FSRS EAS peak at 1200 cm^-1^ two computed peaks, mostly between v_48_-v_52_ were correlated, for 1250 cm^-1^ there was a single peak correlation, most frequently either v_51_ or v_53_, for 1338 cm^-1^ vibrations between v_54_-v_56_ were correlated, for 1381 cm^-1^ up to four computed peaks between v_57_-v_61_ were correlated, for 1416 cm^-1^ either v_64_ or v_65_, for 1498 cm^-1^ mostly v_71_ was correlated and secondarily v_70_ or v_72_, for 1570 cm^-1^ either v_73_ or v_74_ and finally for 1626 cm^-1^ v_75_, the last vibration in the fingerprint region was correlated. Typical vibrations with displacement vectors are shown in **Table S5** in the **ESI** taken from the B3LYP/aug-cc-pVDZ S_1_ frequency calculation.

**Figure 2.**
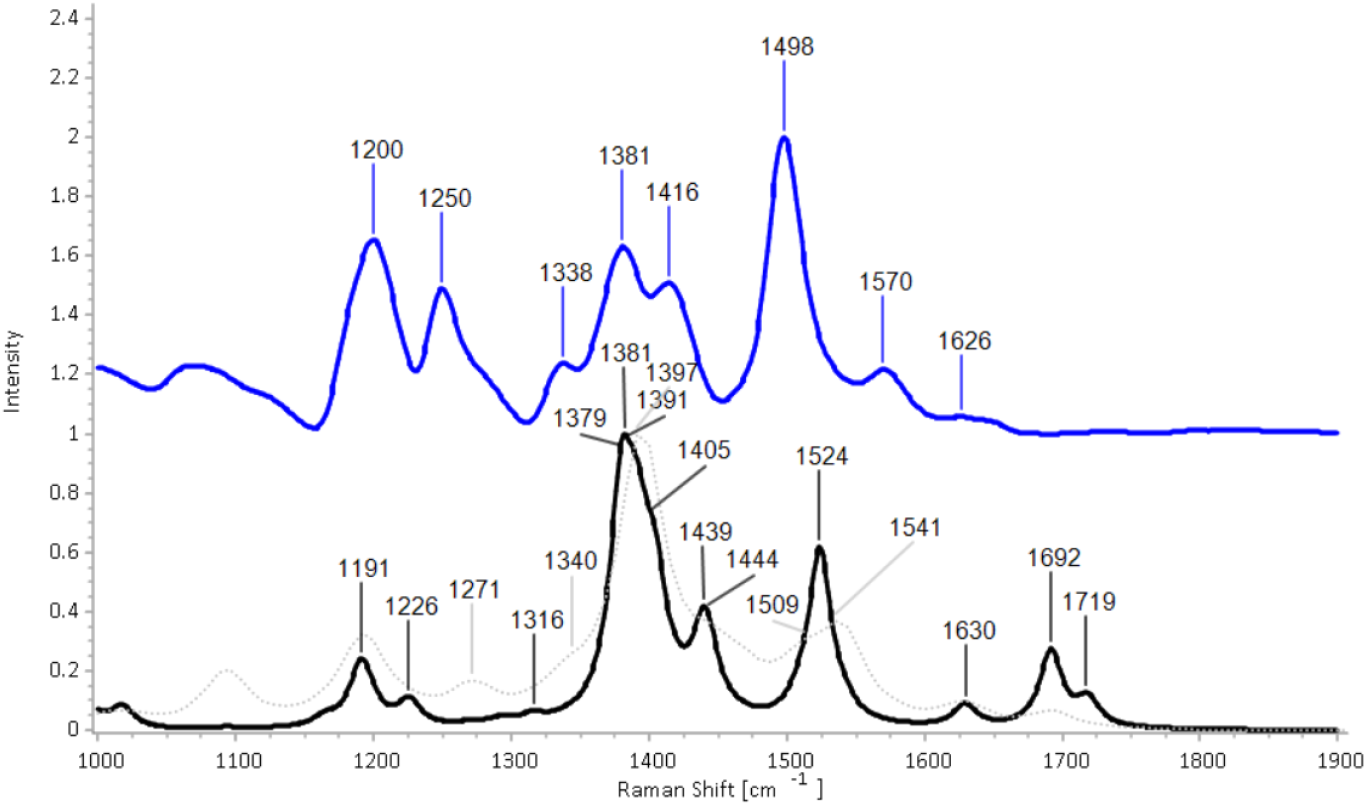
The Experimental FSRS 3^rd^ EAS of FMN assigned to the equilibrated 1FMN* state is shown as a solid blue line,^16^ the calculated off-Resonance S_1_ spectrum at the B3LYP/cc-pVDZ level is shown as a solid black line and the calculated resonance Raman spectrum of the S_1_->r_7_ transition at the at the same LOT is overlayed with a thin dotted grey line. All assigned peaks from **Table 1** have been labelled.

Following analysis and assignment of computed vibrations to normal modes (see **Table S9** in the **ESI**), the computed spectra at each functional was correlated with the eight experimental peaks, as described above. An example of such correlation is shown in **Figure 2** and the left portion of **Table 1** for the unscaled offR S_1_ spectrum computed with B3LYP/cc-pVDZ. Computed peaks at 1381 cm^-1^ (together with peaks 1379, 1391 and 1405 cm^-1^) and 1524 cm^-1^ are easily assigned to the experimental 1381 and 1498 cm^-1^ peaks, respectively. Averaging the deviations **σ** for all experimental/theoretical matches gives the overall agreement, which in this example is **μ**_**σ**_**(offR)** = 37.3 cm^-1^. The large value is mostly due to the large deviations of the assignments for the higher frequency peaks in the spectrum at 1570 and 1626 cm^-1^. The above process was repeated for all off-Resonance spectra from the DFT functionals and yielded **μ**_**σ**_**(offR)** values included in the 3^rd^ column of **Table 2**. Specifically, the very weak peak at 1626 cm^-1^, has been assigned to v_75_ in all correlations. All DFT functionals assign this as the symmetric C=O stretch of lumiflavin, which is expected to be more Raman active than the asymmetric equivalent.

**Table 1.**
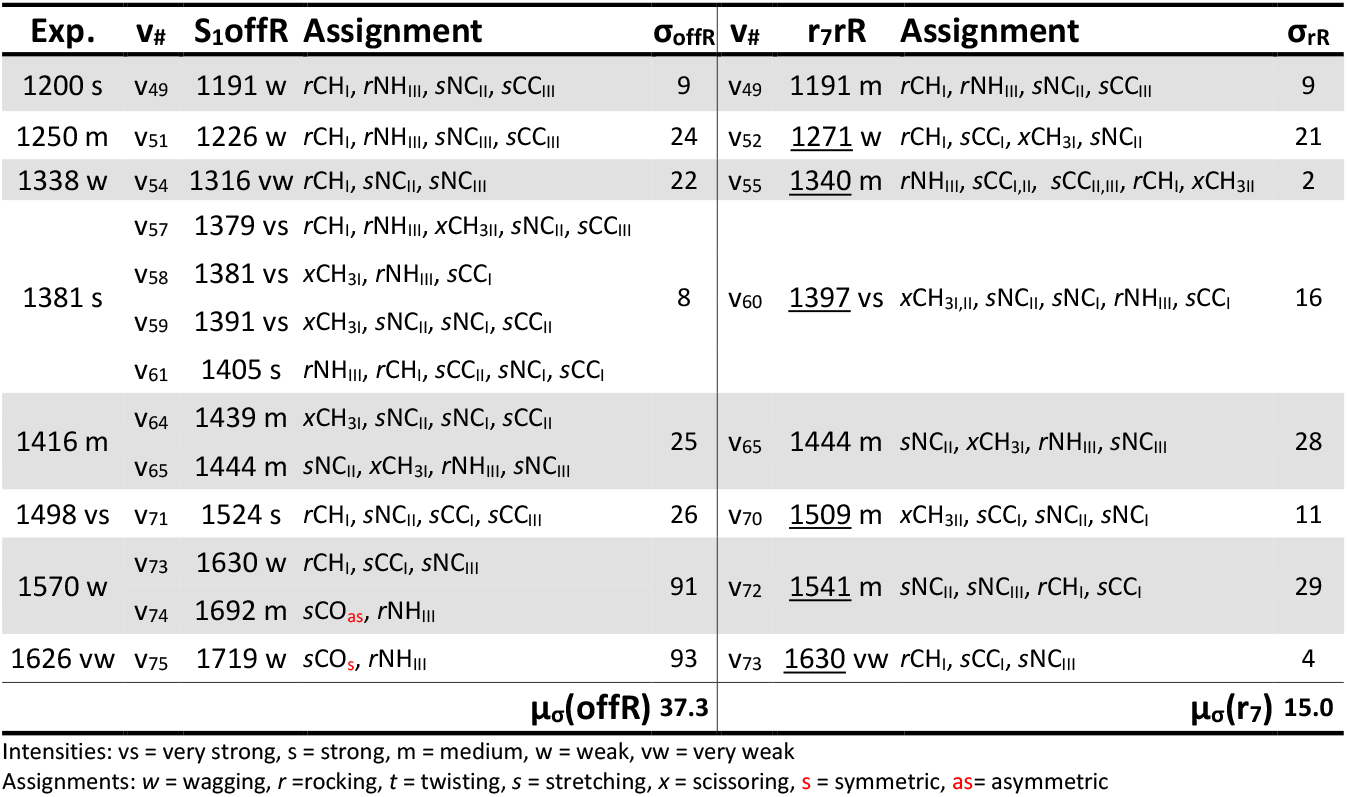
Correlation between the experimental FSRS 3^rd^ EAS of FMN (Exp.) and the off-Resonance S_1_offR and resonance r_7_rR Raman calculations of lumiflavin with B3LYP/cc-pVDZ. Peak intensities, vibration numbers and normal mode assignments are included. The latter are simplified using the Latin numerals I-III to assign the modes to each of the isoalloxazine rings (see **Scheme 1**). Deviation values from the experimental (**σ**_**offR**_, **σ**_**rR**_) are included along with their averages (**μ**_**σ**_**(offR), μ**_**σ**_**(r**_**7**_)) in the last row. New assigned peaks due to the inclusion of resonance in the calculations are underlined.

**Table 2.** Values of the statistical analysis pertaining to the agreement between the experimental spectrum third EAS of 1FMN* and the lumiflavin calculated spectra of S_1_ (offR) and Resonance Raman r_n_. The terms **μ**_**σ**_**(offR)** and **μ**_**σ**_**(r**_**n**_) are the average deviations of the offR and rR spectra, respectively and **Δμ**_**σ**_ their difference. The term **μ**_**σ**_**(Sc**_**L**_) gives the average deviation after applying the literature (or computed with FREQ for this study) scaling factor, and **μ**_**σ**_**(Sc**_**S**_) is the average deviation after applying the specific scaling factor included in **Table S3**. The term **μ**_**σ**_**(Sc**_**S**_**)\*** involves all DFT functionals with a specific scaling factor lower than 0.93, that re-assignment of peaks was performed after scaling. The terms **μ**_**δ**_**(offR)** and **μ**_**δ**_**(r**_**n**_) are the average percent errors of the offR and rR spectra, respectively and **Δμ**_**δ**_their difference. All deviation values (σ) are given in cm^-1^ and percent errors (δ) in %.

| LOT | State | $\mu_\sigma(\text{offR})$ | $\mu_\sigma(r_n)$ | $\Delta\mu_\sigma$ | $\mu_\sigma(\text{Sc}_L)$ | $\mu_\sigma(\text{Sc}_S)$ | $\mu_\sigma(\text{Sc}_S)^*$ | $\mu_\delta(\text{offR})$ | $\mu_\delta(r_n)$ | $\Delta\mu_\delta$ |
| --- | --- | --- | --- | --- | --- | --- | --- | --- | --- | --- |
| APFD/cc-pVDZ | $r_5$ | 47.1 | 51.3 | -4.2 | 39.8 | 53.5 | - | 3.11 | 3.41 | -0.30 |
| B1B95/cc-pVDZ | $r_7$ | 68.5 | 36.9 | 31.7 | 31.6 | 46.6 | 44.3 | 4.75 | 2.53 | 2.22 |
| B3LYP/cc-pVDZ | $r_7$ | 37.3 | 15.0 | 22.3 | 40.0 | 54.1 | - | 2.50 | 1.07 | 1.42 |
| B3LYP/aug-cc-pVDZ | $r_6$ | 23.6 | 17.4 | 6.2 | 34.8 | 27.1 | - | 1.61 | 1.23 | 0.38 |
| B3LYP/cc-pVTZ | $r_7$ | 29.3 | 23.8 | 5.5 | 29.8 | 31.1 | - | 1.99 | 1.63 | 0.35 |
| B3LYP/aug-cc-pVTZ | $r_7$ | 23.5 | 22.3 | 1.3 | 28.7 | 20.9 | - | 1.61 | 1.54 | 0.07 |
| B3LYP/cc-pVQZ | $r_6$ | 27.2 | 25.3 | 1.8 | 27.7 | 26.1 | - | 1.85 | 1.76 | 0.09 |
| B3P86/cc-pVDZ | $r_7$ | 56.8 | 47.1 | 9.7 | 27.5 | 40.9 | - | 3.93 | 3.19 | 0.73 |
| B3PW91/cc-pVDZ | $r_7$ | 38.1 | 20.9 | 17.2 | 39.2 | 57.0 | - | 2.46 | 1.47 | 0.99 |
| B98/cc-pVDZ | $r_7$ | 44.3 | 32.1 | 12.1 | 31.7 | 50.9 | - | 3.02 | 2.15 | 0.87 |
| BHHLYP/cc-pVDZ | $r_4$ | 96.1 | 50.3 | 45.9 | 59.0 | 79.7 | 28.5 | 6.48 | 3.45 | 3.03 |
| BHHLYP/aug-cc-pVDZ | $r_5$ | 83.0 | 48.2 | 34.9 | 48.6 | 51.0 | 16.9 | 5.65 | 3.48 | 2.17 |
| BLYP/cc-pVDZ | $r_7$ | 27.2 | 39.8 | -12.6 | 31.5 | 63.9 | - | 1.96 | 2.92 | -0.97 |
| BMK/cc-pVDZ | - | 61.4 | - | - | 49.5 | 91.6 | 24.4 | 4.06 | - | - |
| BMK/aug-cc-pVDZ | $r_5$ | 47.7 | 32.4 | 15.3 | 36.7 | 57.6 | 40.0 | 3.18 | 2.34 | 0.84 |
| BP86/cc-pVDZ | $r_7$ | 27.8 | 22.2 | 5.6 | 27.5 | 48.3 | - | 2.02 | 1.63 | 0.40 |
| BPBE/cc-pVDZ | $r_7$ | 22.3 | 17.8 | 4.6 | 30.9 | 44.8 | - | 1.59 | 1.30 | 0.30 |
| CAM-B3LYP/cc-pVDZ | $r_5$ | 60.2 | 30.9 | 29.3 | 47.9 | 68.3 | 42.3 | 3.95 | 2.21 | 1.73 |
| HCTH/407/cc-pVDZ | $r_7$ | 30.6 | 16.7 | 13.9 | 29.8 | 49.0 | - | 2.05 | 1.19 | 0.86 |
| HISSbPBE/cc-pVDZ | $r_7$ | 105.6 | 60.5 | 45.0 | 34.9 | 56.2 | 42.6 | 7.39 | 4.16 | 3.24 |
| HSEH1PBE/cc-pVDZ | $r_7$ | 51.2 | 22.4 | 28.8 | 34.9 | 56.2 | 41.1 | 3.39 | 1.59 | 1.80 |
| LC-OPBE/cc-pVDZ | $r_4$ | 144.9 | 233.3 | -88.5 | 64.9 | 80.0 | 38.5 | 9.81 | 16.46 | -6.64 |
| LC-wHPBE/cc-pVDZ | $r_5$ | 154.0 | 188.8 | -34.8 | 62.9 | 28.6 | 20.9 | 11.05 | 13.45 | -2.40 |
| LSDA/cc-pVDZ | $r_8$ | 36.4 | 27.1 | 9.3 | 31.5 | 63.9 | - | 2.48 | 2.02 | 0.46 |
| LSDA/aug-cc-pVDZ | - | 39.0 | - | - | 30.3 | 36.3 | - | 2.77 | - | - |
| M05-2X/aug-cc-pVDZ | $r_4$ | 56.1 | 40.5 | 15.6 | 33.6 | 36.1 | - | 3.78 | 2.71 | 1.08 |
| M06/cc-pVDZ | $r_7$ | 51.3 | 43.2 | 8.1 | 50.2 | 81.1 | 24.8 | 3.39 | 2.85 | 0.54 |
| M06/aug-cc-pVDZ | $r_5$ | 35.8 | 57.2 | -21.4 | 37.4 | 49.7 | - | 2.36 | 4.04 | -1.67 |
| M06-HF/cc-pVDZ | $r_6$ | 42.0 | 18.6 | 23.4 | 58.7 | 84.4 | 26.8 | 2.72 | 1.31 | 1.40 |
| M06L/cc-pVDZ | $r_5$ | 60.4 | 45.7 | 14.6 | 19.9 | 52.5 | 17.1 | 4.21 | 3.14 | 1.06 |
| M06L/aug-cc-pVDZ | $r_7$ | 32.1 | 25.6 | 6.5 | 37.8 | 40.0 | - | 2.17 | 1.75 | 0.43 |
| M11L/cc-pVDZ | $r_6$ | 56.3 | 22.6 | 33.7 | 50.4 | 85.7 | 22.9 | 3.71 | 1.60 | 2.12 |
| MN15/cc-pVDZ | $r_5$ | 52.2 | 53.0 | -0.7 | 49.3 | 73.7 | 16.0 | 3.41 | 3.58 | -0.18 |
| MN15/aug-cc-pVDZ | $r_4$ | 45.2 | 25.1 | 20.1 | 37.4 | 44.7 | - | 3.04 | 1.85 | 1.19 |
| mPW1PW91/cc-pVDZ | $r_7$ | 56.8 | 16.2 | 40.6 | 33.4 | 53.5 | 42.1 | 3.79 | 1.14 | 2.65 |
| mPWLYP/cc-pVDZ | $r_7$ | 20.6 | 31.5 | -10.9 | 22.9 | 27.2 | - | 1.56 | 2.32 | -0.76 |
| O3LYP/cc-pVDZ | $r_7$ | 40.6 | 14.8 | 25.8 | 45.2 | 63.1 | - | 2.75 | 1.05 | 1.70 |
| OLYP/cc-pVDZ | r<sub>7</sub> | 36.1 | 14.4 | 21.7 | 36.2 | 58.8 | - | 2.56 | 1.04 | 1.52 |
| OPBE/cc-pVDZ | r<sub>7</sub> | 39.9 | 22.3 | 17.6 | 45.0 | 67.1 | - | 2.72 | 1.63 | 1.10 |
| PBE1PBE/cc-pVDZ | r<sub>5</sub> | 49.6 | 43.6 | 6.1 | 38.7 | 61.0 | 43.8 | 3.26 | 2.89 | 0.36 |
| PW6B95D3/cc-pVDZ | r<sub>7</sub> | 54.4 | 65.9 | -11.5 | 40.3 | 55.3 | 41.4 | 3.60 | 4.40 | -0.80 |
| revTPSS/aug-cc-pVDZ | r<sub>7</sub> | 7.0 | 15.3 | -8.3 | 25.2 | 6.4 | - | 0.50 | 1.13 | -0.63 |
| revTPSSh/aug-cc-pVDZ | r<sub>7</sub> | 162.3 | 51.3 | 110.9 | 42.6 | 12.7 | 20.6 | 11.49 | 11.39 | 0.10 |
| SOGGA11/cc-pVDZ | r<sub>9</sub> | 25.8 | 24.2 | 1.6 | 43.8 | 65.1 | - | 1.79 | 1.63 | 0.16 |
| SOGGA11x/cc-pVDZ | r<sub>5</sub> | 73.6 | 54.6 | 18.9 | 43.8 | 65.1 | 40.1 | 4.95 | 3.82 | 1.13 |
| tHCTHhyb/cc-pVDZ | r<sub>7</sub> | 34.9 | 22.1 | 12.8 | 42.0 | 56.8 | - | 2.35 | 1.48 | 0.87 |
| TPSS/cc-pVDZ | r<sub>5</sub> | 27.3 | 21.3 | 5.9 | 18.5 | 34.2 | - | 1.90 | 1.46 | 0.44 |
| TPSS/aug-cc-pVDZ | r<sub>7</sub> | 7.0 | 15.5 | -8.6 | 29.6 | 6.6 | - | 0.51 | 1.13 | -0.62 |
| TPSSh/cc-pVDZ | r<sub>7</sub> | 35.7 | 14.8 | 20.9 | 31.1 | 44.0 | - | 2.44 | 1.04 | 1.40 |
| VSXC/cc-pVDZ | r<sub>5</sub> | 40.3 | 33.1 | 7.2 | 22.5 | 43.9 | - | 2.79 | 2.28 | 0.52 |
| VSXC/aug-cc-pVDZ | r<sub>7</sub> | 16.0 | 21.3 | -5.4 | 27.1 | 13.1 | - | 1.14 | 1.51 | -0.37 |
| wB97XD/cc-pVDZ | r<sub>6</sub> | 71.4 | 43.3 | 28.0 | 35.4 | 58.5 | 36.8 | 4.84 | 2.96 | 1.89 |
| X3LYP/cc-pVDZ | r<sub>7</sub> | 44.6 | 43.8 | 0.9 | 29.3 | 40.8 | - | 3.05 | 2.90 | 0.16 |

\*: re-assignment of peaks for functionals with $Sc_s$ lower than 0.93.

The above considerations highlight a problem with all Raman frequency computations and the predicted frequency values. In **Figure 3**, the relationship between the C_2_=O_2’_ and C_4_=O_4’_ bond lengths and their respective symmetric and asymmetric stretching frequencies is plotted. It can be seen that most DFT functionals predict both vibrations to lie between 1800-1700 cm^-1^, and some long range corrected functionals even near 1900 cm^-1^. The experimental value highlighted in yellow, is centred at 1626 cm^-1^ showing the discrepancy. On the other hand, some functionals cluster in the lower range >1700 cm^-1^ including: TPSS and its revision, BLYP, mPWLYP, VSXC, B3LYP, BPBE, M06, OLYP and SOGGA11 mentioned in the order from closest furthest from the experimental value. A structural solution that has been applied widely is microsolvation,^17, 28^ where explicit solvation is combined with the implicit solvent model. Water molecules are placed at key H-bonding positions, including proximity to the two flavin C=O bonds. The resulting bond lengthening, concomitant with the shift of the vibrations to the red, would skew their distribution to the left part of the graph of **Figure 3**, closer to the experimental value.

**Figure 3.**
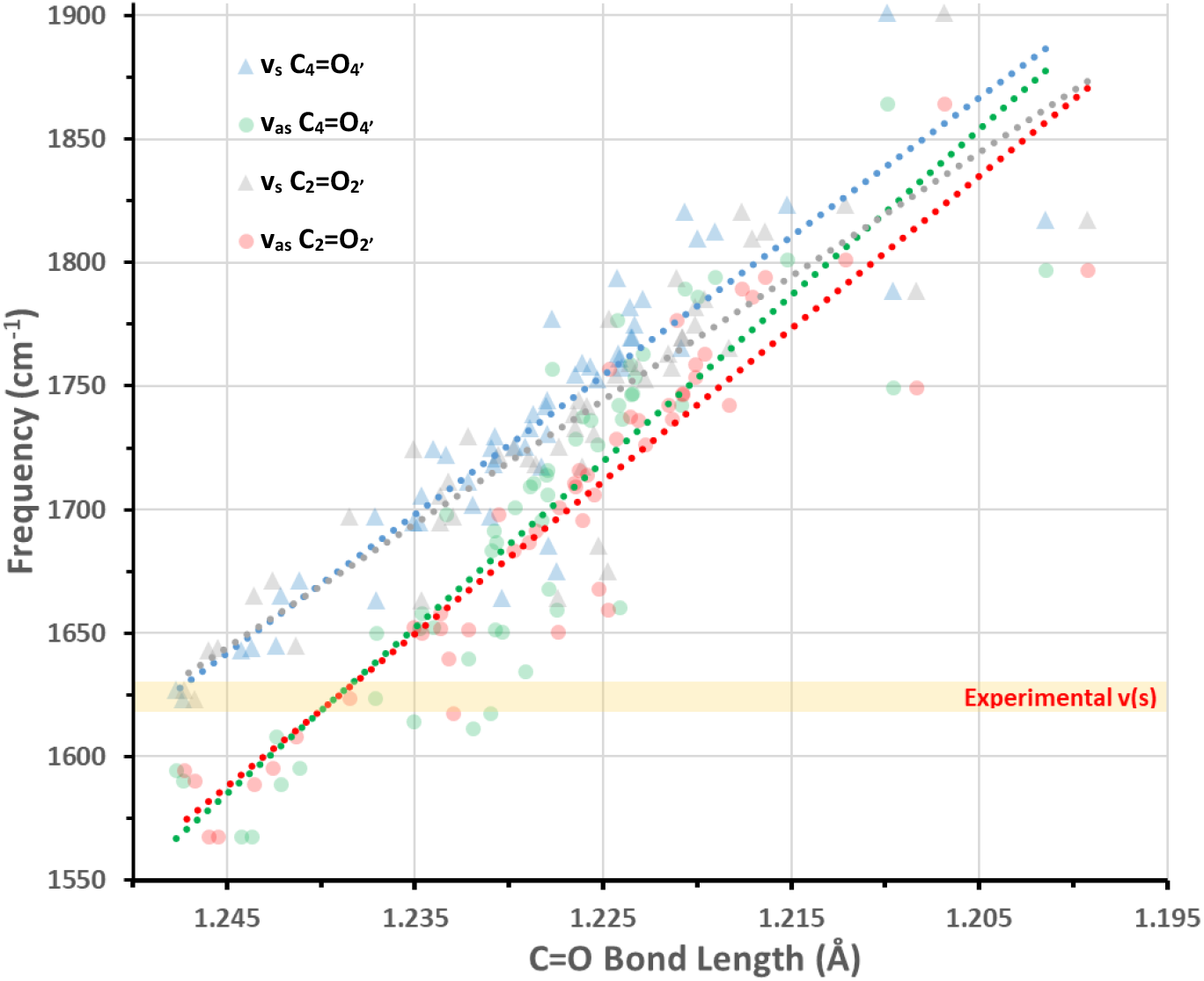
Relationship between the length of the C_2_=O_2’_ and C_4_=O_4’_ bonds (x-axis) calculated with the tested DFT functionals, and their symmetric (s) and asymmetric (as) stretching frequencies calculated with the same DFT functionals (y-axis). The v_s_/ C_2_=O_2’_ and C_4_=O_4’_ bond length values are given in grey and blue triangles respectively, while the v_as_ C_2_=O_2’_ and C_4_=O_4’_ in red and green squares, respectively. Trendlines with corresponding colour have been drawn for each of the four sets. The experimental C_2_=O_2’_/C_4_=O_4’_ symmetric stretching v(s) frequency (1626 cm^-1^) has been highlighted in yellow.

For the context of this benchmark, the discrepancy was addressed by scaling the frequencies uniformly, which was undertaken either by applying literature scaling factors and alternatively the ones computed in this study by FREQ (**Sc**_**L**_), or by devising a specific scaling factor to align v_75_ to 1626 cm^-1^ (**Sc**_**S**_) (**Scheme 2**, Step 3.3). All scaling factors that were employed are included in the **ESI** in **Table S3**. The **Sc**_**L**_ and **Sc**_**S**_ factors were applied uniformly to the unscaled peaks in each assignment table, yielding new **μ**_**σ**_**(Sc**_**L**_) and **μ**_**σ**_**(Sc**_**S**_) values, included in **Table 2** columns 6 and 7 respectively. The application of the scaling factors gave a mixed picture. It improved the worse correlations including the long range corrected LC-wHPBE from 154.0 to 62.9 and 28.6 cm^-1^ for **μ**_**σ**_**(Sc**_**L**_) and **μ**_**σ**_**(Sc**_**S**_), respectively. Conversely, for functionals with reasonable unscaled agreement, such as SOGGA11 (**μ**_**σ**_**(offR)** = 25.8 cm^-1^), the correlation was lowered to 43.8 and 65.1 cm^-1^ for **μ**_**σ**_**(Sc**_**L**_) and **μ**_**σ**_**(Sc**_**S**_), respectively. Specifically for DFT functionals that their **Sc**_**S**_ factor was found below 0.93 (see **Table S3** in the **ESI**, 21 functionals) a re-assignment of peaks was performed, with new corresponding values **μ**_**σ**_**(Sc**_**S**_**)\*** (8^th^ column, **Table 2**). The re-assignment improved the correlations of all 21 functionals compared to **μ**_**σ**_**(Sc**_**L**_) and **μ**_**σ**_**(Sc**_**S**_) values (i.e. the LC-wHPBE **μ**_**σ**_**(Sc**_**S**_**)\*** value was lowered to 20.9 cm^-1^). Thus, scaling factors can be a useful tool, specifically in the absence of anharmonic corrections, but care should be taken in their usage, not with blanket application to the unscaled values. It should be mentioned here that only a few of the DFT functionals gave better agreement either using the **μ**_**σ**_**(S**_**1**_**), μ**_**σ**_**(Sc**_**L**_) or **μ**_**σ**_**(Sc**_**S**_) values, than the full FMN structure calculation with B3LYP/def2-TZVP (**μ**_**σ**_**(Sc**_**L**_**) =** 19 cm^-1^).^17^

The next step in the in the computational regime is the choice of excitations within the experimental resonant window, defined as the wavelength of the Raman pump 800±100 nm (**Scheme 2**, Step 4). The window was in practice extended to more than ±300 nm to accommodate as many states as possible in the study. Transition dipole moments, excitation energies and oscillator strengths between S_1_ and the higher singlet states r_n_ were determined using the program Multiwfn 3.8,^56^ and are collected in **Table S6** in the **ESI**, along with the difference between pump and excitation energy (**Δr**). From all the states chosen, the most suitable candidates are shown in bold, owing to the high Oscillator strengths and proximity to the resonance wavelength. All states included in **Table S6** were optimized (**Scheme 2**, Step 5), either leading to the expected state or regressing to other state potential energy surfaces (PES), including S_1_. This was verified from their respective energies, and additionally from the hole-electron properties and distribution surfaces included in **Tables S7** and **S8** in the **ESI**, respectively). For example, optimization of the r_5_ state with BMK/cc-pVDZ yielded the S_1_ state and no further analysis could be completed for this level of theory (LOT). Thus, the basis set was increased to aug-cc-pVDZ and steps 1-4 were repeated. Similarly, states r_5_-r_8_ yielded the same r_7_ state after optimization with M06/cc-pVDZ, and accordingly the study proceeded only with r_7_ for that LOT.

Subsequently, the excited Raman spectra of all unique states for each LOT were computed. This led to the final step (**Scheme 2**, Step 6), the Resonance Raman calculation using the r_n_ as the resonant state and the corresponding S_1_ state as the reference. Spectra were obtained at the 0-0 transition between reference and resonant state and a broadening of 20 cm^-1^ was applied to the peaks. Correlation and re-assignment of spectra was performed with the FSRS 3^rd^ EAS with the off-Resonance assignment as a basis (**Scheme 2**, Step 6.1) using only unscaled frequencies. In the B3LYP case, the new assignment is shown in right portion of **Table 1**. In that case, six out of eight assignments are new due to resonance enhancement, and this brings a superior corelation **μ**_**σ**_**(r**_**7**_) = 15 cm^-1^ than the off-Resonance. The advantage of the rR is that single peaks that are resonance enhanced are assigned to the experimental ones in contrast to the averaging of peaks that is needed in the off-Resonance correlation. However, as was mentioned in the Introduction, the calculated resonance Raman spectrum of B3LYP produces worse relative intensities than the off-Resonance with respect to the experimental FSRS - which is in resonant conditions itself. This is evident for the calculated peak at 1541 cm^-1^ of the r_7_ spectrum included in **Figure 1** as thin grey dotted line.

The same process of re-assignment and correlation of the rR spectra was repeated for the other functionals producing **μ**_**σ**_**(r**_**n**_) and **μ**_**δ**_**(r**_**n**_) averages for the correlation. To specifically address the issue mentioned above and give a measure of the improvement or worsening of the correlation with the inclusion of resonance in the calculations the **Δμ**_**δ**_(r_n_-offR) term was calculated. This is a simple subtraction of the **μ**_**δ**_**(offR)** and **μ**_**δ**_**(r**_**n**_) values returning a positive number if the correlation improves and negative if it worsens with the inclusion of resonance in the spectra of each combination of DFT functional/basis set. The **μ**_**δ**_**(r**_**n**_) and **Δμ**_**δ**_terms are included together in **Figure 4** from higher to lower **μ**_**δ**_**(r**_**n**_) values. All mentioned terms are included in the final three columns of **Table 2** (**μ**_**δ**_**(offR), μ**_**δ**_**(r**_**n**_**), Δμ**_**δ**_). For most functionals the correlation was improved with the recomputed intensities, with the exception of ten functional/basis set combinations: APFD/cc-pVDZ (-0.30%), M06/aug-cc-pVDZ (-1.67%), BLYP/cc-pVDZ (-0.97%), mPWLYP/cc-pVDZ (-0.76%), PW6B95D3/cc-pVDZ (-0.80%), revTPSS/aug-cc-pVDZ (-0.63%), TPSS/aug-cc-pVDZ (-0.62%) and VSXC/aug-cc-pVDZ (-0.37%). The largest differences were found for the range-corrected functionals LC-OPBE/cc-pVDZ (- 6.64 %) and LC-wHPBE/cc-pVDZ (-2.40%).

**Figure 4.**
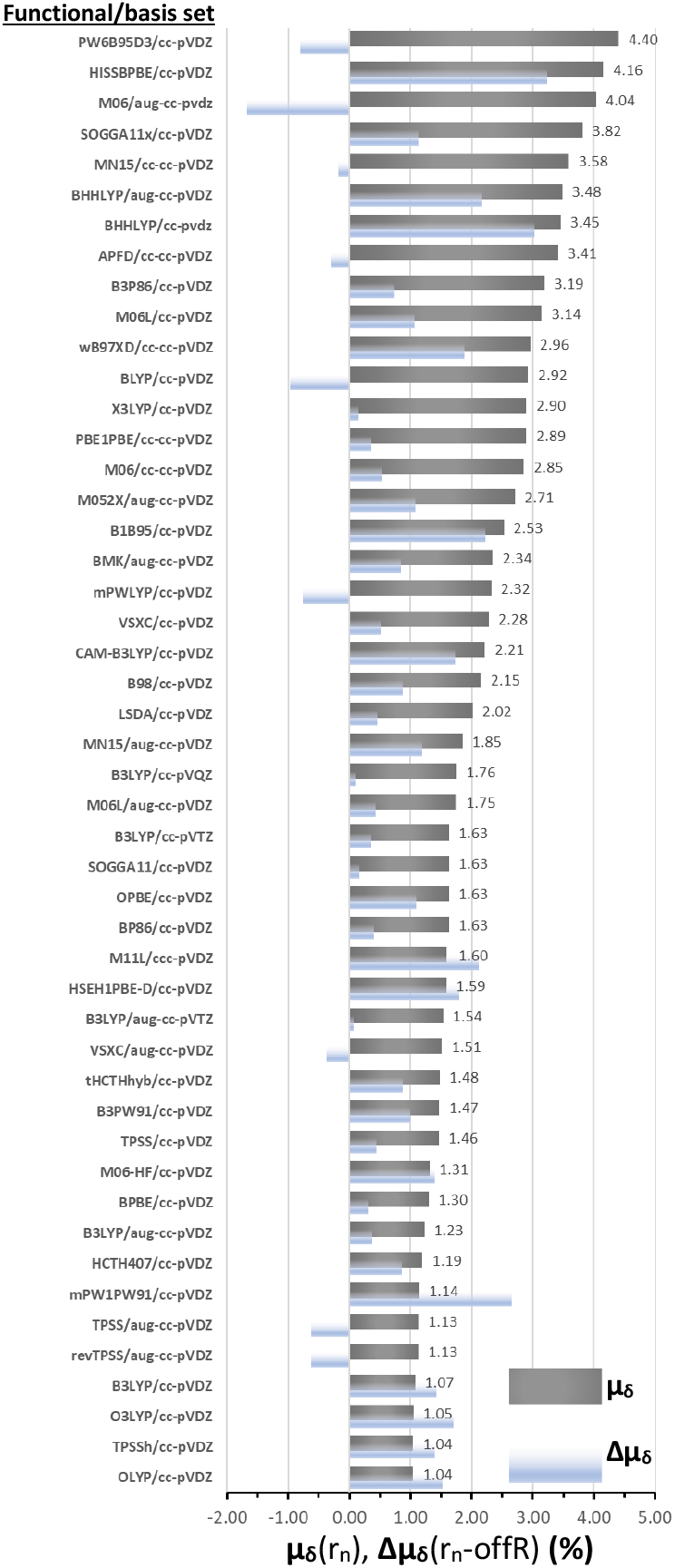
Average Percent error (**μ**_**δ**_, %) of the major peaks of lumiflavin of each of the tested DFT functionals with respect to the experimental FSRS peak values of FMN. The **μ**_**δ**_values are based on the resonance Raman calculations (r_n_) while **Δμ**_**δ**_gives the subtraction between **μ**_**δ**_**(r**_**n**_) and **μ**_**δ**_**(offR)**. The μ_δ_values have been labelled.

Due to the large amount data considered, evaluation of all functionals according to specific criteria was required in order to extract useful information. This was attempted according to the five criteria formulated below. Each functional can obtain up to five positive marks (★) according to its compliance to the criteria thresholds. For cases that lie outside, but close to the thresholds, a half-mark was given. A positive evaluation is given for the DFT functionals that:

a. The mean percent error (**μ**_**δ**_) of their Vertical Excitations S_0_→S_1_ and S_0_→S_2_ lies below 5.1% error, and half mark below 7.1%. The equivalent values in mean deviations (**μ**_**σ**_) would be 24 and 33 nm, respectively.
b. The mean percent error, **μ**_**δ**_**(r**_**n**_) of the rR computed peaks associated with the eight most prominent experimental FSRS peaks of the 3^rd^ EAS is equal or lower than 1.5%. Half-mark is given for DFT functionals between 1.5-2% error. The equivalent mean deviation **μ**_**σ**_**(r**_**n**_) range would be 21-27 cm^-1^. For the functionals that more states were found within the resonant window, the **μ**_**δ**_**(r**_**n**_) values pertain only to the state **r**_**n**_ with the higher oscillator strength for the S_1_→r_n_ transition.
c. The difference between the off-Resonance and resonance Raman mean percent errors (**Δμ**_**δ**_) is positive. A positive value signifies an improvement of the correlation after the inclusion of resonant effects to the computed intensities and *vice versa*. If **Δμ**_**δ**_is negative, no mark is awarded, if it is higher than 1% the full mark is awarded and a half mark is given if for functionals with positive values between 0.3-1%.
d. For this criterion, the focus is on the most prominent experimental peak at 1498 cm^-1^. Instead of peak intensities, the dipole-dipole interactions between reference and resonant state on the X-Y plane are utilized (lumiflavin is almost planar). Alternatively, the Huang-Rhys terms gave similar results. The dipole-dipole values of the eight vibrations in the correlation with the experimental spectrum were normalized in the range of 0-1. Then, the normalized value of the vibration assigned to 1498 cm^-1^ (v_71_ for the majority of functionals) is evaluated according to the following criteria: For values between 0.8-1 a full mark is given, signifying that the DFT functional correctly (or almost correctly) predicts the strongest peak in the spectrum. A half mark is given for normalized values between 0.6-0.8, and below 0.5, no mark is awarded.
e. Finally, a subjective criterion is introduced, that of a visual inspection of the intensities of the computed resonance Raman spectra and their agreement with the experimental FSRS intensities. The computed spectra fall within five categories as follows: The DFT functional is evaluated positively when the correlation of the theoretical-experimental spectra is facile: (i) with a full mark for very similar line shapes to the experimental curve or (ii) half mark for less similar but still providing for a facile correlation. A negative evaluation with no mark is given for computed spectra that either: (iii) bear no visible doublet peaks that can be easily correlated to the 1200-1250 cm^-1^ and 1381-1416 cm^-1^ experimental pairs, or (iv) require a scaling factor for the frequencies, usually when v_73_ is predicted with strong intensity and scaling would align it with the experimental peak at 1498 cm^-1^ or finally, (v) the C=O symmetric or asymmetric stretch is predicted as the strongest peak in the spectrum.

The evaluation of the functionals is given in **Table 3** along with the total positive marks (out of 5) for each functional for the a)-e) criteria.

**Table 3.**
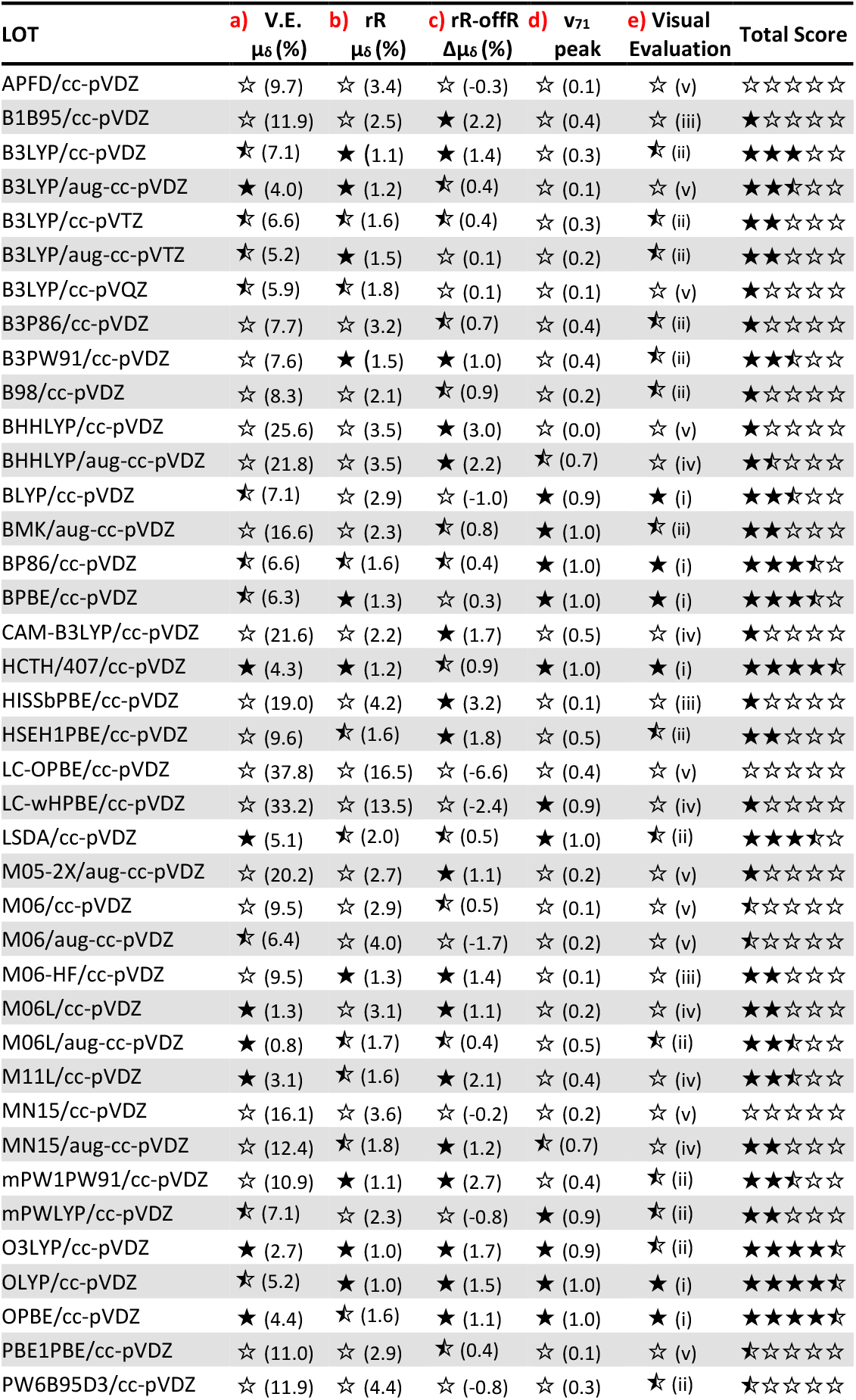
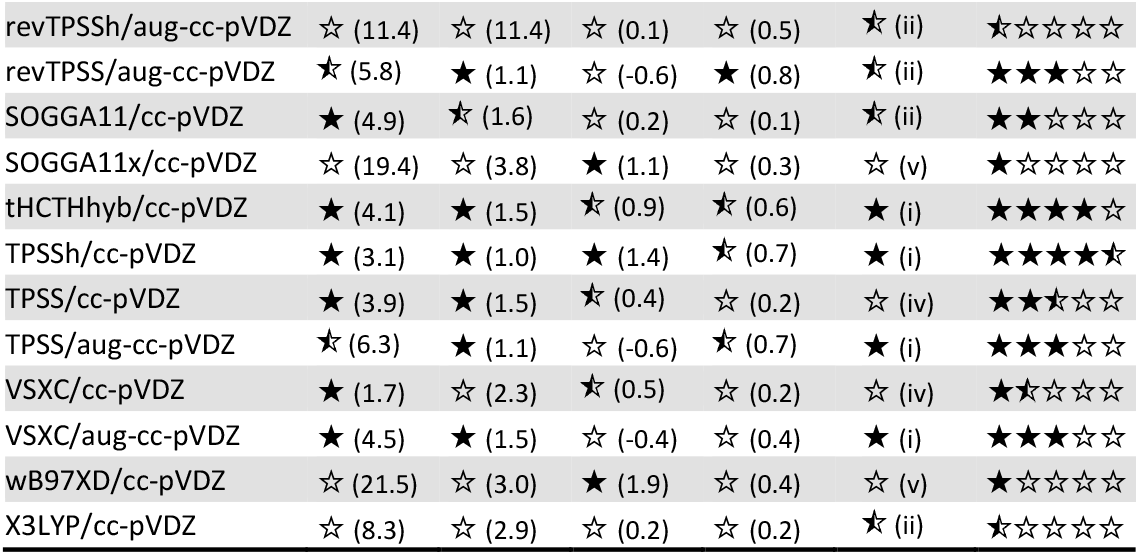
Evaluation of DFT Functionals according to the five criteria: a) the vertical excitation mean percent error, b) the resonance Raman mean percent error, c) the difference in the mean percent errors of the resonance and off-Resonance Raman spectra, d) the intensity of the v_71_ vibration assessed by normalized X-Y dipole-dipole values and e) visual evaluation, where the computed resonance Raman spectra are classified after visual inspection according to the categories (i)-(v) described in the main text.

Six DFT functionals stand out in their performance against the five criteria: the GGA functionals HCTH/407, OLYP and OPBE, the hybrid O3LYP functional with 11.6% HF-exchange and the meta-Hybrid functionals tHCTHhyb and TPSSh with 10 and 16% HF-exchange, respectively (all with the cc-pVDZ basis set). Surprisingly, the revised TPSSh functional fared much worse in the evaluation. The resonance Raman spectra of the six functionals are included in **Figure 5**. As can be seen in **Table 3**, for three of the six functionals (HCTH, OLYP, OPBE) v_71_ is the strongest peak in the fingerprint region, as per the experiment-for O3LYP the second strongest (0.9). Incidentally, by inspection of spectral intensities in **Figure 5**, peaks in the region ∼1350-1450 cm^-1^ appear more intense, however this is due to the concentration of quite a few medium-to-strong peaks in that region. On the whole, all six functionals enabled a facile correlation with the experimental EAS, with clear spectral features close to the experimental peaks marked with grey bars in **Figure 5**.

**Figure 5.**
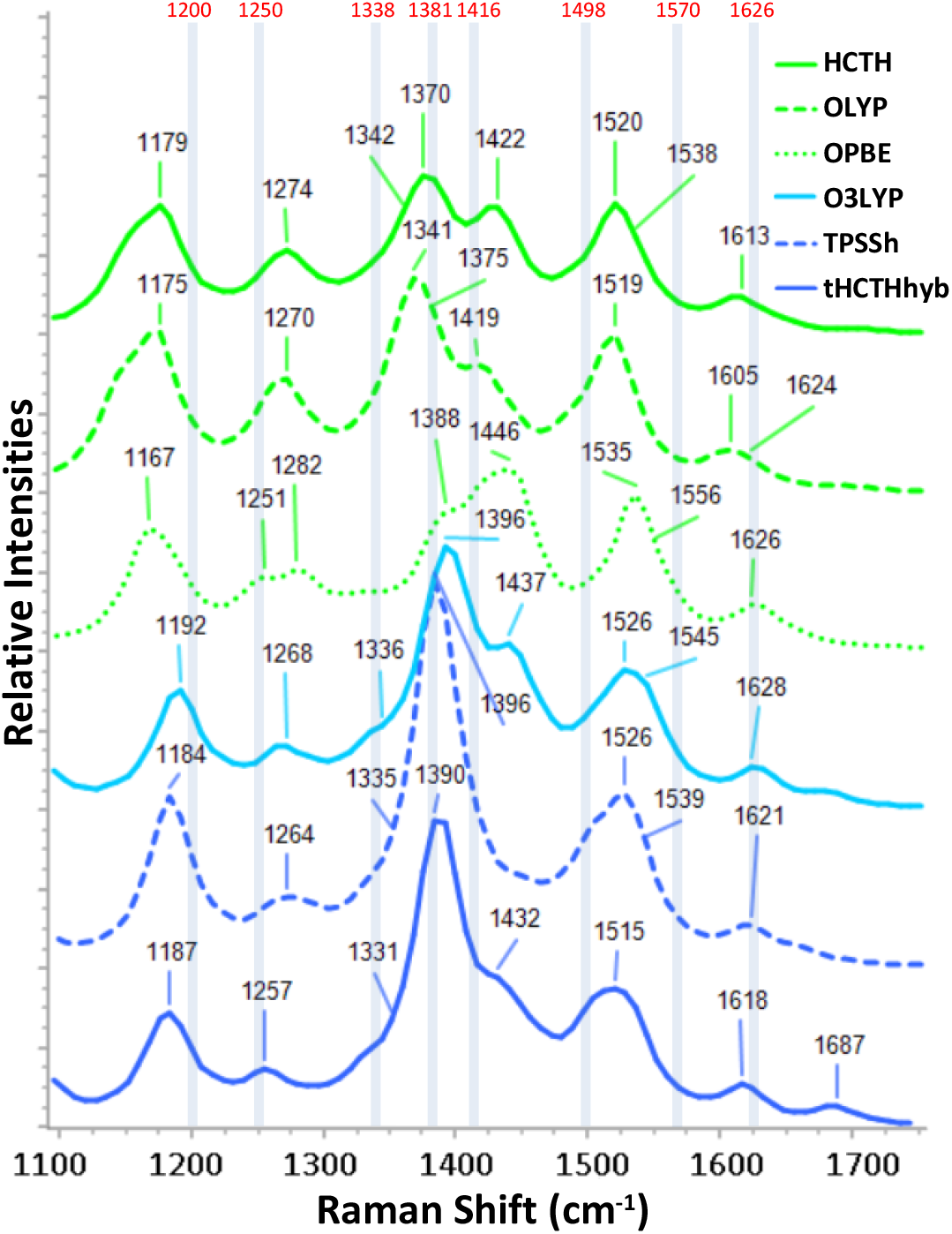
Calculated resonance Raman spectra of the best six performing DFT functionals of the benchmark with the cc-pVDZ basis set. The experimental 3^rd^ EAS of 1FMN* has been labelled and indicated with blue bars.^16^

For the basis set dependence with the B3LYP functional, there is clear improvement in the off-Resonance **μ**_**σ**_**(offR)**/**μ**_**δ**_**(offR)** values with the increase of the basis set which is more pronounced with the inclusion of diffuse functions (see **Table 2** and **μ**_**δ**_values in **Figure 4**). With regards to the rR **μ**_**σ**_(**r**_**n**_) values, marked improvement is achieved when compared to the corresponding **μ**_**σ**_**(offR)** values. Conversely, when comparing the **μ**_**σ**_**(r**_**n**_) values themselves with increasing basis set size, the opposite trend is evidenced, with slight increase in disagreement with every increase in basis set from 15 cm^-1^ with cc-pVDZ to 25.3 cm^-1^ with cc-pVQZ. Scaling with the literature scaling factors **Sc**_**L**_, does not improve any of the offR correlations. The specific ones (**Sc**_**S**_) improve only the higher in the series aug-cc-pVTZ and cc-pVQZ from 23.5 and 27.2 cm^-1^ to 20.9 and 26.1 cm^-1^, respectively. Overall, cc-pVDZ and aug-cc-pVDZ perform better with B3LYP in the five criteria than with the larger basis sets, as can be surmised from their total score in **Table 3**.

The six functionals HCTH/407, OLYP, OPBE, O3LYP, tHCTHhyb and TPSSh can be recommended for resonance Raman studies on the flavin family of chromophores. Yet, the collective knowledge on excited state calculations warns against the usage of the first three, pure GGA functionals, since they tend to underestimate Charge Transfer over Local Excitation state energies.^57^ While this did not pose a significant problem with standalone lumiflavin, in larger systems, numerous CT states real or artificial, would be present, for example, between residues and the π-system of the chromophores - and the inclusion of 10-15% exact-exchange would be warranted.

Further testing is merited, for instance, to check the behaviour of the DFT functionals to the peak shifts evidenced in the FSRS 3FMN* EAS.^17^ Furthermore, it should be investigated if the suggestions presented here are transferable to other well-documented chromophore and photoprotein/chromophore systems. Overall, similar type of vibrational studies would benefit from comprehensive research on excited state scaling factors particularly for some long-range corrected functionals.

## Conclusions

An extensive resonance Raman study was presented here including forty-two different DFT functionals in combination with the double-zeta basis set cc-pVDZ, and the inclusion of diffuse functions in selected cases. Off-Resonance spectra were computed initially, and yielded adequate agreement with experiments. These were fine-tuned with specific scaling factors for each DFT functional, purposed to align the highest computed peak (symmetric C=O stretch) to the corresponding experimental peak in the fingerprint region. Subsequently, resonance Raman intensities were computed with a careful choice of the resonant states. Since the experimental EAS spectrum utilized for the comparisons was taken under resonance conditions, inclusion of resonance enhancements in the calculations improved the agreement in most of the functionals. For six particular functionals, namely HCTH/407, OLYP, OPBE, O3LYP, tHCTHhyb and TPSSh, the correlation proved particularly facile, and the agreement was superior to the other functionals. The narrowing down to the above functionals resulted by the evaluation according to five criteria ranging from the percent error of the main flavin excitations, to visual inspection of the spectra and the improvement of the agreement with experiment with the inclusion of resonance enhancement. Owing to the extent of the data set a few conclusions can be drawn to assist similar studies. The following strategies can be beneficial in the study of excited state resonance Raman calculations:

- For the rR approach utilized in this study, steps 1-6 are required for the computation of resonance Raman spectra. The steps 3.1-3.3 can be optional but were useful in the benchmarking context.
- The usage of diffuse functions in the chosen basis set is recommended, as is for all the excited state calculations, but can be avoided, specifically in the case of large protein cluster calculations with a careful choice of DFT functional. Additionally, as was shown in the B3LYP basis set dependence study, augmentation of the double-zeta basis set is preferable rather than increase in size to triple- or quadruple-ζ.
- Moreover, in the absence of S_1_→S_n_ transitions within the selected resonance window, augmentation of the basis set can solve the issue.
- In case of established marker bands in the studied system, these can help in the definition of a Specific scaling factor, which in turn can assist in the assignments of off-Resonance spectra. Long range corrected functionals required scaling of their frequencies and re-assignment. This improved the correlation more than mere application of the scaling factor to correlated vibrations established by the unscaled
- frequencies. Scaling was not required for required for the majority of computed resonance Raman spectra.
- Setting the incident energy to the reference/resonant 0-0 transition, seems an acceptable choice. Inclusion of the actual experimental value had minimal impact on the computed intensities in this study.
- No particular class of DFT functionals has an advantage in the rR calculations, as members of the GGA, hybrid and meta-hybrid categories performed well in the benchmark. However, in the case of hybrid functionals, small percentage of HF exchange seems more favourable, judging from functionals such as BMK and BHandHLYP with increased percentage.
- Although not tested here, pre-Resonance calculations can give a good agreement,^17^ particularly if resonance conditions are not fully met in the experiments. Inspection of the experimental Transient Absorption spectrum can establish the actual conditions.

The results presented here should be, conceivably, an encouragement for further testing, both with the same flavin system under study, for example with the consideration of the triplet state spectra, and also other chromophore/photoprotein systems.

## Supporting information

Electronic Supplementary Information (ESI)

## Acknowledgements

P.C.A. was supported by the following grants: The project Structural dynamics of biomolecular systems (ELIBIO) (CZ.02.1.01/0.0/0.0/15_003/0000447) from the European Regional Development Fund and the Ministry of Education, Youth and Sports (MEYS) of the Czech Republic (https://www.msmt.cz/). The project of National Institute of Virology and Bacteriology, Programme EXCELES, funded by the European Union, Next Generation EU https://nivb.cz/ (LX22NPO5103). The Institute of Biotechnology of the Czech Academy of Sciences institutional grant RVO86652036 (https://www.msmt.cz/). H. H. was supported by a Deutscher Akademischer Austauschdienst e.V. (DAAD) scholarship. Computational resources were provided by the project “e-Infrastruktura CZ” (e-INFRA CZ LM2018140) supported by the Ministry of Education, Youth and Sports of the Czech Republic (https://www.cesnet.cz/projekty/e-infra_cz/) Computational resources were also provided by the ELIXIR-CZ project (LM2018131), part of the international ELIXIR infrastructure (https://www.cesnet.cz/projekty/elixir/). The funders had no role in study design, data collection and analysis, decision to publish, or preparation of the manuscript.

## Conflicts of interest

There are no conflicts of interest to declare.

**Scheme 1.**
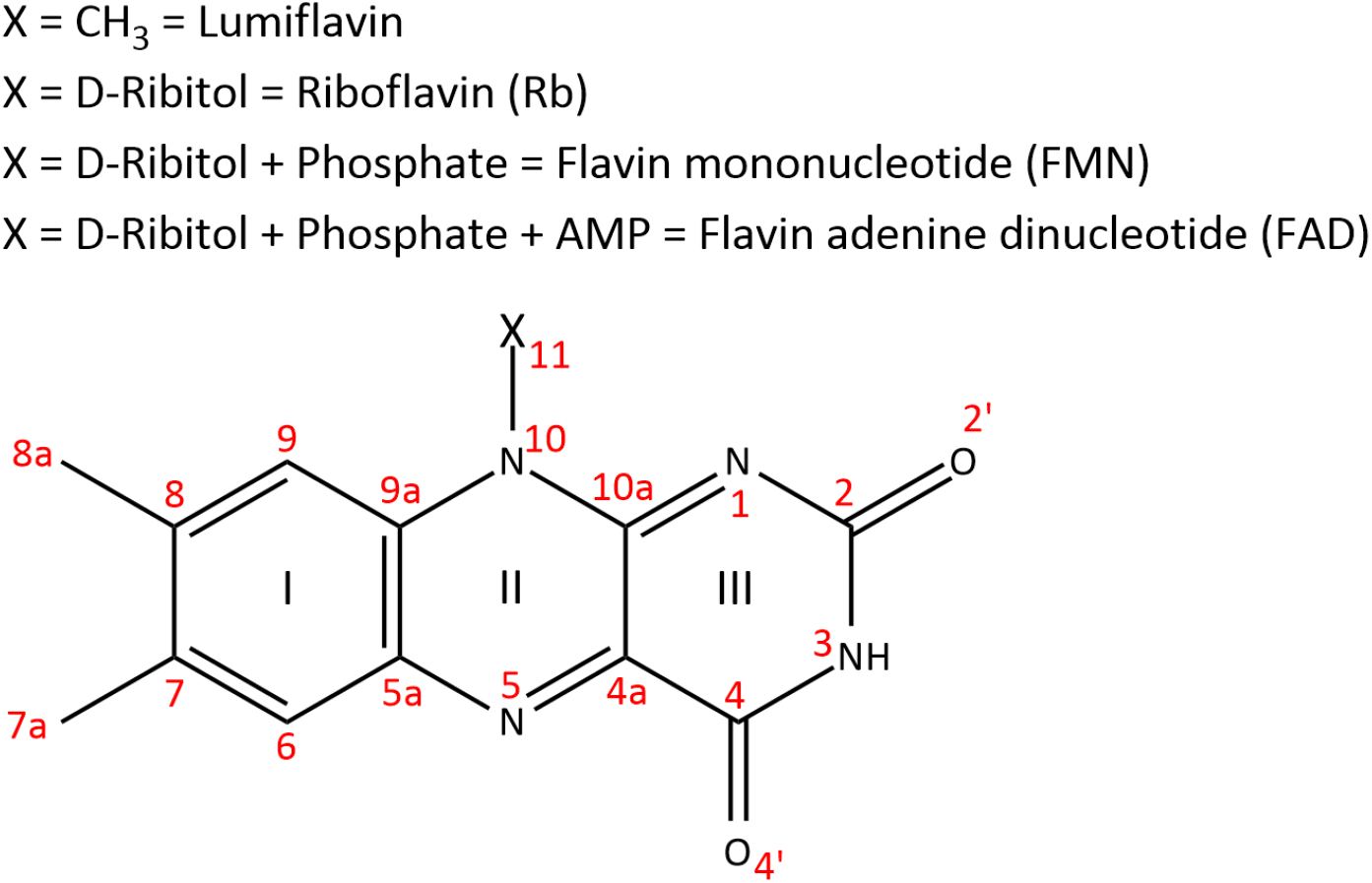
Numbering of atoms (in red), rings (in Latin numbers) and substitutions (X) in the family of Flavin chromophores. Depending on the X substitution at N_10_, a different member of the family is defined starting from the simplest lumiflavin, up to FAD.

**Scheme 2.**
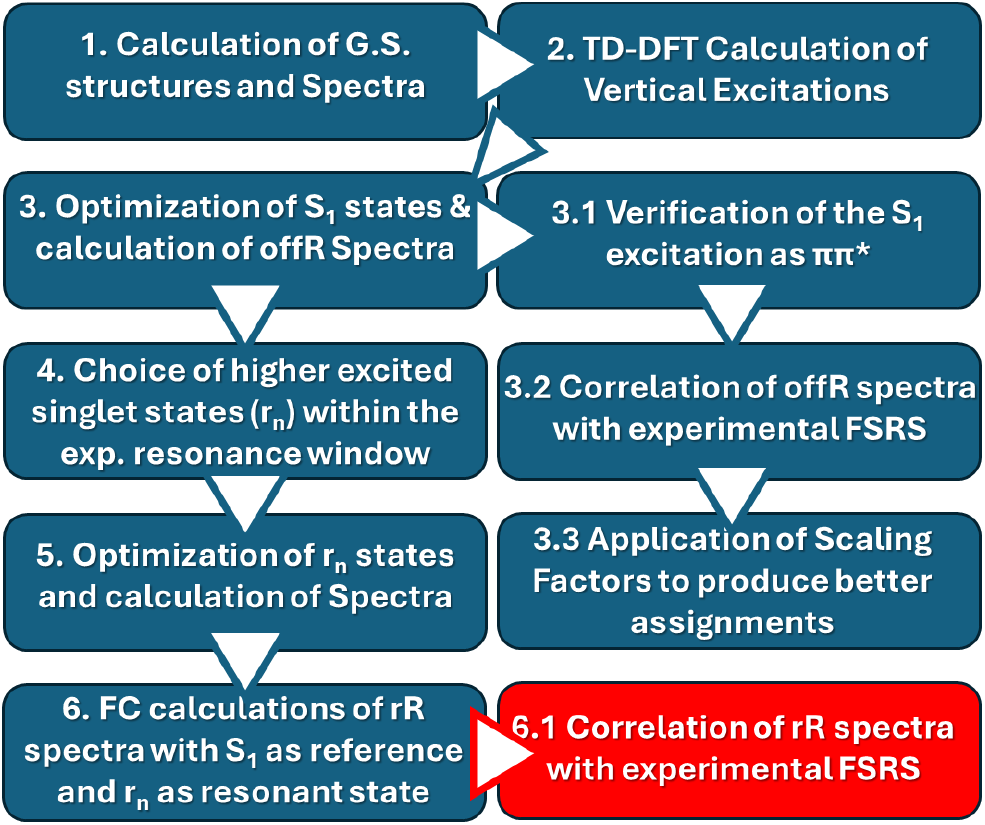
Schematic representation of the computational regime followed in this study.

