## Supplementary material for "Extensive Benchmark Study of the Resonance Raman Spectra of Lumiflavin": Electronic Supplementary Information (ESI)

### CONTENTS


#### 1. Classification, Dispersion and Scaling factors of DFT Functionals

Page 2 **Table S1** DFT Functionals included in the study, short description and citations.
Page 3 **Table S2** Dispersion Corrections included for each functional.
Page 4 **Table S3** Literature and calculated Scaling factors for each DFT functional.

#### 2. Data Tables on the Calculated Excited States for each DFT functional

Page 5-6 **Table S4** Comparison of Excitation energies for each DFT functional to the Experimental Absorption spectra.
Page 6 **Figure S1** Agreement between the experimental Absorption bands of FMN and the calculated excitation energies for all the DFT functionals.
Page 7 **Table S5** Typical vibrations of lumiflavin in the fingerprint region.
Page 7-9 **Table S6** Choice of excitations from $S_1$ to higher $r_n$ states within the Resonance window
Page 9-11 **Table S7** Hole/Electron properties on the $S_1$ and resonant states calculated for each DFT Functional.
Page 12-13 **Table S8** Hole and Electron surfaces for resonant states calculated for each DFT Functional.
Page 14-27 **Table S9** Assignment Tables between the peaks of the experimental FSRS $S_1$ EAS Spectrum and the calculated $S_1$ (off-Resonance) spectra for each DFT Functional.

#### 3. References

### 1. Classification, Dispersion and Scaling factors of DFT Functionals

**Table S1** Classification of the forty-two studied DFT functionals including their type along the DFT Jacob's Ladder, percentage of HF-exchange (% Ex.) and short description including citations.

| Functional | Type | % Ex. | Description + Citations |
|---|---|---|---|
| LSDA | LDA | - | Synonym of SVWN, combining the exchange Slater **S**[1] and correlation **VWN** [2] functionals |
| BLYP | GGA | - | GGA functional combining the exchange **B**[3] and correlation **LYP**[4-5] functionals |
| BP86 | GGA | - | GGA Functional combining the exchange **B**[3] and correlation **P86**[6] functionals |
| BPBE | GGA | - | GGA Functional combining the exchange **B**[3] and **PBE**[7-8] correlation functionals |
| HCTH/407 | GGA | - | Handy's GGA functional including gradient-corrected correlation[9-11] |
| mPWLYP | GGA | - | GGA functional combining the exchange **mPW**[12] and correlation **LYP**[4-5] functionals |
| OLYP | GGA | - | GGA functional combining Handy's **O**PTX modification[13-14] of the B exchange and the **LYP**[4-5] correlation functional |
| OPBE | GGA | - | GGA functional combining Handy's **O**PTX[13-14] modification and the **PBE**[7-8] correlation functionals |
| SOGGA11 | GGA | - | GGA functional from the Truhlar group[15] |
| M06L | meta-GGA | - | meta-GGA functional from the Truhlar group[16] |
| M11L | meta-GGA | - | meta-GGA functional from the Truhlar group[17] |
| revTPSS | meta-GGA | - | The revised **revTPSS** exchange and correlation functionals of Perdew *et. al.*[18-19] |
| TPSS1KCIS | meta-GGA | - | meta-GGA functional combining the exchange **TPSS**[20] and correlation **KCIS**[21-24] Krieger-Chen-Iafrate-Savin correlation functional |
| TPSSTPSS | meta-GGA | - | meta-GGA functional combining the **TPSS**[20] exchange and correlation functionals of Tao, Perdew, Staroverov, and Scuseria |
| VSXC | meta-GGA | - | van Voorhis and Scuseria τ-dependent gradient-corrected correlation functional[25] |
| APFD | Hybrid | 23 | The **A**ustin-**F**risch-**P**etersson hybrid functional[26] |
| B1B95 | Hybrid | 28 | **B**ecke's one-parameter hybrid functional[27] |
| B3LYP | Hybrid | 20 | Part of Becke Three-Parameter Hybrid Functionals[28] that follow the formulation: $A*E_X^{Slater}+(1-A)*E_X^{HF}+B*\Delta E_X^{Becke}+E_C^{VWN}+C*\Delta E_C^{non\text{-}local}$ where the non-local correlation is provided by the **LYP**,[4-5] **P86**[6] and **PW91**[29-31] correlation functionals |
| B3P86 | Hybrid | 20 | |
| B3PW91 | Hybrid | 20 | |
| B98 | Hybrid | 22 | Becke's 1998 revisions to B97[32-33] |
| BHHLYP | Hybrid | 50 | Half-and-Half Functional proposed by Becke[34] |
| mPW1PW91 | Hybrid | 25 | Hybrid Functional combining the **mPW** exchange modified by Adamo and Barone[12] combined with **PW91**[29-31] correlation functional |
| O3LYP | Hybrid | 11.6 | Hybrid three-parameter functional similar to B3LYP[28] with the A, B and C terms defined by Cohen & Handy[35] |
| PBE1PBE | Hybrid | 25 | The pure **PBE** functional[7-8] made into hybrid by Adamo[36] |
| SOGGA11x | Hybrid | 40.1 | Global hybrid functional from the Truhlar group[37] |
| X3LYP | Hybrid | 21.8 | Functional of **X**u and Goddard[38] |
| BMK | m-Hybrid | 42 | **B**oese and **M**artin's τ-dependent 2004 hybrid functional[39] |
| M05-2X | m-Hybrid | 56 | meta-Hybrid functional from the Truhlar group[40] |
| M06 | m-Hybrid | 27 | meta-Hybrid functional from the Truhlar group[41] |
| M06-HF | m-Hybrid | 27 | meta-Hybrid functional from the Truhlar group[42-43] |
| MN15 | m-Hybrid | 44 | meta-Hybrid functional from the Truhlar group[44] |
| PW6B95D3 | m-Hybrid | 28 | meta-Hybrid functional from the Truhlar group[45] |
| revTPSSh | m-Hybrid | 15 | meta-Hybrid functional employing the revised **revTPSS** exchange and correlation functionals[18-19] |
| tHCTHhyb | m-Hybrid | 15 | Hybrid functional using the **tHCTH** functional[46] |
| TPSSh | m-Hybrid | 10 | meta-Hybrid functional using the **TPSS** functionals[20, 47] |
| LC-OPBE | RS-GGA | - | GGA functional combining **O**[13-14] exchange and **PBE**[7-8] correlation together with the **L**ong-range **C**orrection of Hirao and coworkers[48] |
| CAM-B3LYP | RS-Hybrid | 10 | Handy and coworkers' long-range-corrected version of B3LYP using the Coulomb-attenuating method[49] |
| HISSbPBE | RS-Hybrid | 10 | Middle-range corrected hybrid employing the **HISS** functional[50] |
| HSEH1PBE | RS-Hybrid | 25 | **HSE06** range-corrected hybrid functional[51-56] employing **PBE**[7-8] correlation. |
| LC-wHPBE | RS-Hybrid | 10 | Updated version of the long-range corrected hybrid **ωPBE** functional [54, 57-59] |
| wB97XD | RS-Hybrid | 10 | Long-range corrected hybrid functional from Head-Gordon and coworkers[60] |

**Table S2** Empirical Dispersion correction parameters utilized for each DFT functional together with the source they were obtained from.

| Functional | S6 | S8 | SR6 | ABJ1 | ABJ2 | Type, Source |
|---|---|---|---|---|---|---|
| APFD | - | - | $R_{6APFD}$1.050 | - | - | PFD,[26] Gaussian[61] |
| B1B95 | 1.0000 | 1.4507 | - | 0.2092 | 5.5545 | GD3BJ, Grimme[62] |
| B3LYP | 1.0000 | 1.9889 | - | 0.3981 | 4.4211 | GD3BJ, Gaussian[61] |
| B3P86 | 1.0000 | 2.4830 | - | 0.5410 | 4.3060 | GD3BJ, MolSSI[63] |
| B3PW91 | 1.0000 | 2.8524 | - | 0.4312 | 4.4693 | GD3BJ, Gaussian[61] |
| B98 | 1.0000 | 0.7086 | - | -1.0000 | 6.0672 | GD3BJ, MolSSI[63] |
| BHHLYP | 1.0000 | 1.0354 | - | 0.2793 | 4.9615 | GD3BJ, Grimme[62] |
| BLYP | 1.0000 | 2.6996 | - | 0.4298 | 4.2359 | GD3BJ, Gaussian[61] |
| BMK | 1.0000 | 2.0860 | - | 0.1940 | 5.9197 | GD3BJ, Grimme[62] |
| BP86 | 1.0000 | 3.2822 | - | 0.3946 | 4.8516 | GD3BJ, Gaussian[61] |
| BPBE | 1.0000 | 4.0728 | - | 0.4567 | 4.3908 | GD3BJ, Gaussian[61] |
| CAM-B3LYP | 1.0000 | 2.0674 | - | 0.3708 | 5.4743 | GD3BJ, Gaussian[61] |
| HCTH/407 | 1.0000 | 1.0821 | - | 0.3563 | 4.3360 | GD3BJ, Grimme[62] |
| HISSbPBE | 1.0000 | 1.6112 | - | -1.0000 | 7.3539 | GD3BJ, MolSSI[63] |
| HSEH1PBE | 1.0000 | 2.3100 | - | 0.3830 | 5.6850 | GD3BJ, MolSSI[63] |
| LC-OPBE | 1.0000 | 3.3816 | - | 0.5512 | 2.9444 | GD3BJ, Grimme[62] |
| LC-wHPBE | 1.0000 | 1.8541 |  | 0.3919 | 5.0897 | GD3BJ, Gaussian[61] |
| LSDA | - | - | - | - | - | - |
| M05-2X | 1.0000 | 0.0000 | 1.4170 | - | - | GD3, Gaussian[61] |
| M06 | 1.0000 | 0.0000 | 1.3250 | - | - | GD3, Gaussian[61] |
| M06-HF | 1.0000 | 0.0000 | 1.4460 | - | - | GD3, Gaussian[61] |
| M06L | 1.0000 | 0.0000 | 1.5810 | - | - | GD3, Gaussian[61] |
| M11L | 1.0000 | 1.1129 | 2.3933 | - | - | GD3[64] |
| MN15 | 1.0000 | 2.0971 | - | 0.7862 | 7.5923 | GD3BJ, MolSSI[63] |
| mPW1PW91 | 1.0000 | 1.9467 | - | 0.3168 | 4.7732 | GD3BJ, MolSSI[63] |
| mPWLYP | 1.0000 | 2.0077 | - | 0.4831 | 4.5323 | GD3BJ, Grimme[62] |
| O3LYP | 1.0000 | 1.8171 | - | 0.0963 | 5.9940 | GD3BJ, MolSSI[63] |
| OLYP | 1.0000 | 2.6205 | - | 0.5299 | 2.8065 | GD3BJ, Grimme[62] |
| OPBE | 1.0000 | 3.3816 | - | 0.5512 | 2.9444 | GD3BJ, Grimme[62] |
| PBE1PBE | 1.0000 | 1.2177 | - | 0.4145 | 4.8593 | GD3BJ, Gaussian[61] |
| PW6B95D3 | 1.0000 | 0.7257 | - | 0.2076 | 6.3750 | GD3BJ, Gaussian[61] |
| revTPSSh | 1.0000 | 1.4076 | - | 0.2660 | 5.3761 | GD3BJ, MolSSI[63] |
| revTPSS | 1.0000 | 1.6151 | - | 0.2218 | 5.7985 | GD3BJ, MolSSI[63] |
| SOGGA11 | - | - | - | - | - | - |
| SOGGA11x | 1.0000 | 1.1426 |  | 0.1330 | 5.7381 | GD3BJ[64] |
| tHCTHhyb | 1.0000 | 1.0821 | - | 0.3563 | 4.3360 | GD3BJ, Grimme[62] |
| TPSS1KCIS | 10000 | 1.0542 | - | -1.0000 | 6.0201 | GD3BJ, MolSSI[63] |
| TPSSh | 1.0000 | 2.2382 | - | 0.4529 | 4.6550 | GD3BJ, Grimme[62] |
| TPSSTPSS | 1.0000 | 1.9435 | - | 0.4535 | 4.4752 | GD3BJ, Gaussian[61] |
| VSXC | - | - | - | - | - | - |
| wB97XD | 1.0000 | - | 1.1000 | - | - | GD2, Gaussian[61] |
| X3LYP | 1.0000 | 1.5744 | - | 0.2022 | 5.4184 | GD3BJ, MolSSI[63] |

**Table S3** Scaling factors (**Sc$_L$**) utilized to correct the spectra of each DFT functional along with the source they were obtained from. The specific scaling factor **Sc$_S$** is applied to each DFT calculated spectrum in order to align the $\nu_{75}$ C=O symmetric stretch vibration to the experimental FSRS 1FMN* 3$^{rd}$ EAS value of 1626 cm$^{-1}$.[65]

| Functional | Scaling Factor (Sc$_L$)<br>cc-pVDZ/aug-cc-pVDZ | Specific Sc.F. (Sc$_S$)<br>cc-pVDZ/aug-cc-pVDZ | Source |
|---|---|---|---|
| APFD | 0.9545 | 0.9322 | Calculated in this work with FREQ[66-68] |
| B1B95 | 0.9612 | 0.9222 | CCCBDB[69] |
| B3LYP | 0.9700/0.9704/0.9571/<br>0.9585/1.0000 | 0.9461/0.9776/0.9647/<br>0.9769/0.9706 | CCCBDB[69] |
| B3P86 | 0.9572 | 0.9334 | Laury *et al*[70] |
| B3PW91 | 0.9650 | 0.9352 | CCCBDB[69] |
| B98 | 0.9710 | 0.9383 | Tantirungrotechai *et al*[71] |
| BHHLYP | 0.9328/0.9326 | 0.8922/0.9210 | Laury *et al*[70] |
| BLYP | 1.0016 | 0.9896 | CCCBDB[69] |
| BMK | 0.9588/0.9588 | 0.8985/0.9277 | Merrick *et al*[72] |
| BP86 | 1.0006 | 0.9764 | Kesharwani *et al*[73] |
| BPBE | 0.9869 | 0.9728 | Calculated in this work with FREQ[66-68] |
| CAM-B3LYP | 0.9530 | 0.9160 | Calculated in this work with FREQ[66-68] |
| HCTH/407 | 0.9721 | 0.9502 | Laury *et al*[70] |
| HISSbPBE | 0.9283 | 0.8971 | Calculated in this work with FREQ[66-68] |
| HSEH1PBE | 0.9619 | 0.9272 | CCCBDB[69] |
| LC-OPBE | 0.9300 | 0.8554 | Calculated in this work with FREQ[66-68] |
| LC-wHPBE | 0.9417 | 0.8932 | Calculated in this work with FREQ[66-68] |
| LSDA | 0.9890/0.9887 | 0.9401/0.9554 | CCCBDB[69] |
| M05-2X | 0.9495/0.9501 | -/0.9395 | Laury *et al*[70] |
| M06 | 0.9670/0.9675 | 0.9187/0.9468 | Laury *et al*[70] |
| M06-HF | 0.9584 | 0.9187 | Calculated in this work with FREQ[66-68] |
| M06L | 0.9630/0.9630 | 0.9232/0.9593 | Kesharwani *et al*[73]/Palafox[74] |
| M11L | 0.9616 | 0.9092 | Calculated in this work with FREQ[66-68] |
| MN15 | 0.9512/0.9563 | 0.9148/0.9441 | Calculated in this work with FREQ[66-68] |
| mPW1PW91 | 0.9583 | 0.9249 | CCCBDB[69] |
| mPWLYP | 0.9953 | 0.9890 | Calculated in this work with FREQ[66-68] |
| O3LYP | 0.9696 | 0.9427 | Tantirungrotechai *et al*[71] |
| OLYP | 0.9875 | 0.9581 | Tantirungrotechai *et al*[71] |
| OPBE | 0.9702 | 0.9428 | Calculated in this work with FREQ[66-68] |
| PBE1PBE | 0.9615 | 0.9242 | CCCBDB[69] |
| PW6B95D3 | 0.9502 | 0.9252 | Calculated in this work with FREQ[66-68] |
| revTPSSh | -/0.9239 | -/0.8947 | Calculated in this work with FREQ[66-68] |
| revTPSS | -/0.9798 | -/0.9993 | Calculated in this work with FREQ[66-68] |
| SOGGA11 | 0.9788 | 0.9580 | Calculated in this work with FREQ[66-68] |
| SOGGA11x | 0.9403 | 0.9065 | Calculated in this work with FREQ[66-68] |
| tHCTHhyb | 0.9663 | 0.9449 | Calculated in this work with FREQ[66-68] |
| TPSS1KCIS | 0.9767 | - | Calculated in this work with FREQ[66-68] |
| TPSSh | 0.9720 | 0.9532 | CCCBDB[69] |
| TPSS | 0.9756/0.9801 | 0.9593/1.0018 | Calculated in this work with FREQ[66-68] |
| VSXC | 0.9770/0.9758 | 0.9424/0.9580 | Tantirungrotechai *et al*[71] |
| wB97XD | 0.9526 | 0.9124 | CCCBDB[69] |
| X3LYP | 0.9614 | 0.9423 | Calculated in this work with FREQ[66-68] |

### 2. Data Tables on the Calculated Excited States for each DFT functional

**Table S4** Values and statistical analysis pertaining to the main excitation energies of lumiflavin for the different DFT functionals. The terms in the table include the **$S_0 \rightarrow S_1$** and **$S_0 \rightarrow S_2$** experimental absorption and computed excitation values (in nm), corresponding deviations (**$\sigma_{S1}$**, **$\sigma_{S2}$**), absolute mean deviation (**$\mu_\sigma$**), and individual (**$\delta_{S1}$**, **$\delta_{S2}$**) and averaged (**$\mu_\delta$**) percent errors. For comparison, the full FMN values calculated at the B3LYP/def2-TZVP level are provided.

| LOT | $S_0 \rightarrow S_1$ | $S_0 \rightarrow S_2$ | $\sigma_{S1}$ | $\sigma_{S2}$ | $\mu_\sigma$ | $\delta_{S1}$ (%) | $\delta_{S2}$ (%) | $\mu_\delta$ (%) |
|---|---|---|---|---|---|---|---|---|
| **Experimental Abs.** | **445** | **372** | | | | | | |
| **FMN B3LYP/def2-TZVP 2'**[75] | 433 | 364 | -12 | -8 | 10 | 2.1 | 2.8 | 2.4 |
| **FMN B3LYP/def2-TZVP 2''**[75] | 431 | 366 | -15 | -6 | 10 | 1.6 | 3.4 | 2.5 |
| APFD/cc-pVDZ | 413 | 333 | -32 | -39 | 35 | 37.4 | 29.0 | 33.2 |
| B1B95/cc-pVDZ | 405 | 326 | -40 | -46 | 43 | 3.8 | 9.5 | 6.6 |
| B3LYP/cc-pVDZ | 423 | 341 | -22 | -31 | 27 | 9.1 | 5.2 | 7.1 |
| B3LYP/aug-cc-pVDZ | 433 | 354 | -12 | -18 | 15 | 5.1 | 2.8 | 4.0 |
| B3LYP/cc-pVTZ | 423 | 345 | -22 | -27 | 25 | 7.9 | 5.3 | 6.6 |
| B3LYP/aug-cc-pVTZ | 427 | 350 | -18 | -22 | 20 | 6.3 | 4.1 | 5.2 |
| B3LYP/cc-pVQZ | 425 | 347 | -20 | -25 | 22 | 7.1 | 4.8 | 5.9 |
| B3P86/cc-pVDZ | 421 | 339 | -24 | -33 | 28 | 24.2 | 19.1 | 21.6 |
| B3PW91/cc-pVDZ | 421 | 339 | -24 | -33 | 28 | 12.9 | 9.0 | 11.0 |
| B98/cc-pVDZ | 418 | 338 | -27 | -34 | 31 | 1.0 | 6.7 | 3.9 |
| BHHLYP/cc-pVDZ | 357 | 294 | -88 | -79 | 83 | 5.5 | 0.6 | 3.1 |
| BHHLYP/aug-cc-pVDZ | 366 | 305 | -79 | -67 | 73 | 4.0 | 8.6 | 6.3 |
| BLYP/cc-pVDZ | 494 | 389 | 49 | 17 | 33 | 1.9 | 6.8 | 4.3 |
| BMK/cc-pVDZ | 379 | 304 | -66 | -68 | 67 | 18.1 | 15.0 | 16.6 |
| BMK/aug-cc-pVDZ | 387 | 315 | -58 | -57 | 58 | 22.2 | 17.5 | 19.9 |
| BP86/cc-pVDZ | 492 | 387 | 47 | 15 | 31 | 11.3 | 7.7 | 9.5 |
| BPBE/cc-pVDZ | 490 | 386 | 45 | 14 | 29 | 2.0 | 8.1 | 5.1 |
| CAM-B3LYP/cc-pVDZ | 374 | 300 | -71 | -72 | 72 | 12.6 | 9.1 | 10.8 |
| HCTH/407/cc-pVDZ | 478 | 379 | 33 | 7 | 20 | 22.1 | 21.5 | 21.8 |
| HISSbPBE/cc-pVDZ | 377 | 310 | -68 | -62 | 65 | 3.5 | 9.2 | 6.3 |
| HSEH1PBE/cc-pVDZ | 413 | 334 | -32 | -38 | 35 | 24.3 | 18.8 | 21.5 |
| LC-OPBE/cc-pVDZ | 333 | 262 | -112 | -110 | 111 | 4.4 | 9.9 | 7.1 |
| LC-wHPBE/cc-pVDZ | 345 | 271 | -100 | -101 | 101 | 26.1 | 21.6 | 23.8 |
| LSDA/cc-pVDZ | 484 | 380 | 39 | 8 | 24 | 9.9 | 9.0 | 9.5 |
| LSDA/aug-cc-pVDZ | 495 | 391 | 50 | 19 | 35 | 21.7 | 18.8 | 20.2 |
| M05-2X/cc-pVDZ | 366 | 295 | -79 | -77 | 78 | 1.1 | 2.7 | 0.8 |
| M05-2X/aug-cc-pVDZ | 375 | 306 | -70 | -66 | 68 | 3.6 | 0.9 | 1.3 |
| M06/cc-pVDZ | 408 | 338 | -37 | -34 | 35 | 9.9 | 9.0 | 9.5 |
| M06/aug-cc-pVDZ | 418 | 350 | -27 | -22 | 24 | 4.5 | 1.6 | 3.1 |
| M06-HF/cc-pVDZ | 408 | 338 | -37 | -34 | 35 | 6.3 | 6.4 | 6.4 |
| M06L/cc-pVDZ | 449 | 359 | 4 | -13 | 8 | 18.2 | 14.0 | 16.1 |
| M06L/aug-cc-pVDZ | 457 | 368 | 12 | -4 | 8 | 4.6 | 0.6 | 2.6 |
| M11L/cc-pVDZ | 438 | 356 | -7 | -16 | 12 | 2.3 | 7.4 | 4.9 |
| MN15/cc-pVDZ | 390 | 315 | -55 | -57 | 56 | 2.2 | 6.8 | 4.5 |
| MN15/aug-cc-pVDZ | 400 | 327 | -45 | -45 | 45 | 1.0 | 4.5 | 1.7 |
| mPW1PW91/cc-pVDZ | 408 | 330 | -37 | -42 | 40 | 14.0 | 9.9 | 11.9 |
| mPWLYP/cc-pVDZ | 494 | 389 | 49 | 17 | 33 | 4.8 | 0.6 | 2.7 |
| O3LYP/cc-pVDZ | 442 | 355 | -3 | -17 | 10 | 12.8 | 9.1 | 10.9 |
| OLYP/cc-pVDZ | 483 | 381 | 38 | 9 | 24 | 18.2 | 14.0 | 16.1 |
| OPBE/cc-pVDZ | 480 | 378 | 35 | 6 | 20 | 11.4 | 7.7 | 9.6 |
| PBE1PBE/cc-pVDZ | 408 | 330 | -37 | -42 | 40 | 9.7 | 5.7 | 7.7 |
| PW6B95D3/cc-pVDZ | 405 | 327 | -40 | -45 | 43 | 10.2 | 6.4 | 8.3 |
| revTPSSh/aug-cc-pVDZ | 404 | 330 | -41 | -42 | 41 | 10.1 | 6.4 | 8.3 |
| revTPSS/aug-cc-pVDZ | 485 | 385 | 40 | 13 | 26 | 13.9 | 9.8 | 11.9 |
| SOGGA11/cc-pVDZ | 481 | 381 | 36 | 9 | 22 | 20.0 | 18.0 | 19.0 |
| SOGGA11x/cc-pVDZ | 378 | 308 | -67 | -64 | 66 | 1.8 | 7.5 | 4.7 |

| | | | | | | | | |
|---|---|---|---|---|---|---|---|---|
| tHCTHhyb/cc-pVDZ | 436 | 351 | -9 | -21 | 15 | 4.5 | 1.7 | 3.1 |
| TPSS1KCIS/cc-pVDZ# | 481 | 379 | 36 | 7 | 22 | 3.4 | 8.2 | 5.8 |
| TPSSh/cc-pVDZ | 442 | 353 | -3 | -19 | 11 | 12.6 | 10.3 | 11.4 |
| TPSSTPSS/cc-pVDZ | 477 | 376 | 32 | 4 | 18 | 3.4 | 8.2 | 5.8 |
| TPSSTPSS/aug-cc-pVDZ | 487 | 387 | 42 | 15 | 29 | 4.9 | 10.2 | 7.5 |
| VSXC/cc-pVDZ | 466 | 368 | 21 | -4 | 12 | 5.1 | 2.8 | 4.0 |
| VSXC/aug-cc-pVDZ | 477 | 380 | 32 | 8 | 20 | 9.1 | 5.2 | 7.1 |
| wB97XD/cc-pVDZ | 375 | 299 | -70 | -73 | 71 | 7.9 | 5.3 | 6.6 |
| X3LYP/cc-pVDZ | 418 | 338 | -27 | -34 | 30 | 6.3 | 4.1 | 5.2 |

**Figure S1** Deviation values (**σ**) for the $S_0 \rightarrow S_1$ (light red), for $S_0 \rightarrow S_2$ (light blue) and absolute mean deviations (**$\mu_\sigma$**, black) for the different functionals. All values are given in nm and are based on the values of **Table S4**. Functionals are sorted from top to bottom from higher **$\mu_\sigma$** values (worse) to lower (better agreement with experiment).

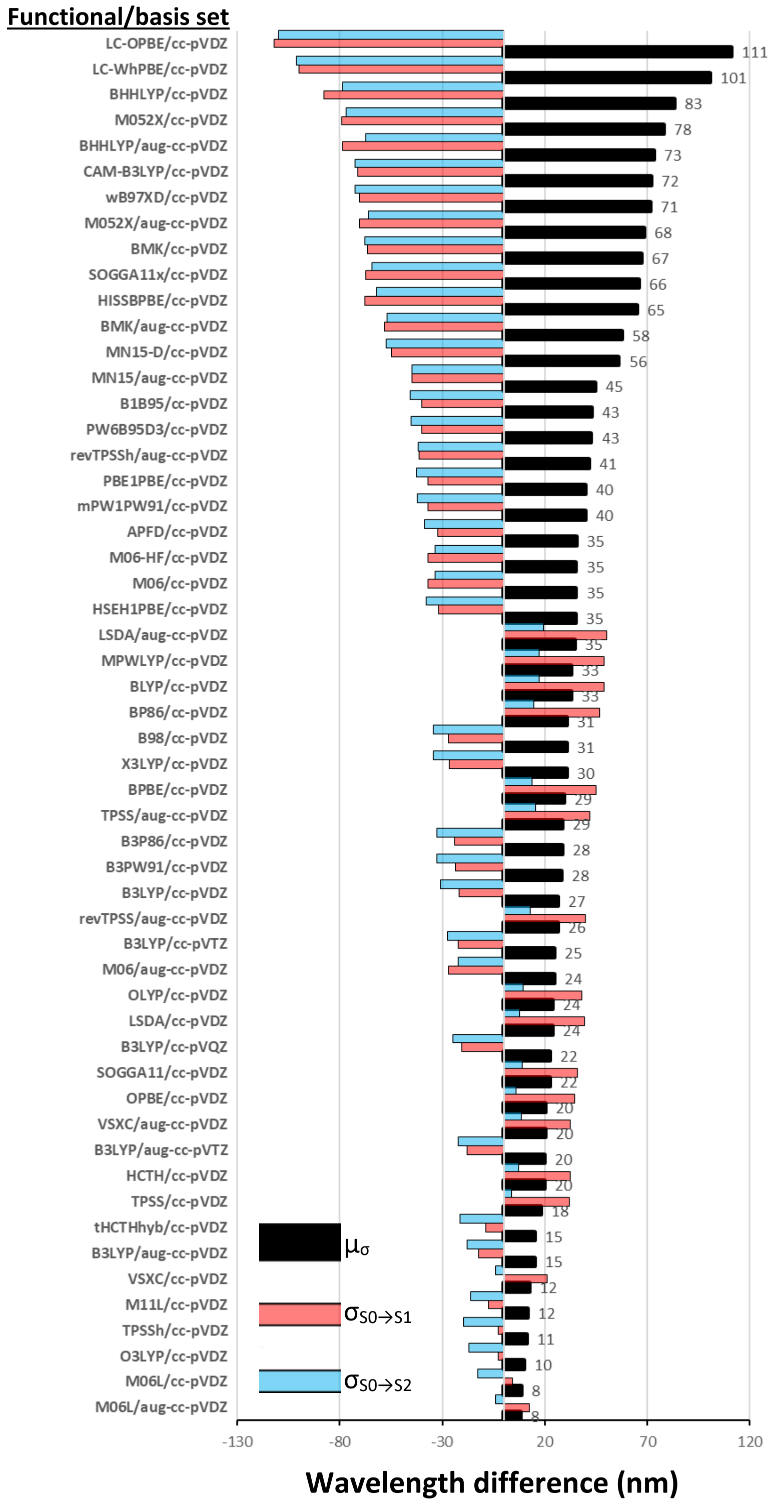

**Table S5** Typical vibrations showing the displacement vector arrows in the fingerprint region of lumiflavin, based on the B3LYP/aug-cc-pVDZ $S_1$ calculation.

| | | | | |
|---|---|---|---|---|
| 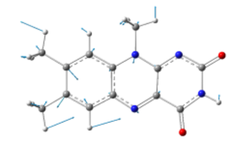 | 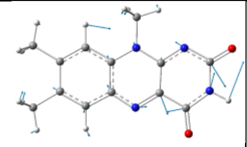 | 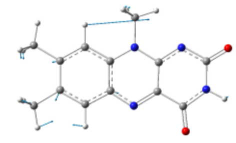 | 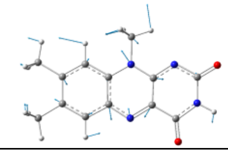 | 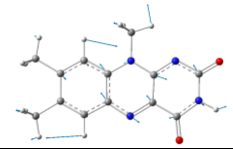 |
| $v_{48}$ | $v_{51}$ | $v_{52}$ | $v_{53}$ | $v_{54}$ |
| 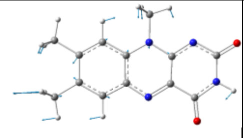 | 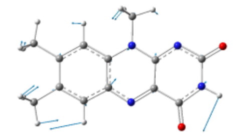 | 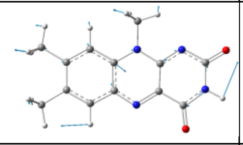 | 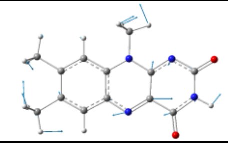 | 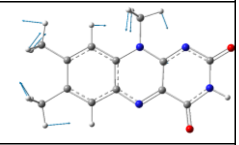 |
| $v_{56}$ | $v_{57}$ | $v_{61}$ | $v_{64}$ | $v_{65}$ |
| 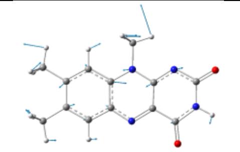 | 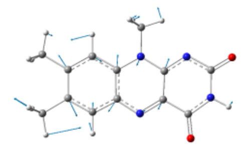 | 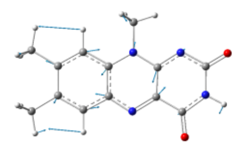 | 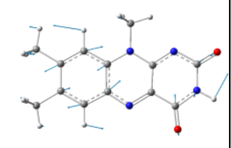 | 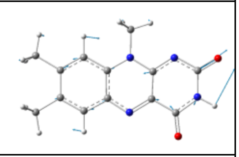 |
| $v_{70}$ | $v_{71}$ | $v_{72}$ | $v_{73}$ | $v_{74}$ |
| 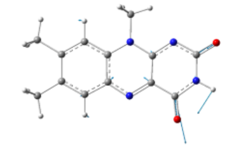 | | | | |
| $v_{75}$ | | | | |

**Table S6** Analysis of Excitations from the first excited state of lumiflavin ($S_1$) to higher singlet states $r_n$. Transition dipole moments (**TDM x, y, z**) are included, energies **EE** in eV and oscillator strengths **Osc.** as well as the difference **Δr** of the excitation from the wavelength of experimental Raman pump. Excitations choses as predominant at each LOT, after optimization are highlighted in bold.

| LOT | Excitation | TDM x | TDM y | TDM z | EE (eV) | Osc. | Δr (nm) |
|---|---|---|---|---|---|---|---|
| APFD/cc-pVDZ | **$S_1 \rightarrow r_5$** | -0.075 | 0.006 | 0.019 | 1.44 | 0.00021 | 4 |
| | $S_1 \rightarrow r_6$ | -0.357 | -0.544 | -0.032 | 1.55 | 0.01615 | 69 |
| B1B95/cc-pVDZ | $S_1 \rightarrow r_5$ | 0.032 | -0.004 | 0.007 | 1.56 | 0.00004 | 72 |
| | $S_1 \rightarrow r_6$ | -0.171 | -0.620 | -0.018 | 1.64 | 0.01665 | 119 |
| | **$S_1 \rightarrow r_7$** | -3.574 | 1.299 | -0.279 | 1.89 | 0.67387 | 259 |
| B3LYP/cc-pVDZ | $S_1 \rightarrow r_6$ | 0.009 | -0.699 | 0.000 | 1.47 | 0.01763 | 26 |
| | **$S_1 \rightarrow r_7$** | -3.518 | -1.642 | 0.000 | 1.76 | 0.65090 | 187 |
| B3LYP/aug-cc-pVDZ | $S_1 \rightarrow r_5$ | 0.001 | 0.000 | 0.008 | 1.56 | 0.00000 | 71 |
| | **$S_1 \rightarrow r_6$** | -0.943 | -0.292 | 0.000 | 1.61 | 0.03837 | 100 |
| | $S_1 \rightarrow r_6$ | 3.384 | -1.798 | 0.000 | 1.76 | 0.63213 | 184 |
| B3LYP/cc-pVTZ | $S_1 \rightarrow r_5$ | 0.001 | 0.000 | 0.011 | 1.52 | 0.00000 | 49 |
| | $S_1 \rightarrow r_6$ | -0.622 | -0.427 | 0.000 | 1.60 | 0.02228 | 94 |
| | **$S_1 \rightarrow r_7$** | 3.443 | -1.742 | 0.000 | 1.79 | 0.65149 | 200 |
| B3LYP/aug-cc-pVTZ | $S_1 \rightarrow r_5$ | -0.003 | -0.001 | -0.009 | 1.58 | 0.00000 | 82 |
| | $S_1 \rightarrow r_6$ | 0.987 | -0.258 | 0.000 | 1.63 | 0.04162 | 114 |
| | **$S_1 \rightarrow r_7$** | -3.352 | -1.789 | 0.000 | 1.79 | 0.63157 | 200 |
| B3LYP/cc-pVQZ | $S_1 \rightarrow r_5$ | 0.003 | 0.001 | 0.011 | 1.56 | 0.00001 | 76 |
| | **$S_1 \rightarrow r_6$** | 0.910 | -0.287 | 0.000 | 1.63 | 0.03627 | 111 |
| B3P86/cc-pVDZ | $S_1 \rightarrow r_6$ | -0.466 | -0.498 | -0.039 | 1.49 | 0.01702 | 33 |
| | $S_1 \rightarrow r_7$ | -3.636 | 1.380 | -0.283 | 1.77 | 0.65807 | 189 |
| B3PW91/cc-pVDZ | $S_1 \rightarrow r_6$ | -0.488 | -0.491 | -0.040 | 1.48 | 0.01744 | 28 |
| | **$S_1 \rightarrow r_7$** | -3.624 | 1.382 | -0.283 | 1.77 | 0.65427 | 189 |
| B98/cc-pVDZ | $S_1 \rightarrow r_6$ | -0.316 | -0.555 | -0.027 | 1.54 | 0.01547 | 63 |
| | **$S_1 \rightarrow r_7$** | -3.632 | 1.353 | -0.282 | 1.79 | 0.66419 | 205 |
| BHHLYP/cc-pVDZ | **$S_1 \rightarrow r_4$** | 0.020 | 0.003 | 0.005 | 1.60 | 0.00002 | 97 |
| BHHLYP/aug-cc-pVDZ | $S_1 \rightarrow r_4$ | 0.004 | -0.005 | -0.005 | 1.76 | 0.00000 | 188 |
| | **$S_1 \rightarrow r_5$** | -0.503 | 0.532 | -0.041 | 2.03 | 0.02675 | 336 |
| BLYP/cc-pVDZ | **$S_1 \rightarrow r_7$** | -3.338 | -1.562 | 0.000 | 1.49 | 0.49596 | 34 |

| LOT | Excitation | TDM x | TDM y | TDM z | EE (eV) | Osc. | Δr (nm) |
|---|---|---|---|---|---|---|---|
| | $S_1 \rightarrow r_8$ | 0.000 | -0.001 | -0.057 | 1.70 | 0.00013 | 154 |
| BMK/cc-pVDZ | **$S_1 \rightarrow r_5$** | 0.552 | -0.738 | 0.042 | 1.94 | 0.04041 | 285 |
| BMK/aug-cc-pVDZ | $S_1 \rightarrow r_4$ | 0.012 | 0.004 | 0.002 | 1.37 | 0.00001 | -32 |
| | **$S_1 \rightarrow r_5$** | -0.709 | 0.801 | -0.055 | 1.93 | 0.05418 | 279 |
| BP86/cc-pVDZ | **$S_1 \rightarrow r_7$** | -3.484 | 1.319 | -0.270 | 1.49 | 0.50759 | 32 |
| | $S_1 \rightarrow r_8$ | 0.022 | -0.013 | -0.052 | 1.71 | 0.00014 | 154 |
| BPBE/cc-pVDZ | **$S_1 \rightarrow r_7$** | -3.484 | 1.319 | -0.269 | 1.49 | 0.50852 | 33 |
| | $S_1 \rightarrow r_8$ | 0.033 | -0.015 | -0.052 | 1.69 | 0.00017 | 147 |
| CAM-B3LYP/cc-pVDZ | **$S_1 \rightarrow r_5$** | 0.802 | -0.102 | 0.000 | 1.98 | 0.03179 | 311 |
| HCTH/407/cc-pVDZ | **$S_1 \rightarrow r_7$** | -3.412 | -1.588 | 0.000 | 1.50 | 0.51938 | 38 |
| | $S_1 \rightarrow r_8$ | 0.000 | 0.000 | -0.055 | 1.69 | 0.00012 | 146 |
| HISSbPBE/cc-pVDZ | $S_1 \rightarrow r_5$ | -0.080 | 0.009 | 0.014 | 1.86 | 0.00031 | 243 |
| | $S_1 \rightarrow r_6$ | 0.163 | -0.672 | 0.011 | 1.90 | 0.02226 | 265 |
| | **$S_1 \rightarrow r_7$** | -3.539 | 1.384 | -0.277 | 2.06 | 0.73311 | 354 |
| HSEH1PBE/cc-pVDZ | $S_1 \rightarrow r_5$ | 0.048 | -0.001 | 0.007 | 1.42 | 0.00008 | -6 |
| | $S_1 \rightarrow r_6$ | -0.478 | -0.474 | -0.040 | 1.53 | 0.01710 | 58 |
| | **$S_1 \rightarrow r_7$** | -3.650 | 1.436 | -0.284 | 1.80 | 0.68145 | 207 |
| LC-OPBE/cc-pVDZ | $S_1 \rightarrow r_3$ | 0.128 | -0.593 | 0.007 | 1.22 | 0.01103 | -116 |
| | **$S_1 \rightarrow r_4$** | 0.011 | 0.002 | 0.008 | 1.67 | 0.00001 | 137 |
| LC-wHPBE/cc-pVDZ | $S_1 \rightarrow r_4$ | 0.000 | 0.000 | 0.009 | 1.49 | 0.00000 | 33 |
| | **$S_1 \rightarrow r_5$** | 0.140 | -0.090 | 0.000 | 2.07 | 0.00140 | 358 |
| LSDA/cc-pVDZ | $S_1 \rightarrow r_7$ | -3.442 | -1.615 | 0.000 | 1.48 | 0.52431 | 29 |
| | $S_1 \rightarrow r_8$ | 0.001 | 0.000 | -0.054 | 1.57 | 0.00011 | 79 |
| M05-2X/aug-cc-pVDZ | **$S_1 \rightarrow r_4$** | -0.023 | 0.005 | 0.007 | 1.44 | 0.00002 | 8 |
| | $S_1 \rightarrow r_5$ | -0.290 | 0.495 | -0.021 | 1.96 | 0.01584 | 298 |
| M06/cc-pVDZ | $S_1 \rightarrow r_5$ | 0.000 | 0.000 | -0.011 | 1.57 | 0.00000 | 77 |
| | $S_1 \rightarrow r_6$ | 1.214 | -0.062 | 0.000 | 1.71 | 0.06203 | 159 |
| | **$S_1 \rightarrow r_7$** | -3.301 | -1.732 | 0.000 | 1.85 | 0.62971 | 235 |
| | $S_1 \rightarrow r_8$ | 0.000 | 0.000 | -0.064 | 1.99 | 0.00020 | 317 |
| M06/aug-cc-pVDZ | **$S_1 \rightarrow r_5$** | 0.007 | 0.002 | -0.007 | 1.75 | 0.00000 | 180 |
| | $S_1 \rightarrow r_6$ | 2.539 | 0.669 | 0.000 | 1.77 | 0.29882 | 191 |
| | $S_1 \rightarrow r_7$ | -2.379 | -1.621 | 0.000 | 1.92 | 0.38895 | 272 |
| M06-HF/cc-pVDZ | $S_1 \rightarrow r_5$ | -0.002 | -0.001 | -0.011 | 1.57 | 0.00001 | 77 |
| | **$S_1 \rightarrow r_6$** | 1.213 | -0.062 | 0.000 | 1.71 | 0.06193 | 159 |
| M06L/cc-pVDZ | **$S_1 \rightarrow r_5$** | 1.124 | 0.096 | 0.086 | 1.45 | 0.04561 | 14 |
| | $S_1 \rightarrow r_6$ | -0.035 | 0.000 | -0.013 | 1.53 | 0.00005 | 55 |
| M06L/aug-cc-pVDZ | $S_1 \rightarrow r_5$ | -0.011 | -0.002 | 0.006 | 1.03 | 0.00000 | -222 |
| | $S_1 \rightarrow r_6$ | -0.707 | -0.368 | -0.057 | 1.27 | 0.01989 | -88 |
| | **$S_1 \rightarrow r_7$** | -3.700 | 1.444 | -0.288 | 1.52 | 0.58864 | 48 |
| M11L/cc-pVDZ | $S_1 \rightarrow r_6$ | -0.497 | -0.372 | -0.041 | 1.28 | 0.01210 | -85 |
| | $S_1 \rightarrow r_7$ | -3.773 | 1.459 | -0.293 | 1.51 | 0.60863 | 45 |
| MN15/cc-pVDZ | **$S_1 \rightarrow r_5$** | 0.052 | 0.001 | -0.010 | 1.81 | 0.00013 | 216 |
| | $S_1 \rightarrow r_6$ | 0.418 | -0.743 | 0.029 | 1.89 | 0.03359 | 256 |
| MN15/aug-cc-pVDZ | **$S_1 \rightarrow r_4$** | -0.083 | 0.011 | 0.014 | 1.23 | 0.00022 | -114 |
| | $S_1 \rightarrow r_5$ | -0.790 | 0.865 | -0.060 | 1.89 | 0.06373 | 259 |
| mPW1PW91/cc-pVDZ | $S_1 \rightarrow r_5$ | -0.037 | 0.005 | 0.015 | 1.51 | 0.00006 | 48 |
| | $S_1 \rightarrow r_6$ | -0.280 | -0.570 | -0.024 | 1.61 | 0.01590 | 101 |
| | **$S_1 \rightarrow r_7$** | -3.614 | 1.344 | -0.280 | 1.85 | 0.67928 | 238 |
| mPWLYP/cc-pVDZ | **$S_1 \rightarrow r_7$** | -3.343 | -1.565 | 0.000 | 1.49 | 0.49702 | 34 |
| | $S_1 \rightarrow r_8$ | 0.000 | 0.001 | -0.057 | 1.70 | 0.00013 | 153 |
| O3LYP/cc-pVDZ | **$S_1 \rightarrow r_7$** | -3.617 | 1.392 | -0.280 | 1.64 | 0.60777 | 120 |
| | $S_1 \rightarrow r_8$ | 0.043 | -0.006 | -0.060 | 1.87 | 0.00025 | 244 |
| OLYP/cc-pVDZ | **$S_1 \rightarrow r_7$** | -3.473 | 1.324 | -0.269 | 1.50 | 0.51151 | 42 |
| | $S_1 \rightarrow r_8$ | 0.025 | -0.014 | -0.053 | 1.72 | 0.00015 | 161 |
| OPBE/cc-pVDZ | **$S_1 \rightarrow r_7$** | 0.029 | -0.018 | -0.051 | 1.71 | 0.00016 | 155 |
| | $S_1 \rightarrow r_8$ | 2.265 | -1.801 | 0.172 | 0.84 | 0.17341 | -328 |
| PBE1PBE/cc-pVDZ | **$S_1 \rightarrow r_5$** | -0.001 | 0.000 | -0.013 | 1.50 | 0.00001 | 39 |
| | $S_1 \rightarrow r_6$ | 0.199 | -0.605 | 0.000 | 1.60 | 0.01587 | 95 |
| PW6B95D3/cc-pVDZ | $S_1 \rightarrow r_5$ | 0.081 | -0.008 | 0.001 | 1.57 | 0.00025 | 80 |
| | $S_1 \rightarrow r_6$ | -0.181 | -0.617 | -0.019 | 1.64 | 0.01665 | 119 |
| | **$S_1 \rightarrow r_7$** | -3.570 | 1.299 | -0.278 | 1.89 | 0.67277 | 259 |
| revTPSSh/aug-cc-pVDZ | $S_1 \rightarrow r_5$ | 0.000 | 0.000 | -0.015 | 1.53 | 0.00001 | 59 |
| | $S_1 \rightarrow r_6$ | 0.271 | -0.561 | 0.000 | 1.64 | 0.01562 | 120 |
| | **$S_1 \rightarrow r_7$** | -3.398 | -1.647 | 0.000 | 1.85 | 0.64449 | 233 |
| revTPSS/aug-cc-pVDZ | **$S_1 \rightarrow r_7$** | -3.439 | -1.626 | 0.000 | 1.52 | 0.53727 | 49 |

| LOT | Excitation | TDM x | TDM y | TDM z | EE (eV) | Osc. | Δr (nm) |
|---|---|---|---|---|---|---|---|
| | $S_1 \rightarrow r_8$ | 0.001 | 0.000 | -0.052 | 1.88 | 0.00012 | 254 |
| SOGGA11/cc-pVDZ | $S_1 \rightarrow r_7$ | -3.454 | 1.327 | -0.265 | 1.50 | 0.50535 | 39 |
| | $S_1 \rightarrow r_8$ | 0.096 | -0.026 | -0.053 | 1.55 | 0.00048 | 67 |
| | **$S_1 \rightarrow r_9$** | -0.085 | 0.006 | 0.007 | 1.77 | 0.00032 | 192 |
| SOGGA11x/cc-pVDZ | **$S_1 \rightarrow r_5$** | 0.953 | -0.906 | 0.073 | 1.97 | 0.08383 | 305 |
| tHCTHhyb/cc-pVDZ | $S_1 \rightarrow r_6$ | -0.064 | -0.722 | 0.000 | 1.37 | 0.01766 | -32 |
| | **$S_1 \rightarrow r_7$** | -0.002 | 0.000 | -0.064 | 1.93 | 0.00020 | 279 |
| TPSSh/cc-pVDZ | **$S_1 \rightarrow r_7$** | -3.494 | -1.595 | 0.000 | 1.66 | 0.60097 | 131 |
| | $S_1 \rightarrow r_8$ | 0.000 | 0.000 | -0.063 | 1.94 | 0.00019 | 286 |
| TPSS/cc-pVDZ | **$S_1 \rightarrow r_5$** | 0.697 | 0.703 | 0.000 | 1.43 | 0.03447 | 3 |
| | $S_1 \rightarrow r_6$ | 0.000 | -0.001 | -0.015 | 1.55 | 0.00001 | 65 |
| TPSS/aug-cc-pVDZ | **$S_1 \rightarrow r_7$** | -3.449 | -1.638 | 0.000 | 1.50 | 0.53468 | 38 |
| | $S_1 \rightarrow r_8$ | 0.002 | -0.001 | -0.051 | 1.84 | 0.00012 | 229 |
| VSXC/cc-pVDZ | **$S_1 \rightarrow r_5$** | 0.947 | -0.053 | 0.070 | 1.53 | 0.03397 | 58 |
| | $S_1 \rightarrow r_6$ | 0.024 | -0.003 | -0.017 | 1.58 | 0.00003 | 82 |
| VSXC/aug-cc-pVDZ | **$S_1 \rightarrow r_7$** | -3.668 | 1.407 | -0.282 | 1.50 | 0.57065 | 40 |
| | $S_1 \rightarrow r_8$ | 0.082 | -0.005 | -0.055 | 1.93 | 0.00046 | 279 |
| wB97XD/cc-pVDZ | $S_1 \rightarrow r_5$ | 0.674 | -0.687 | 0.051 | 2.00 | 0.04547 | 319 |
| | **$S_1 \rightarrow r_6$** | -0.025 | 0.005 | -0.033 | 2.14 | 0.00009 | 396 |
| X3LYP/cc-pVDZ | $S_1 \rightarrow r_6$ | -0.424 | -0.519 | -0.037 | 1.52 | 0.01680 | 52 |
| | **$S_1 \rightarrow r_7$** | -3.615 | 1.370 | -0.282 | 1.79 | 0.65973 | 203 |

**Table S7** Various hole/electron properties of all computed excited states for the various DFT functionals. **D** gives the distance of the centroids of the hole and electron in Å, **Sr** is the integral of hole and electron with 1 signifying perfect match, **H** is the overall measure of spatial distribution of the hole and electron in Å, **τ** is the measure of hole-electron separation in the CT direction, $\mathbf{E_{coul}}$ is the coulomb attractive energy between hole and electron, **HDI**/**EDI** are hole and electron delocalization indexes, respectively, **x,y,z** is the transition dipole moment in a.u., **EE** the excitation energy in eV (from the G.S.) and **Ex.** is the description of the excitation. Information about state and hole-electron properties is provided by the Multiwfn program.[76-77] State numbers are signified as "roots" (i.e. root 5 = $r_5$, root 7 = $r_7$). The first excited singlet state $S_1$ is usually the first root $r_1$.

| LOT | Excitation | D (Å) | Sr (a.u.) | H (Å) | t (Å) | $E_{coul}$ (eV) | HDI | EDI | x | y | z | EE (eV) | Ex. |
|---|---|---|---|---|---|---|---|---|---|---|---|---|---|
| APFD/cc-pVDZ | $S_0 \rightarrow S_1$ ($r_1$) | 0.889 | 0.693 | 2.861 | -1.064 | 5.20 | 8.94 | 9.39 | 0.993 | -2.063 | 0.068 | 2.52 | ππ* |
| | $S_0 \rightarrow r_5$ | 3.148 | 0.440 | 2.518 | 1.478 | 3.84 | 16.45 | 9.38 | 0.233 | 0.082 | 0.064 | 3.49 | nπ* |
| B1B95/cc-pVDZ | $S_0 \rightarrow S_1$ ($r_1$) | 0.857 | 0.697 | 2.852 | -1.091 | 5.23 | 8.92 | 9.38 | 1.024 | -2.106 | 0.070 | 2.57 | ππ* |
| | $S_0 \rightarrow r_5/r_6$ | 3.142 | 0.437 | 2.478 | 1.501 | 4.42 | 16.39 | 9.41 | 0.305 | 0.052 | 0.063 | 3.61 | CT |
| | $S_0 \rightarrow r_7$ | 1.042 | 0.709 | 3.037 | -0.937 | 4.88 | 7.50 | 9.28 | -0.424 | 0.873 | -0.025 | 4.22 | ππ* |
| B3LYP/cc-pVDZ | $S_0 \rightarrow S_1$ ($r_1$) | 0.939 | 0.682 | 2.871 | -0.411 | 5.37 | 9.21 | 9.51 | 2.132 | -0.706 | 0.000 | 2.45 | ππ* |
| | $S_0 \rightarrow r_7$ | 1.287 | 0.712 | 3.006 | -0.880 | 5.17 | 7.18 | 9.37 | -1.072 | 0.260 | 0.000 | 4.00 | ππ* |
| B3LYP/aug-cc-pVDZ | $S_0 \rightarrow S_1$ ($r_1$) | 0.858 | 0.693 | 2.928 | -0.517 | 5.30 | 8.57 | 9.23 | -2.159 | -0.745 | 0.001 | 2.40 | ππ* |
| | $S_0 \rightarrow r_5$ | 0.750 | 0.493 | 2.471 | -1.102 | 6.12 | 14.88 | 9.16 | -0.006 | 0.003 | 0.099 | 2.69 | nπ* |
| | $S_0 \rightarrow r_6/r_7$ | 3.237 | 0.446 | 2.504 | 1.410 | 4.51 | 15.75 | 9.30 | -0.398 | 0.368 | -0.042 | 3.48 | nπ* |
| B3LYP/cc-pVTZ | $S_0 \rightarrow S_1$ ($r_1$) | 0.862 | 0.693 | 2.885 | -0.455 | 5.37 | 8.74 | 9.45 | -2.144 | -0.725 | 0.000 | 2.45 | ππ* |
| | $S_0 \rightarrow r_7$ | 1.236 | 0.473 | 2.553 | -0.650 | 5.81 | 14.32 | 9.35 | -0.001 | 0.000 | -0.077 | 3.99 | nπ* |
| B3LYP/aug-cc-pVTZ | $S_0 \rightarrow S_1$ ($r_1$) | 0.854 | 0.694 | 2.913 | -0.545 | 5.33 | 8.54 | 9.27 | 2.158 | -0.742 | 0.000 | 2.42 | ππ* |
| | $S_0 \rightarrow r_5$ | 2.841 | 0.405 | 2.736 | 0.532 | 4.73 | 10.14 | 9.06 | 0.000 | 0.000 | 0.094 | 4.73 | nπ* |
| | $S_0 \rightarrow r_6$ | 3.239 | 0.442 | 2.491 | 1.422 | 4.52 | 15.92 | 9.35 | 0.409 | 0.365 | 0.043 | 3.51 | CT |
| | $S_0 \rightarrow r_7$ | 0.946 | 0.707 | 3.161 | -0.644 | 5.04 | 7.91 | 9.20 | -0.937 | 0.413 | -0.012 | 3.96 | ππ* |
| B3LYP/cc-pVQZ | $S_0 \rightarrow S_1$ ($r_1$) | 0.859 | 0.695 | 2.891 | -0.498 | 5.36 | 8.61 | 9.40 | 2.150 | -0.734 | 0.000 | 2.44 | ππ* |
| | $S_0 \rightarrow r_6$ | 3.239 | 0.440 | 2.481 | 1.590 | 4.37 | 16.01 | 9.39 | 0.498 | -0.034 | -0.010 | 3.51 | CT |
| B3P86/cc-pVDZ | $S_0 \rightarrow S_1$ ($r_1$) | 0.922 | 0.687 | 2.861 | -1.033 | 5.20 | 9.06 | 9.46 | 0.965 | -2.013 | 0.067 | 2.47 | ππ* |
| | $S_0 \rightarrow r_7$ | 1.290 | 0.708 | 3.012 | -0.656 | 4.84 | 7.15 | 9.33 | -0.560 | 0.936 | -0.033 | 4.03 | ππ* |
| B3PW91/cc-pVDZ | $S_0 \rightarrow S_1$ ($r_1$) | 0.925 | 0.684 | 2.872 | -1.041 | 5.75 | 9.14 | 9.47 | 0.970 | -2.016 | 0.067 | 2.47 | ππ* |
| | $S_0 \rightarrow$ **$r_6$** | 0.861 | 0.688 | 2.893 | -1.130 | 5.07 | 8.92 | 9.34 | -1.390 | 2.846 | -0.097 | 2.47 | ππ* |
| | $S_0 \rightarrow r_7$ | 1.262 | 0.711 | 3.012 | -0.686 | 4.24 | 7.16 | 9.33 | -0.562 | 0.937 | -0.033 | 4.03 | ππ* |
| B98/cc-pVDZ | $S_0 \rightarrow S_1$ ($r_1$) | 0.906 | 0.690 | 2.861 | -1.046 | 5.20 | 8.98 | 9.43 | 0.987 | -2.048 | 0.068 | 2.48 | ππ* |

| LOT | Excitation | D (Å) | Sr (a.u.) | H (Å) | t (Å) | $E_{coul}$ (eV) | HDI | EDI | x | y | z | EE (eV) | Ex. |
|---|---|---|---|---|---|---|---|---|---|---|---|---|---|
| | $S_0 \rightarrow$ **$r_6$** | 2.963 | 0.343 | 2.380 | 1.437 | 4.67 | 21.55 | 9.63 | -0.002 | 0.006 | -0.008 | 2.74 | CT |
| | $S_0 \rightarrow r_7$ | 1.157 | 0.713 | 3.042 | -0.814 | 4.84 | 7.27 | 9.23 | -0.507 | 0.930 | -0.029 | 4.05 | ππ* |
| BHHLYP/cc-pVDZ | $S_0 \rightarrow S_1 (r_1)$ | 0.603 | 0.736 | 2.786 | -1.279 | 5.37 | 8.01 | 9.07 | 1.239 | -2.407 | 0.086 | 2.89 | ππ* |
| | $S_0 \rightarrow r_4$ | 2.003 | 0.406 | 2.043 | 0.574 | 6.06 | 25.48 | 10.3 | 0.005 | 0.001 | -0.048 | 3.97 | nπ* |
| BHHLYP/aug-cc-pVDZ | $S_0 \rightarrow S_1 (r_1)$ | 0.593 | 0.737 | 2.839 | -1.302 | 5.26 | 7.75 | 8.76 | 1.267 | -2.481 | 0.088 | 2.82 | ππ* |
| | $S_0 \rightarrow r_4$ | 2.035 | 0.395 | 2.092 | 0.595 | 5.90 | 25.13 | 9.82 | -0.002 | -0.003 | 0.042 | 4.12 | nπ* |
| | $S_0 \rightarrow r_5$ | 0.418 | 0.810 | 3.007 | -1.542 | 5.05 | 7.78 | 7.50 | 2.770 | -1.175 | 0.228 | 4.61 | ππ* |
| BLYP/cc-pVDZ | $S_0 \rightarrow S_1 (r_2)$ | 1.386 | 0.613 | 2.925 | -0.208 | 5.14 | 10.62 | 9.83 | 1.828 | -0.573 | 0.000 | 2.05 | ππ* |
| | $S_0 \rightarrow r_7$ | 1.988 | 0.706 | 2.967 | -0.422 | 5.01 | 6.93 | 9.62 | -1.327 | 0.162 | -0.006 | 3.43 | ππ* |
| | $S_0 \rightarrow r_8$ | 1.017 | 0.465 | 2.590 | -0.962 | 5.85 | 14.81 | 9.01 | 0.000 | 0.000 | -0.079 | 3.29 | nπ* |
| BMK/cc-pVDZ | $S_0 \rightarrow S_1(r_1)/r_5$ | 0.747 | 0.717 | 2.830 | -1.182 | 5.28 | 8.51 | 9.20 | 1.116 | -2.268 | 0.076 | 2.74 | ππ* |
| BMK/aug-cc-pVDZ | $S_0 \rightarrow S_1 (r_1)$ | 0.701 | 0.724 | 2.880 | -1.239 | 5.19 | 8.19 | 8.90 | 1.153 | -2.345 | 0.079 | 2.68 | ππ* |
| | $S_0 \rightarrow r_4$ | 2.225 | 0.399 | 2.212 | 0.725 | 5.54 | 23.88 | 9.54 | 0.008 | -0.001 | -0.051 | 3.64 | nπ* |
| | $S_0 \rightarrow r_5$ | 0.479 | 0.776 | 2.884 | -1.291 | 5.18 | 8.50 | 7.95 | 2.500 | -1.214 | 0.373 | 4.05 | ππ* |
| BP86/cc-pVDZ | $S_0 \rightarrow S_1 (r_2)$ | 1.325 | 0.618 | 2.937 | -0.708 | 5.00 | 10.40 | 9.73 | 0.855 | -1.718 | 0.060 | 2.07 | ππ* |
| | $S_0 \rightarrow r_7$ | 1.990 | 0.703 | 2.976 | 0.025 | 4.63 | 6.84 | 9.54 | -0.808 | 1.051 | -0.054 | 3.45 | ππ* |
| | $S_0 \rightarrow r_8$ | 1.011 | 0.466 | 2.602 | -0.784 | 5.72 | 14.48 | 8.94 | 0.007 | 0.002 | -0.078 | 3.31 | nπ* |
| BPBE/cc-pVDZ | $S_0 \rightarrow S_1 (r_1)$ | 1.321 | 0.618 | 2.934 | -0.711 | 5.00 | 10.41 | 9.74 | 0.855 | -1.717 | 0.061 | 2.08 | ππ* |
| | $S_0 \rightarrow r_7$ | 1.993 | 0.703 | 2.973 | 0.029 | 4.63 | 6.85 | 9.55 | -0.805 | 1.046 | -0.053 | 3.46 | ππ* |
| | $S_0 \rightarrow r_8$ | 1.006 | 0.467 | 2.599 | -0.788 | 5.73 | 14.49 | 8.95 | 0.007 | 0.001 | -0.078 | 3.30 | nπ* |
| CAM-B3LYP/cc-pVDZ | $S_0 \rightarrow S_1 (r_1)$ | 0.659 | 0.732 | 2.787 | -0.651 | 5.59 | 8.41 | 9.33 | 2.562 | -0.761 | 0.000 | 2.76 | ππ* |
| | $S_0 \rightarrow r_5$ | 1.609 | 0.720 | 2.474 | -0.306 | 5.66 | 8.45 | 9.37 | 2.057 | 0.550 | 0.035 | 3.39 | ππ* |
| HCTH/407/cc-pVDZ | $S_0 \rightarrow S_1 (r_2)$ | 1.239 | 0.626 | 2.923 | -0.227 | 5.20 | 10.20 | 9.75 | 1.834 | -0.594 | 0.000 | 2.14 | ππ* |
| | $S_0 \rightarrow r_7$ | 1.934 | 0.703 | 2.964 | -0.470 | 5.04 | 6.94 | 9.52 | -1.266 | 0.149 | -0.012 | 3.53 | ππ* |
| | $S_0 \rightarrow r_8$ | 1.007 | 0.467 | 2.582 | -0.967 | 5.87 | 14.58 | 8.95 | -0.001 | -0.001 | -0.075 | 3.36 | nπ* |
| HISSbPBE/cc-pVDZ | $S_0 \rightarrow S_1 (r_1)$ | 0.702 | 0.719 | 2.858 | -1.213 | 5.24 | 8.27 | 9.14 | 1.121 | -2.250 | 0.077 | 2.76 | ππ* |
| | $S_0 \rightarrow r_6$ | 3.270 | 0.429 | 2.445 | 1.646 | 4.37 | 16.45 | 9.02 | 0.393 | 0.058 | 0.000 | 4.05 | CT |
| | $S_0 \rightarrow r_7$ | 0.892 | 0.705 | 3.093 | -1.179 | 4.84 | 8.08 | 9.15 | -0.136 | 0.744 | -0.001 | 4.48 | ππ* |
| HSEH1PBE/cc-pVDZ | $S_0 \rightarrow S_1 (r_1)$ | 0.866 | 0.693 | 2.880 | -1.099 | 5.18 | 8.88 | 9.29 | 0.992 | -2.055 | 0.068 | 2.52 | ππ* |
| | $S_0 \rightarrow r_7$ | 1.208 | 0.709 | 3.028 | -0.745 | 4.85 | 7.20 | 9.19 | -0.517 | 0.908 | -0.030 | 4.10 | ππ* |
| LC-OPBE/cc-pVDZ | $S_0 \rightarrow S_1 (r_1)$ | 0.442 | 0.755 | 2.685 | -1.378 | 5.58 | 8.06 | 9.22 | 1.441 | -2.572 | 0.102 | 3.02 | ππ* |
| | $S_0 \rightarrow r_4$ | 1.608 | 0.427 | 1.835 | 0.294 | 6.90 | 26.68 | 11.2 | -0.002 | 0.004 | -0.031 | 4.05 | nπ* |
| LC-wHPBE/cc-pVDZ | $S_0 \rightarrow S_1 (r_1)$ | 0.514 | 0.750 | 2.691 | -0.827 | 5.78 | 8.29 | 9.41 | 2.816 | -0.747 | 0.000 | 2.90 | ππ* |
| | $S_0 \rightarrow r_5$ | 0.282 | 0.812 | 2.894 | -2.016 | 5.63 | 8.19 | 7.63 | -2.686 | -1.111 | 0.067 | 4.79 | ππ* |
| LSDA/cc-pVDZ | $S_0 \rightarrow S_1 (r_2)$ | 1.230 | 0.625 | 2.949 | -0.277 | 5.17 | 10.20 | 9.71 | 1.817 | -0.606 | 0.000 | 2.13 | ππ* |
| | $S_0 \rightarrow r_7$ | 4.546 | 0.253 | 2.151 | 2.979 | 3.64 | 26.32 | 9.52 | 0.000 | 0.000 | -0.023 | 1.37 | CT |
| | $S_0 \rightarrow r_8$ | 1.974 | 0.701 | 2.992 | -0.458 | 4.98 | 6.68 | 9.47 | -1.282 | 0.114 | 0.001 | 3.50 | ππ* |
| LSDA/aug-cc-pVDZ | $S_0 \rightarrow S_1(r_2)/r_1$ | 4.498 | 0.258 | 2.221 | 2.975 | 3.46 | 25.33 | 9.38 | -0.001 | 0.000 | 0.032 | 1.66 | CT |
| | $S_0 \rightarrow r_5$ | 0.672 | 1.458 | 2.831 | -0.522 | 5.06 | 7.79 | 9.60 | -1.777 | 1.677 | -0.132 | 2.778 | ππ* |
| M05-2X/cc-pVDZ | $S_0 \rightarrow S_1(r_1)/r_5$ | 0.679 | 0.704 | 2.850 | -1.249 | 5.13 | 8.23 | 8.96 | 1.328 | -2.893 | 0.090 | 2.83 | ππ* |
| M05-2X/aug-cc-pVDZ | $S_0 \rightarrow S_1 (r_1)$ | 0.605 | 0.734 | 2.845 | -1.315 | 5.26 | 8.01 | 8.79 | 1.247 | -2.446 | 0.087 | 2.76 | ππ* |
| | $S_0 \rightarrow r_4$ | 1.867 | 0.409 | 2.053 | 0.436 | 6.13 | 25.43 | 10.1 | 0.015 | -0.007 | -0.033 | 3.72 | nπ* |
| M06/cc-pVDZ | $S_0 \rightarrow S_1 (r_1)$ | 0.759 | 0.711 | 2.861 | -0.602 | 5.44 | 8.34 | 9.38 | 2.280 | -0.747 | 0.000 | 2.55 | ππ* |
| | $S_0 \rightarrow r_5/r_6/r_7/r_8$ | 3.141 | 0.385 | 2.521 | 1.367 | 4.58 | 18.71 | 9.26 | 0.072 | 0.073 | -0.033 | 3.60 | CT |
| M06/aug-cc-pVDZ | $S_0 \rightarrow S_1 (r_1)$ | 0.776 | 0.716 | 2.903 | -0.860 | 5.37 | 7.96 | 9.07 | 2.324 | -0.776 | 0.000 | 2.48 | ππ* |
| | $S_0 \rightarrow r_5/r_6$ | 3.193 | 0.433 | 2.551 | 1.320 | 4.52 | 16.57 | 9.13 | 0.374 | 0.283 | -0.029 | 3.72 | nπ* |
| M06-HF/cc-pVDZ | $S_0 \rightarrow S_1 (r_1)$ | 0.759 | 0.711 | 2.861 | -0.602 | 5.44 | 8.34 | 9.38 | 2.280 | -0.747 | 0.000 | 2.55 | ππ* |
| | $S_0 \rightarrow r_5$ | 3.130 | 0.378 | 2.519 | 1.363 | 4.59 | 18.84 | 9.30 | 0.000 | 0.000 | -0.032 | 3.60 | nπ* |
| M06L/cc-pVDZ | $S_0 \rightarrow S_1 (r_1)/r_6$ | 4.609 | 0.241 | 2.134 | 3.141 | 3.39 | 26.19 | 9.44 | 0.002 | -0.002 | -0.024 | 1.65 | CT |
| | $S_0 \rightarrow r_5$ | 3.197 | 0.363 | 2.557 | 1.506 | 4.39 | 18.38 | 9.41 | 0.002 | -0.001 | -0.017 | 2.59 | CT |
| M06L/aug-cc-pVDZ | $S_0 \rightarrow S_1 (r_1)$ | 0.977 | 0.666 | 2.961 | -1.017 | 5.03 | 9.00 | 9.32 | 0.912 | -1.893 | 0.063 | 2.26 | ππ* |
| | $S_0 \rightarrow r_6$ | 4.567 | 0.242 | 2.196 | 3.056 | 3.43 | 25.29 | 9.33 | 0.002 | 0.004 | -0.031 | 1.90 | CT |
| | $S_0 \rightarrow r_7$ | 1.677 | 0.693 | 3.023 | -0.301 | 4.69 | 6.95 | 9.25 | -0.691 | 1.006 | -0.042 | 3.62 | ππ* |
| M11L/cc-pVDZ | $S_0 \rightarrow S_1 (r_1)$ | 0.924 | 0.669 | 2.914 | -1.027 | 5.10 | 8.73 | 9.20 | 0.872 | -1.871 | 0.061 | 2.36 | ππ* |
| | $S_0 \rightarrow r_6$ | 1.423 | 0.701 | 2.708 | -0.474 | 5.25 | 7.68 | 9.29 | 1.835 | -1.611 | 0.137 | 3.01 | ππ* |
| | $S_0 \rightarrow r_7$ | 1.501 | 0.700 | 3.003 | -0.446 | 4.77 | 6.86 | 9.03 | -0.596 | 0.914 | -0.036 | 3.72 | ππ* |
| MN15/cc-pVDZ | $S_0 \rightarrow S_1 (r_1)$ | 0.735 | 0.720 | 2.826 | -1.195 | 5.29 | 8.61 | 9.37 | 1.176 | -2.292 | 0.081 | 2.65 | ππ* |
| | $S_0 \rightarrow r_5/r_6$ | 2.637 | 0.424 | 2.531 | 0.864 | 4.93 | 20.36 | 8.89 | -0.003 | 0.000 | 0.030 | 3.97 | nπ* |
| MN15/aug-cc-pVDZ | $S_0 \rightarrow S_1 (r_1)/r_5$ | 0.735 | 0.720 | 2.826 | -1.195 | 5.29 | 8.62 | 9.37 | 1.176 | -2.292 | 0.081 | 2.65 | ππ* |

| LOT | Excitation | D (Å) | Sr (a.u.) | H (Å) | t (Å) | $E_{coul}$ (eV) | HDI | EDI | x | y | z | EE (eV) | Ex. |
|---|---|---|---|---|---|---|---|---|---|---|---|---|---|
| | $S_0 \rightarrow r_4$ | 2.637 | 0.425 | 2.531 | 0.864 | 4.93 | 20.36 | 8.89 | -0.003 | 0.000 | 0.030 | 3.97 | CT |
| mPW1PW91/cc-pVDZ | $S_0 \rightarrow S_1 (r_1)_{/r_5}$ | 0.869 | 0.696 | 2.855 | -1.077 | 5.22 | 8.88 | 9.39 | 1.011 | -2.087 | 0.070 | 2.55 | ππ* |
| | $S_0 \rightarrow r_6$ | 3.177 | 0.433 | 2.480 | 1.535 | 4.39 | 16.40 | 9.41 | 0.290 | 0.062 | 0.064 | 4.53 | CT |
| | $S_0 \rightarrow r_7$ | 1.099 | 0.709 | 3.046 | -0.872 | 4.85 | 7.35 | 9.21 | -0.454 | 0.890 | -0.025 | 4.17 | ππ* |
| mPWLYP/cc-pVDZ | $S_0 \rightarrow S_1 (r_2)$ | 1.377 | 0.613 | 2.926 | -0.209 | 5.14 | 10.60 | 9.84 | 1.828 | -0.574 | 0.000 | 2.05 | ππ* |
| | $S_0 \rightarrow r_7$ | 1.989 | 0.706 | 2.968 | -0.422 | 5.01 | 6.93 | 9.62 | -1.325 | 0.160 | -0.007 | 3.43 | ππ* |
| | $S_0 \rightarrow r_8$ | 1.020 | 0.465 | 2.591 | -0.959 | 5.85 | 14.81 | 9.00 | 0.000 | 0.000 | -0.079 | 3.28 | nπ* |
| O3LYP/cc-pVDZ | $S_0 \rightarrow S_1 (r_1)$ | 1.051 | 0.660 | 2.899 | -0.938 | 5.12 | 9.50 | 9.57 | 0.913 | -1.897 | 0.063 | 2.34 | ππ* |
| | $S_0 \rightarrow r_7$ | 1.562 | 0.709 | 2.986 | -0.385 | 4.79 | 7.02 | 9.38 | -0.673 | 0.980 | -0.040 | 3.82 | ππ* |
| OLYP/cc-pVDZ | $S_0 \rightarrow S_1 (r_2)$ | 1.301 | 0.622 | 2.923 | -0.722 | 5.02 | 10.35 | 9.73 | 0.867 | -1.726 | 0.061 | 2.12 | ππ* |
| | $S_0 \rightarrow r_7$ | 1.961 | 0.704 | 2.961 | 0.005 | 4.66 | 6.91 | 9.52 | -0.790 | 1.037 | -0.053 | 3.51 | ππ* |
| | $S_0 \rightarrow r_8$ | 1.007 | 0.468 | 2.589 | -0.780 | 5.75 | 14.54 | 8.95 | 0.008 | 0.003 | -0.075 | 3.37 | nπ* |
| OPBE/cc-pVDZ | $S_0 \rightarrow S_1 (r_2)$ | 1.246 | 0.626 | 2.931 | -0.783 | 5.03 | 10.17 | 9.65 | 0.858 | -1.729 | 0.062 | 2.14 | ππ* |
| | $S_0 \rightarrow r_7$ | 1.962 | 0.701 | 2.969 | 0.003 | 4.65 | 6.83 | 9.45 | -0.785 | 1.017 | -0.052 | 3.54 | ππ* |
| | $S_0 \rightarrow r_8$ | 0.991 | 0.470 | 2.603 | -0.808 | 5.73 | 14.18 | 8.89 | 0.007 | 0.004 | -0.074 | 3.38 | nπ* |
| PBE1PBE/cc-pVDZ | $S_0 \rightarrow S_1 (r_1)$ | 0.870 | 0.696 | 2.860 | -0.455 | 5.41 | 8.88 | 9.36 | 2.205 | -0.729 | 0.000 | 2.55 | ππ* |
| | $S_0 \rightarrow r_5/r_6$ | 3.167 | 0.436 | 2.495 | 1.346 | 4.58 | 16.41 | 9.39 | 0.147 | 0.247 | -0.043 | 3.56 | nπ* |
| PW6B95D3/cc-pVDZ | $S_0 \rightarrow S_1 (r_1)$ | 0.857 | 0.698 | 2.854 | -1.093 | 5.22 | 8.92 | 9.41 | 1.028 | -2.107 | 0.070 | 2.57 | ππ* |
| | $S_0 \rightarrow r_6$ | 3.142 | 0.439 | 2.471 | 1.495 | 4.46 | 16.28 | 9.45 | 0.325 | 0.050 | -0.012 | 3.61 | CT |
| | $S_0 \rightarrow r_7$ | 1.028 | 0.711 | 3.043 | -0.956 | 4.88 | 7.52 | 9.23 | -0.418 | 0.874 | -0.023 | 4.21 | ππ* |
| revTPSSh/aug-cc-pVDZ | $S_0 \rightarrow S_1 (r_1)_{/r_5}$ | 0.860 | 0.683 | 2.832 | -0.454 | 5.48 | 8.96 | 9.60 | 2.015 | -0.697 | 0.000 | 2.58 | ππ* |
| | $S_0 \rightarrow r_6$ | 1.483 | 0.690 | 2.604 | -0.614 | 5.62 | 8.27 | 9.75 | -2.454 | -0.168 | 0.000 | 3.23 | ππ* |
| | $S_0 \rightarrow r_7$ | 1.033 | 0.633 | 3.012 | -0.856 | 4.83 | 7.81 | 9.26 | 0.811 | 0.779 | -0.003 | 4.21 | ππ* |
| revTPSS/aug-cc-pVDZ | $S_0 \rightarrow S_1 (r_1)$ | 1.119 | 0.641 | 2.957 | -0.231 | 2.11 | 9.56 | 9.55 | 1.888 | -0.624 | 0.000 | 2.11 | ππ* |
| | $S_0 \rightarrow r_7$ | 1.858 | 0.694 | 3.002 | -0.535 | 5.00 | 6.92 | 9.38 | -1.339 | 0.223 | -0.010 | 3.49 | ππ* |
| | $S_0 \rightarrow r_8$ | 0.947 | 0.475 | 2.617 | -1.035 | 5.81 | 13.90 | 9.04 | -0.011 | 0.005 | -0.088 | 3.53 | nπ* |
| SOGGA11/cc-pVDZ | $S_0 \rightarrow S_1 (r_2)$ | 1.323 | 0.620 | 2.912 | -0.693 | 5.04 | 10.62 | 9.83 | 0.861 | -1.715 | 0.060 | 2.13 | ππ* |
| | $S_0 \rightarrow r_7/r_8$ | 1.015 | 0.469 | 2.574 | -0.756 | 5.78 | 14.93 | 9.07 | 0.015 | -0.003 | -0.073 | 3.22 | nπ* |
| | $S_0 \rightarrow r_9$ | 0.362 | 3.703 | 2.253 | 2.257 | 4.01 | 17.28 | 9.76 | -0.757 | 0.181 | -0.060 | 2.59 | CT |
| SOGGA11x/cc-pVDZ | $S_0 \rightarrow S_1 (r_1)$ | 0.711 | 0.723 | 2.833 | -1.206 | 5.27 | 8.21 | 9.07 | 1.151 | -2.297 | 0.080 | 2.75 | ππ* |
| | $S_0 \rightarrow r_5$ | 3.063 | 0.453 | 2.446 | 1.433 | 4.49 | 15.14 | 9.21 | 0.490 | -0.041 | 0.053 | 4.17 | CT |
| tHCTHhyb/cc-pVDZ | $S_0 \rightarrow S_1 (r_1)_{/r_6}$ | 1.007 | 0.670 | 2.897 | -0.336 | 5.31 | 9.30 | 9.53 | 2.046 | -0.687 | 0.000 | 2.37 | ππ* |
| | $S_0 \rightarrow r_7$ | 1.428 | 0.709 | 3.000 | -0.810 | 5.13 | 7.06 | 9.39 | -1.138 | 0.228 | -0.005 | 3.88 | ππ* |
| TPSSh/cc-pVDZ | $S_0 \rightarrow S_1 (r_1)$ | 1.110 | 0.655 | 2.894 | -0.311 | 5.28 | 9.70 | 9.62 | 1.994 | -0.655 | 0.000 | 2.32 | ππ* |
| | $S_0 \rightarrow r_7$ | 1.586 | 0.710 | 2.967 | -0.726 | 5.14 | 6.97 | 9.45 | -1.207 | 0.212 | -0.005 | 3.83 | ππ* |
| | $S_0 \rightarrow r_8$ | 1.108 | 0.471 | 2.580 | -0.827 | 5.83 | 14.14 | 9.15 | 0.000 | 0.000 | -0.081 | 3.82 | nπ* |
| TPSS/cc-pVDZ | $S_0 \rightarrow S_1 (r_2)$ | 4.586 | 0.249 | 2.143 | 3.035 | 3.60 | 26.45 | 9.61 | 0.000 | 0.000 | -0.023 | 1.47 | CT |
| | $S_0 \rightarrow r_5/r_6$ | 3.247 | 0.356 | 2.543 | 1.479 | 4.48 | 19.28 | 9.59 | 0.000 | 0.000 | 0.012 | 2.34 | nπ* |
| TPSS/aug-cc-pVDZ | $S_0 \rightarrow S_1 (r_1)$ | 1.112 | 0.642 | 2.962 | -0.227 | 5.17 | 9.52 | 9.56 | 1.874 | -0.632 | 0.000 | 2.10 | ππ* |
| | $S_0 \rightarrow r_7$ | 1.865 | 0.693 | 3.007 | -0.532 | 4.99 | 6.94 | 9.39 | -1.334 | 0.220 | -0.007 | 3.46 | ππ* |
| | $S_0 \rightarrow r_8$ | 0.989 | 0.473 | 2.615 | -0.993 | 5.80 | 14.07 | 9.00 | 0.006 | -0.004 | -0.088 | 3.48 | nπ* |
| VSXC/cc-pVDZ | $S_0 \rightarrow S_1 (r_2)_{/r_6}$ | 4.642 | 0.241 | 2.109 | 3.194 | - | 26.84 | 9.57 | 0.002 | 0.000 | -0.022 | 1.51 | CT |
| | $S_0 \rightarrow r_5$ | 3.285 | 0.349 | 2.535 | 1.640 | 4.31 | 19.46 | 9.56 | -0.004 | 0.000 | 0.011 | 2.40 | CT |
| VSXC/aug-cc-pVDZ | $S_0 \rightarrow S_1 (r_1)$ | 1.057 | 0.651 | 2.969 | -0.967 | 5.00 | 9.38 | 9.52 | 0.912 | -1.857 | 0.062 | 2.15 | ππ* |
| | $S_0 \rightarrow r_7$ | 1.760 | 0.696 | 3.027 | -0.232 | 4.66 | 6.93 | 9.36 | -0.741 | 1.085 | -0.050 | 3.51 | ππ* |
| | $S_0 \rightarrow r_8$ | 1.003 | 0.476 | 2.622 | -0.813 | 5.68 | 14.25 | 8.86 | -0.048 | 0.116 | -0.092 | 3.60 | nπ* |
| wB97XD/cc-pVDZ | $S_0 \rightarrow S_1 (r_1)_{/r_5}$ | 0.682 | 0.730 | 2.761 | -1.200 | 5.41 | 8.49 | 9.46 | 1.229 | -2.367 | 0.085 | 2.76 | ππ* |
| | $S_0 \rightarrow r_6$ | 2.656 | 0.500 | 2.547 | 0.925 | 4.79 | 15.08 | 9.29 | -0.374 | 0.011 | -0.019 | 4.28 | nπ* |
| X3LYP/cc-pVDZ | $S_0 \rightarrow S_1 (r_1)$ | 0.916 | 0.688 | 2.860 | -1.038 | 5.20 | 9.08 | 9.48 | 0.993 | -2.049 | 0.069 | 2.48 | ππ* |
| | $S_0 \rightarrow r_6$ | 3.135 | 0.448 | 2.513 | 1.457 | 4.45 | 16.08 | 9.47 | -0.249 | -0.090 | 0.025 | 3.44 | CT |
| | $S_0 \rightarrow r_7$ | 1.195 | 0.713 | 3.026 | -0.764 | 4.85 | 7.25 | 9.28 | -0.523 | 0.932 | -0.031 | 4.05 | ππ* |

**Table S8** Hole-Electron distribution surfaces for the resonant states of each functional. The hole surfaces are depicted in blue colour while green is the position of the electron typically overlapping with the π* MO of lumiflavin. All figures were produced with the Multiwfn program.[76] Three main types of states can be clearly distinguished: (i) ππ* (*i.e.* B3LYP and MPWLYP $r_7$ states) (ii) intramolecular “Charge transfer” type (*i.e.* APFD, LC-wHPBE) and (iii) nπ* (*i.e.* BPBE and OPBE $r_8$ states).

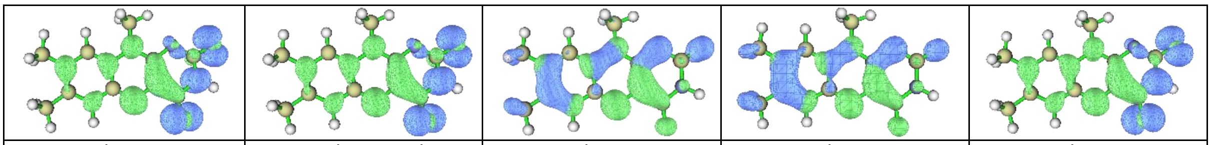

| | | | | |
|---|---|---|---|---|
| APFD/cc-pVDZ_r5 | B1B95/cc-pVDZ_r5/r6 | B1B95/cc-pVDZ_r7 | B3LYP/cc-pVDZ_r7 | B3LYP/cc-pVQZ_r6 |

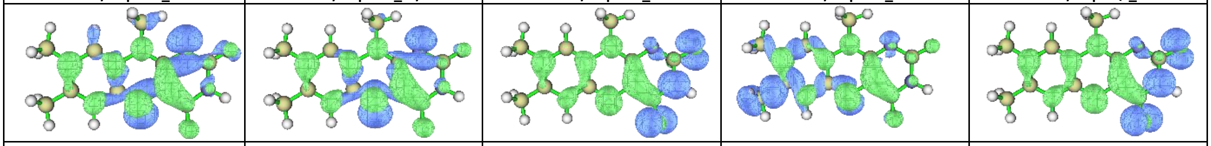

| | | | | |
|---|---|---|---|---|
| B3LYP/cc-pVTZ_r7 | B3LYP/aug-cc-pVDZ_r5 | B3LYP/aug-cc-pVDZ_r6/r7 | B3LYP/aug-cc-pVTZ_r5 | B3LYP/aug-cc-pVTZ_r6 |

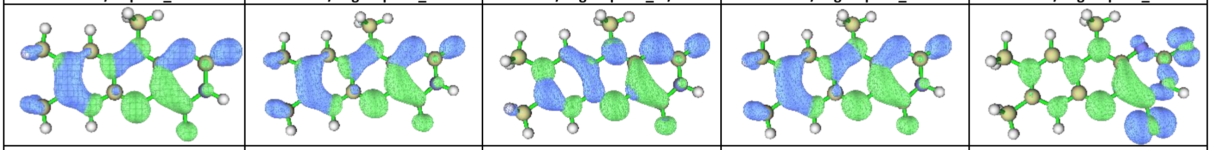

| | | | | |
|---|---|---|---|---|
| B3LYP/aug-cc-pVTZ_r7 | B3P86/cc-pVDZ_r7 | B3PW91/cc-pVDZ_r6 | B3PW91/cc-pVDZ_r7 | B98/cc-pVDZ_r6 |

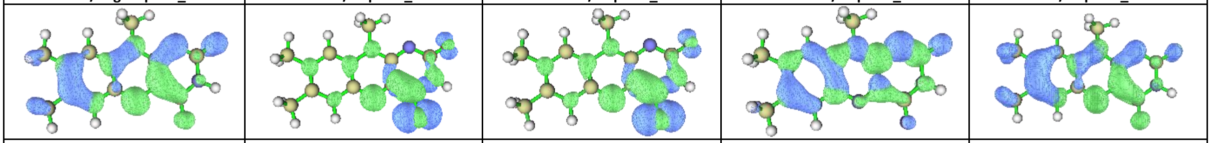

| | | | | |
|---|---|---|---|---|
| B98/cc-pVDZ_r7 | BHHLYP/cc-pVDZ_r4 | BHHLYP/aug-cc-pVDZ_r4 | BHHLYP/aug-cc-pVDZ_r5 | BLYP/cc-pVDZ_r7 |

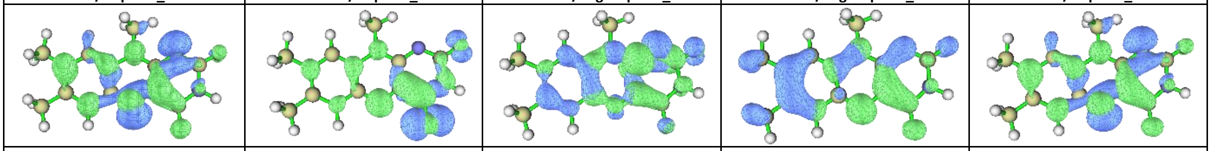

| | | | | |
|---|---|---|---|---|
| BLYP/cc-pVDZ_r8 | BMK/aug-cc-pVDZ_r4 | BMK/aug-cc-pVDZ_r5 | BP86/cc-pVDZ_r7 | BP86/cc-pVDZ_r8 |

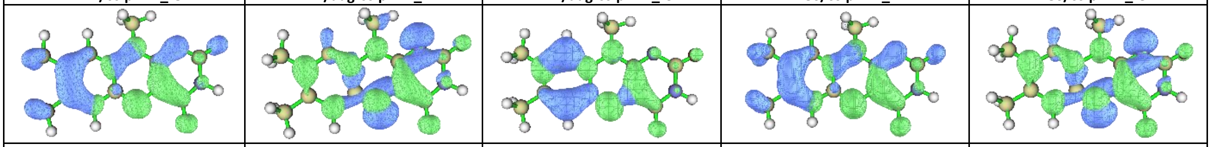

| | | | | |
|---|---|---|---|---|
| BPBE/cc-pVDZ_r7 | BPBE/cc-pVDZ_r8 | CAM-B3LYP/cc-pVDZ_r5 | HCTH/cc-pVDZ_r7 | HCTH/cc-pVDZ_r8 |

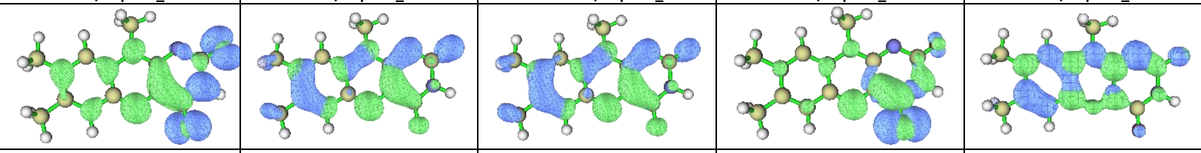

| | | | | |
|---|---|---|---|---|
| HISSBPBE/cc-pVDZ_r6 | HISSBPBE/cc-pVDZ_r7 | HSEH1PBE/cc-pVDZ_r7 | LC-OPBE/cc-pVDZ_r4 | LC-wHPBE/cc-pVDZ_r5 |

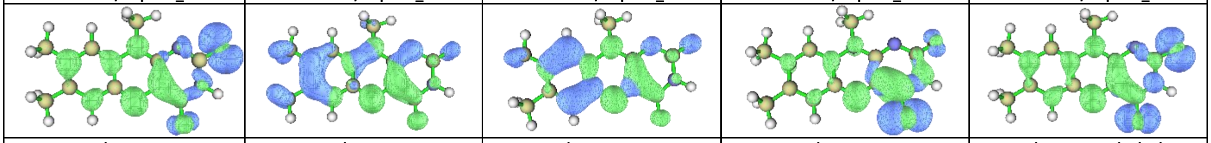

| | | | | |
|---|---|---|---|---|
| LSDA/cc-pVDZ_r7 | LSDA/cc-pVDZ_r8 | LSDA/aug-cc-pVDZ_r5 | M052X/aug-cc-pVDZ_r4 | M06/cc-pVDZ_r5/r7/r7/r8 |

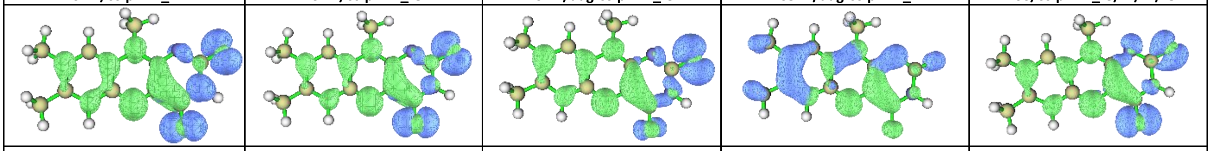

| | | | | |
|---|---|---|---|---|
| M06/aug-cc-pVDZ_r5/r6 | M06-HF/cc-pVDZ_r5 | M06L/aug-cc-pVDZ_r6 | M06L/aug-cc-pVDZ_r7 | M06L/cc-pVDZ_r5 |

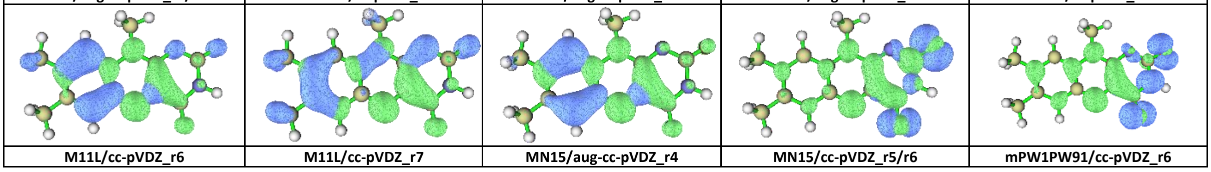

| | | | | |
|---|---|---|---|---|
| M11L/cc-pVDZ_r6 | M11L/cc-pVDZ_r7 | MN15/aug-cc-pVDZ_r4 | MN15/cc-pVDZ_r5/r6 | mPW1PW91/cc-pVDZ_r6 |

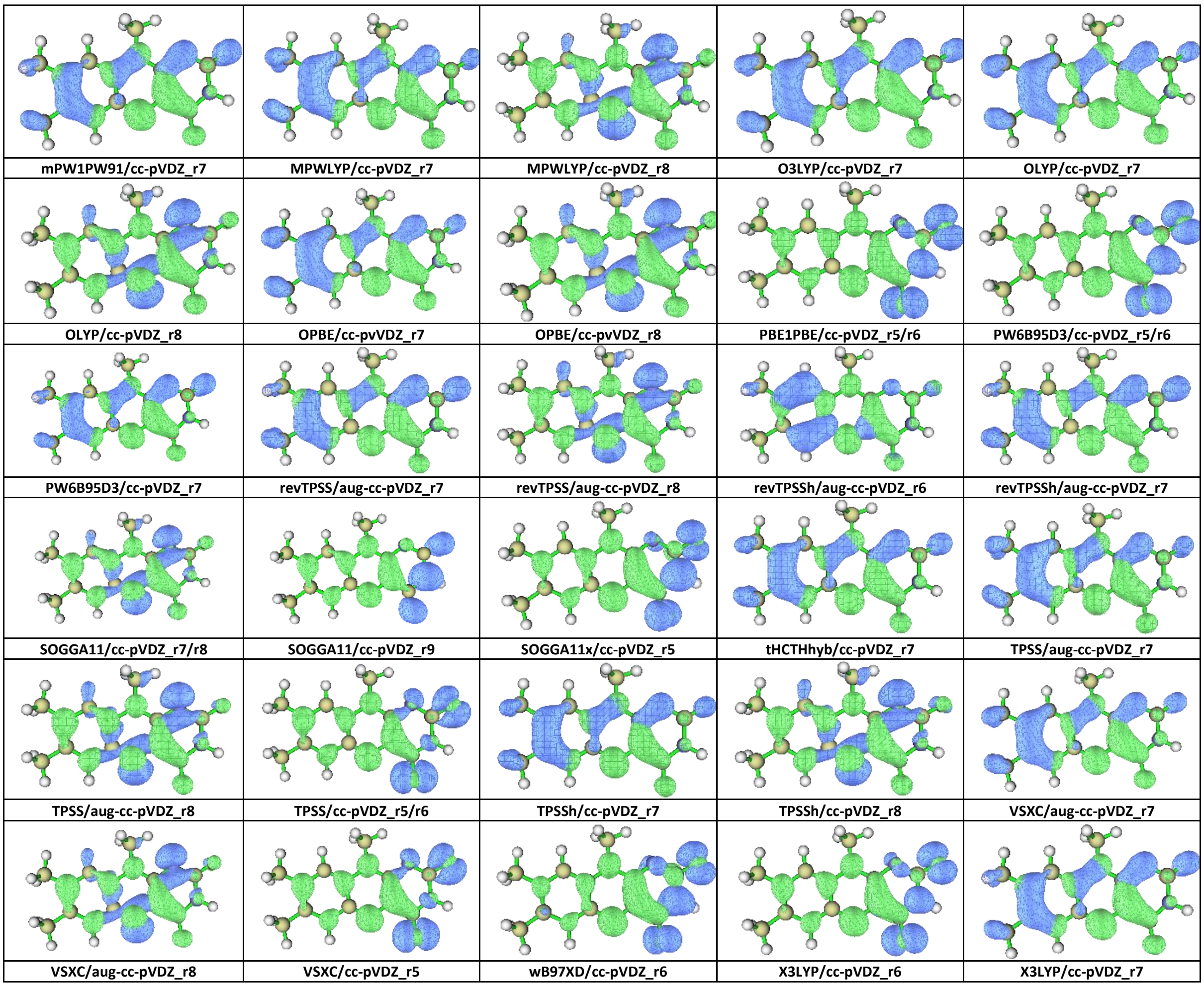


**Table S9** Assignment Tables between the experimental FSRS 3rd EAS of 1FMN* (**Exp. FSRS**)[65, 75] and the calculated first excited singlet state ($S_1$) off-Resonance spectra of each DFT functional (**$S_1$offR**), including vibration number **$v_\#$** and **Assignment** of each vibration to normal modes. The numbering of atoms is taken from **Scheme 1** of the main text. Symmetric and asymmetric C=O stretches are signified as (as) and (**s**).

| Exp. FSRS | APFD/cc-pvdz | | | B1B95/cc-pVDZ | | |
|---|---|---|---|---|---|---|
| | $v_\#$ | $S_1$offR | Assignment | $v_\#$ | $S_1$offR | Assignment |
| 1200 | - | - | - | $v_{51}$ | 1263 | $C_4$-$N_3$, $N_3$-H, $C_2$-$N_3$, $C_{10a}$-$C_{4a}$, $N_1$-$C_2$, $C_9$-H, $C_{9a}$-$C_{5a}$ |
| | $v_{50}$ | 1209 | $C_6$-H, $C_9$-$C_{9a}$, $C_7$-$C_6$, $N_5$-$C_{4a}$, $C_{10a}$-$N_1$, $C_{4a}$-$C_4$ | - | - | - |
| 1250 | $v_{51}$ | 1243 | $C_2$-$N_3$, $N_3$-H, $C_{4a}$-$C_4$, $N_1$-$C_2$, $N_5$-$C_{4a}$, $C_7$-$C_6$, $C_9$-H | $v_{53}$ | 1323 | $N_{10}$-$C_{10a}$, $C_{11}$-$H_3$, $N_5$-$C_{4a}$, $C_8$-$C_7$, $C_6$-$C_{5a}$, $N_5$-$C_{4a}$ |
| 1338 | - | - | - | - | - | - |
| | $v_{56}$ | 1368 | $C_{7a}$-$H_3$, $N_3$-H, $C_{5a}$-$N_5$, $C_{10a}$-$N_1$ | $v_{56}$ | 1378 | $N_3$-H, $C_{5a}$-$N_5$, $C_{7a}$-$H_3$, $C_{10a}$-$N_1$, $C_4$-$N_3$ |
| 1381 | $v_{58}$ | 1388 | $C_{11}$-$H_3$, $N_3$-H, $C_6$-H, $C_4$-$N_3$, $C_7$-$C_6$, $C_6$-$C_{5a}$, $C_{9a}$-$N_{10}$ | - | - | - |
| | $v_{59}$ | 1400 | $C_{8a,11}$-$H_3$, $N_3$-H, $C_9$-$C_7$, $C_6$-$C_{5a}$, $C_{9a}$-$C_{5a}$, $C_4$-$N_3$ | $v_{58}$ | 1395 | $C_{11,7a}$-$H_3$, $N_3$-H, $C_7$-$C_6$, $C_4$-$N_3$, $N_{10}$-$C_{4a}$ |
| | $v_{60}$ | 1416 | $C_{11}$-$H_3$, $C_{9a}$-$N_{10}$, $C_{10a}$-$N_1$, $N_1$-$C_2$, $C_4$-$N_3$ | $v_{59}$ | 1406 | $C_{11,8a}$-$H_3$, $C_8$-$C_7$, $C_{9a}$-$C_{5a}$, $N_3$-H, $C_8$-$C_9$, $C_4$-$N_3$ |
| | $v_{62}$ | 1422 | $N_3$-H, $C_{7a,11}$-$H_3$, $C_6$-H, $C_4$-$N_3$, $C_6$-$C_{5a}$, $C_{9a}$-$C_{5a}$, $C_{10a}$-$C_{4a}$ | - | - | - |
| 1416 | $v_{65}$ | 1448 | $C_{8a}$-$H_3$, $C_9$-H, $C_{10a}$-$C_{4a}$, $N_1$-$C_2$, $C_{11}$-$H_3$ | $v_{63}$ | 1435 | $N_3$-H, $C_6$-H, $C_{9a}$-$C_{5a}$, $C_7$-$C_6$, C1$0_a$-$C_{4a}$, $C_4$-$N_3$, $C_{10a}$-$N_1$ |

| | | | | | | |
|---|---|---|---|---|---|---|
| | - | - | - | $\nu_{65}$ | 1453 | $C_{8a}$-$H_3$, $C_9$-H, $N_1$-$C_2$, $C_{10a}$-$C_{4a}$, $C_{9a}$-$C_{5a}$ |
| 1498 | $\nu_{71}$ | 1539 | $C_8$-$C_7$, $C_{9a}$-$C_{5a}$, $C_{9,6}$-H, $C_6$-$C_{5a}$, $C_9$-$C_{9a}$, $C_{7a}$-$H_3$, $N_5$-$C_{4a}$, $C_{10a}$-$C_{4a}$ | $\nu_{71}$ | 1553 | $C_8$-$C_7$, $C_9$-$C_{9a}$, $C_{9a}$-$C_{5a}$, $C_6$-$C_{5a}$, $C_{6,9}$-H, $C_{7a}$-$H_3$, $N_5$-$C_{4a}$, $C_{10a}$-$C_{4a}$ |
| 1570 | $\nu_{73}$ | 1650 | $C_8$-$C_9$, $C_6$-$C_{5a}$, $C_{9,6}$-H, $C_7$-$C_6$, $C_9$-$C_{9a}$, $N_5$-$C_{4a}$, $C_{10a}$-$N_1$ | $\nu_{73}$ | 1663 | $C_8$-$C_9$, $C_6$-$C_{5a}$, $C_{6,9}$-H, $C_7$-$C_6$, $N_{10}$-$C_{9a}$, $N_5$-$C_{4a}$, $C_{10a}$-$N_1$ |
| | $\nu_{74}$ | 1716 | $C_2$-$O_{2'}$, $N_3$-H, $C_4$-$O_{4'}$, $C_{10a}$-$N_1$ (as) | $\nu_{74}$ | 1742 | $C_2$-$O_{2'}$, $C_4$-$O_{4'}$ (as), $N_3$-H, $C_{4a}$-$C_{10a}$ |
| 1626 | $\nu_{75}$ | 1744 | $C_4$-$O_{4'}$, $N_3$-H, $C_2$-$O_{2'}$, $C_{4a}$-$C_4$ (s) | $\nu_{75}$ | 1763 | $C_4$-$O_{4'}$, $C_2$-$O_{2'}$ (s), $N_3$-H, $C_{4a}$-$C_{10a}$ |
| **Exp. RR FMN $S_1$** | **B3LYP/cc-pVDZ** | | | **B3LYP/aug-cc-pVDZ** | | |
| | **ν#** | **$S_1$offR** | **Assignment** | **ν#** | **$S_1$offR** | **Assignment** |
| 1200 | $\nu_{49}$ | 1191 | $C_{6,9}$-H, $N_3$-H, $N_5$-$C_{5a}$, $C_6$-$C_7$ | $\nu_{49}$ | 1193 | $C_{6,9}$-H, $C_{11}$-$H_3$, $N_3$-H, $N_3$-$C_4$, $C_{5a}$-$N_5$ |
| | - | - | - | - | - | - |
| 1250 | $\nu_{51}$ | 1226 | $C_{6,9}$-H, $N_3$-H, $N_3$-$C_2$, $C_{10a}$-$C_{4a}$, $C_{4a}$-$C_4$ | $\nu_{51}$ | 1223 | $N_3$-H, $N_3$-$C_2$, $C_{10a}$-$C_{4a}$, $C_{4a}$-$C_4$, $C_6$-H |
| 1338 | $\nu_{54}$ | 1316 | $C_{6,9}$-H, $N_5$-$C_{5a}$, $C_{10a}$-$N_1$, $C_{9a}$-$N_{10}$ | $\nu_{54}$ | 1317 | $C_{6,9}$-H, $N_3$-H, $N_3$-$C_2$, $C_{10a}$-$C_{4a}$, $C_{4a}$-$C_4$ |
| | - | - | - | $\nu_{55}$ | 1339 | $C_{6,9}$-H, $C_{5a}$-$N_5$ $N_{10}$-C10a, $C_2$-$N_3$ |
| 1381 | $\nu_{57}$ | 1379 | $C_6$-H, $N_3$-H, $C_{11}H_3$, $C_{9a}$-$N_{10}$, $C_{10a}$-$C_{4a}$ | $\nu_{57}$ | 1376 | $C_6$-H, $N_3$-H, $C_{11}H_3$, $C_6$-$C_{5a}$ |
| | $\nu_{58}$ | 1381 | $C_{7a,8a}$-$H_3$, $N_3$-H, $C_8$-$C_7$, $C_{9a}$-$C_{5a}$ | $\nu_{58}$ | 1382 | $C_{7a,8a}$-$H_3$, $C_{9a}$-$C_{5a}$ |
| | $\nu_{59}$ | 1391 | $C_{7a,8a}$-$H_3$, $C_{9a}$-$N_{10}$, $N_1$-$C_2$, $C_{9a}$-$C_{5a}$ | $\nu_{59}$ | 1394 | $C_{8a}$-$H_3$, $N_3$-H, $N_5$-$C_{4a}$, $N_{10}$-$C_{10a}$ |
| | $\nu_{61}$ | 1405 | $N_3$-H, $C_6$-H, $C_{9a}$-$C_{5a}$, $N_3$-$C_4$, $C_{10a}$-$C_{4a}$ | - | - | - |
| 1416 | $\nu_{64}$ | 1439 | $C_{7a,8a}$-$H_3$, $C_{9a}$-$N_{10}$, $N_1$-$C_2$, $C_{9a}$-$C_{5a}$ | $\nu_{64}$ | 1436 | $N_5$-$C_{4a}$, $C_{7a}$-$H_3$, $C_{11}$-$H_3$, $N_3$-H |
| | $\nu_{65}$ | 1444 | $N_5$-$C_{4a}$, $C_{7a}$-$H_3$, $N_3$-H, $N_4$-$C_3$, $C_{10a}$-$N_1$ | - | - | - |
| 1498 | $\nu_{71}$ | 1524 | $C_9$-H, $C_{9a}$-$N_{10}$, $C_8$-$C_7$, $C_{5a}$-$C_{9a}$, $C_{10a}$-$C_{4a}$ | $\nu_{71}$ | 1514 | $C_{7a}$-$H_3$, $C_{9a}$-$C_{5a}$, $C_8$-$C_7$, $C_{10a}$-$C_{4a}$ |
| 1570 | $\nu_{73}$ | 1630 | $C_{6,9}$-H, $C_8$-$C_9$, $C_{5a}$-$C_6$, $C_{10a}$-$N_1$ | $\nu_{73}$ | 1625 | $C_{6,9}$-H, $C_8$-$C_9$, $C_{5a}$-$C_6$, $N_3$-H, $C_2$-$O_{2'}$, $C_4$-$O_{4'}$ (as) |
| | $\nu_{74}$ | 1692 | $C_2$-$O_{2'}$, $N_3$-H, $C_4$-$O_{4'}$ (as) | $\nu_{74}$ | 1650 | $C_2$-$O_{2'}$, $N_3$-H, $C_{6,9}$-H, $C_8$-$C_9$, $C_{5a}$-$C_6$ (as) |
| 1626 | $\nu_{75}$ | 1719 | $C_4$-$O_{4'}$, $N_3$-H, $C_2$-$O_{2'}$ (s) | $\nu_{75}$ | 1663 | $C_4$-$O_{4'}$, $N_3$-H, $C_2$-$O_{2''}$, $C_{5a}$-$C_6$ (s) |
| **Exp. RR FMN $S_1$** | **B3LYP/cc-pVTZ** | | | **B3LYP/aug-cc-pVTZ** | | |
| | **ν#** | **$S_1$offR** | **Assignment** | **ν#** | **$S_1$offR** | **Assignment** |
| 1200 | $\nu_{49}$ | 1194 | $C_{6,9}$-H, $C_{11}$-$H_3$, $N_3$-H, $N_3$-$C_4$, $C_2$-$N_1$ | $\nu_{49}$ | 1193 | $C_{6,9}$-H, $N_3$-H, $N_3$-$C_4$, $C_{11}$-$H_3$, $C_{5a}$-$N_5$ |
| | $\nu_{51}$ | 1213 | $C_9$-H, $N_3$-$C_2$, $C_{10a}$-$C_{4a}$, $C_{4a}$-$C_4$, $C_{11}$-$H_3$ | $\nu_{51}$ | 1211 | $C_9$-H, $N_3$-$C_2$, $C_{10a}$-$C_{4a}$, $C_{4a}$-$C_4$, $C_{11}$-$H_3$ |
| 1250 | $\nu_{52}$ | 1284 | $C_9$-H, $C_8$-$C_7$, $C_{7a}$-$H_3$ | $\nu_{52}$ | 1285 | $C_9$-H, $C_{5a}$-$N_5$, $C_8$-$C_7$, $C_{7a}$-$H_3$ |
| 1338 | $\nu_{54}$ | 1317 | $C_{6,9}$-H, $N_5$-$C_{5a}$, $C_{10a}$-$N_{10}$ | $\nu_{54}$ | 1316 | $C_{6,9}$-H, $N_5$-$C_{5a}$, $C_9$-$C_{9a}$, $C_{10a}$-$N_{10}$, $N_3$-H |
| | - | - | - | $\nu_{55}$ | 1337 | $C_{6,9}$-H, $N_3$-H, $N_5$-$C_{5a}$, $N_{10}$-$C_{9a}$, $C_{10a}$-$N_1$ |
| 1381 | $\nu_{57}$ | 1381 | $C_6$-H, $N_3$-H, $C_{11}H_3$, $C_6$-$C_{5a}$, $C_{10a}$-$C_{4a}$ | $\nu_{57}$ | 1378 | $C_6$-H, $N_3$-H, $C_{11}H_3$, $C_6$-$C_{5a}$, $C_{10a}$-$C_{4a}$ |
| | $\nu_{58}$ | 1391 | $N_3$-H, $N_5$-$C_{4a}$, $C_{10a}$-$N_1$, $C_{9a}$-$N_{10}$ | $\nu_{58}$ | 1392 | $N_3$-H, $N_5$-$C_{4a}$, $C_{10a}$-$N_1$, $C_{9a}$-$N_{10}$ |
| | - | - | - | - | - | - |
| | - | - | - | - | - | - |
| 1416 | $\nu_{62}$ | 1429 | $C_{11}$-$H_3$, $N_5$-$C_{4a}$, $N_3$-H, $C_{8a}$-$H_3$ | $\nu_{62}$ | 1428 | $N_5$-$C_{4a}$, $C_{11}$-$H_3$, $N_3$-H, $N_3$-$C_4$ |
| | $\nu_{63}$ | 1441 | $C_{11}$-$H_3$, $C_{7a}$-$H_3$, $N_5$-$C_{4a}$ | $\nu_{63}$ | 1440 | $C_{11}$-$H_3$, $C_{7a}$-$H_3$, $N_5$-$C_{4a}$ |
| 1498 | $\nu_{71}$ | 1513 | $C_{7a}$-$H_3$, $C_8$-$C_7$, $C_{9a}$-$C_{5a}$, $C_{10a}$-$C_{4a}$, $C_{11}$-$H_3$ | $\nu_{71}$ | 1511 | $C_{7a}$-$H_3$, $C_8$-$C_7$, $C_{9a}$-$C_{5a}$, $C_{10a}$-$C_{4a}$, $C_{11}$-$H_3$ |
| 1570 | $\nu_{73}$ | 1625 | $C_8$-$C_9$, $C_6$-$C_{5a}$, $C_9$-H, $N_5$-$C_{4a}$ | $\nu_{73}$ | 1622 | $C_8$-$C_9$, $C_6$-$C_{5a}$, $C_9$-H, $N_3$-H, $C_2$-$O_{2'}$, $C_4$-$O_{4'}$ (as) |
| | $\nu_{74}$ | 1668 | $C_2$-$O_{2'}$, $N_3$-H, $C_4$-$O_{4'}$ (as) | $\nu_{74}$ | 1651 | $C_2$-$O_{2'}$, $N_3$-H, $C_4$-$O_{4'}$, $C_8$-$C_9$, $C_6$-$C_{5a}$, $C_9$-H (as) |
| 1626 | $\nu_{75}$ | 1686 | $C_4$-$O_{4'}$, $C_2$-$O_{2'}$, $N_3$-H (s) | $\nu_{75}$ | 1664 | $C_4$-$O_{4'}$, $C_2$-$O_{2'}$, $N_3$-H (s) |
| **Exp. RR FMN $S_1$** | **B3LYP/cc-pVQZ** | | | **B3P86/cc-pVDZ** | | |
| | **ν#** | **$S_1$offR** | **Assignment** | **ν#** | **$S_1$offR** | **Assignment** |

| | | | | | | |
|---|---|---|---|---|---|---|
| 1200 | $v_{49}$ | 1194 | $C_{6,9}$-H, $N_3$-H, $N_3$-$C_4$, $C_{11}$-$H_3$, $C_{5a}$-$N_5$ | $v_{51}$ | 1263 | $C_4$-$N_3$, $N_3$-H, $C_2$-$N_3$, $C_{10a}$-$C_{4a}$, $N_1$-$C_2$, $C_9$-H, $C_{9a}$-$C_{5a}$ |
| | $v_{51}$ | 1211 | $C_9$-H, $N_3$-$C_2$, $C_{9a}$-$C_9$, $C_{10a}$-$C_{4a}$, $C_{4a}$-$C_4$, $C_{11}$-$H_3$ | - | - | - |
| 1250 | $v_{52}$ | 1285 | $C_9$-H, $C_{5a}$-$N_5$, $C_8$-$C_7$, $C_{7a}$-$H_3$ | $v_{53}$ | 1323 | $N_{10}$-$C_{10a}$, $C_{11}$-$H_3$, $N_5$-$C_{4a}$, $C_8$-$C_7$, $C_6$-$C_{5a}$, $N_5$-$C_{4a}$ |
| 1338 | $v_{54}$ | 1317 | $C_{6,9}$-H, $N_5$-$C_{5a}$, $C_9$-$C_{9a}$, $C_{10a}$-$N_{10}$, $N_3$-H | - | - | - |
| | - | - | - | $v_{56}$ | 1378 | $N_3$-H, $C_{5a}$-$N_5$, $C_{7a}$-$H_3$, $C_{10a}$-$N_1$, $C_4$-$N_3$ |
| 1381 | $v_{57}$ | 1379 | $C_6$-H, $N_3$-H, $C_{11}H_3$, $C_6$-$C_{5a}$, $C_{10a}$-$C_{4a}$ | - | - | - |
| | $v_{58}$ | 1393 | $N_3$-H, $C_{10a}$-$N_1$, $N_5$-$C_{4a}$, $C_{9a}$-$N_{10}$ | $v_{58}$ | 1395 | $C_{11,7a}$-$H_3$, $N_3$-H, $C_7$-$C_6$, $C_4$-$N_3$, $N_{10}$-$C_{4a}$ |
| | - | - | - | $v_{59}$ | 1406 | $C_{11,8a}$-$H_3$, $C_8$-$C_7$, $C_{9a}$-$C_{5a}$, $N_3$-H, $C_8$-$C_9$, $C_4$-$N_3$ |
| | - | - | - | - | - | - |
| 1416 | $v_{62}$ | 1429 | $N_5$-$C_{4a}$, $C_{11}$-$H_3$, $N_3$-H, $N_3$-$C_4$, $N_1$-$C_2$ | $v_{63}$ | 1435 | $N_3$-H, $C_6$-H, $C_{9a}$-$C_{5a}$, $C_7$-$C_6$, $C_{10a}$-$C_{4a}$, $C_4$-$N_3$, $C_{10a}$-$N_1$ |
| | $v_{63}$ | 1441 | $C_{11}$-$H_3$, $C_{7a}$-$H_3$, $N_5$-$C_{4a}$, $N_3$-H | $v_{65}$ | 1453 | $C_{8a}$-$H_3$, $C_9$-H, $N_1$-$C_2$, $C_{10a}$-$C_{4a}$, $C_{9a}$-$C_{5a}$ |
| 1498 | $v_{71}$ | 1512 | $C_{7a}$-$H_3$, $C_8$-$C_7$, $C_{9a}$-$C_{5a}$, $C_{10a}$-$C_{4a}$, $C_{11}$-$H_3$ | $v_{71}$ | 1553 | $C_8$-$C_7$, $C_9$-$C_{9a}$, $C_{9a}$-$C_{5a}$, $C_6$-$C_{5a}$, $C_{6,9}$-H, $C_{7a}$-$H_3$, $N_5$-$C_{4a}$, $C_{10a}$-$C_{4a}$ |
| 1570 | $v_{73}$ | 1623 | $C_{6,9}$-H, $C_8$-$C_9$, $C_6$-$C_{5a}$, $N_3$-H, $C_2$-$O_{2'}$, $C_4$-$O_{4'}$ (as) | $v_{73}$ | 1663 | $C_8$-$C_9$, $C_6$-$C_{5a}$, $C_{6,9}$-H, $C_7$-$C_6$, $N_{10}$-$C_{9a}$, $N_5$-$C_{4a}$, $C_{10a}$-$N_1$ |
| | $v_{74}$ | 1659 | $C_2$-$O_{2'}$, $N_3$-H, $C_4$-$O_{4'}$, $C_8$-$C_9$, $C_6$-$C_{5a}$, $C_{10a}$-$N_1$ (as) | $v_{74}$ | 1742 | $C_2$-$O_{2'}$, $C_4$-$O_{4'}$ (as), $N_3$-H, $C_{4a}$-$C_{10a}$ |
| 1626 | $v_{75}$ | 1675 | $C_4$-$O_{4'}$, $C_2$-$O_{2'}$, $N_3$-H (s) | $v_{75}$ | 1763 | $C_4$-$O_{4'}$, $C_2$-$O_{2'}$ (s), $N_3$-H, $C_{4a}$-$C_{10a}$ |

| **Exp. RR FMN $S_1$** | **B3PW91/cc-pVDZ** | | | **B98/cc-pVDZ** | | |
|---|---|---|---|---|---|---|
| | **$v_\#$** | **$S_1$offR** | **Assignment** | **$v_\#$** | **$S_1$offR** | **Assignment** |
| 1200 | $v_{49}$ | 1197 | $C_6$-H, $C_8$-$C_7$, $N_3$-H, $C_{5a}$-$N_5$, $C_7$-$C_6$, $C_{10a}$-$N_1$, $C_4$-$N_3$ | $v_{51}$ | 1228 | $C_2$-$N_3$, $C_{4a}$-$C_4$, $N_3$-H, $C_9$-H, $N_{10}$-$C_{10a}$, $C_{5a}$-$N_5$, $C_{5a}$-$C_{9a}$ |
| | $v_{50}$ | 1209 | $C_{6,9}$-H, $C_{11}$-$H_3$, $C_9$-$C_{9a}$, $C_{4a}$-$C_4$, $C_4$-$N_3$, $N_3$-H, $C_7$-$C_6$, $C1_{0a}$-$N_1$ | - | - | - |
| 1250 | $v_{51}$ | 1245 | $C_2$-$N_3$, $C_{4a}$-$C_4$, $N_3$-H, $C_9$-H, $N_1$-$C_2$, $N_5$-$C_{4a}$, $C_{9a}$-$C_{5a}$, $C_7$-$C_6$ | $v_{53}$ | 1294 | $N_{10}$-$C_{10}$a, $C_{9a}$-$C_{5a}$, $C_7$-$C_6$, $C_{11}$-$H_3$, $N_5$-$C_{4a}$, $N_3$-$C_2$, $C_{6,9}$-H |
| 1338 | $v_{53}$ | 1309 | $N_{10}$-$C_{10a}$, $C_6$-$C_{5a}$, $N_5$-$C_{4a}$, $C_8$-$C_7$, $C_{9a}$-$C_{5a}$, $C_2$-$N_3$, $C_{11}$-$H_3$, $C_{6,9}$-H, $C_{8a}$-$H_3$, $N_3$-H | $v_{54}$ | 1314 | $C_{5a}$-$N_5$, $C_{6,9}$-H, $C_9$-$C_{9a}$, $N_{10}$-$C_{10a}$, $C_{10a}$-$C_{4a}$, $C_2$-$N_3$ |
| | $v_{56}$ | 1366 | $C_{7a}$-$H_3$, $N_3$-H, $C_{6,9}$-H, $C_{10a}$-$N_1$, $C_{5a}$-$N_5$ | - | - | - |
| 1381 | $v_{58}$ | 1384 | $C_{11,7a}$-$H_3$, $N_3$-H, $C_6$-H, $C_7$-$C_6$, $C_6$-$C_{5a}$, $C_{5a}$-$C_{9a}$, $C_{9a}$-$N_{10}$, $C_{10a}$-$C_{4a}$ | $v_{57}$ | 1378 | $C_{11,7a}$-$H_3$, $N_3$-H, $C_6$-H, $C_{10a}$-$C_{4a}$, $C_7$-$C_6$, $C_{9a}$-$C_{5a}$, $N_{10}$-$C_{11}$ |
| | $v_{59}$ | 1397 | $C_{8a,11}$-$H_3$, $N_3$-H, $C_9$-H, $C_6$-$C_{5a}$, $C_4$-$N_3$, $C_{9a}$-$C_{5a}$, $C_8$-$C_7$, $C_2$-$N_3$ | $v_{58}$ | 1380 | $C_{7a,8a}$-$H_3$, $C_8$-$C_7$, $C_{9a}$-$C_{5a}$ |
| | $v_{60}$ | 1412 | $C_{11}$-$H_3$, $C_{9a}$-$N_{10}$, $C_{10a}$-$N_1$, $C_{7a,8a}$-$H_3$, $C_{10a}$-$C_{4a}$, $N_5$-$C_{4a}$, $N_1$-$C_2$ | $v_{59}$ | 1390 | $C_{8a}$-$H_3$, $C_{9a}$-$C_{5a}$, $C_{9a}$-$N_{10}$, $C_2$-$N_3$ |
| | $v_{61}$ | 1414 | $N_3$-H, $C_{11}$-$H_3$, $C_6$-H, $C_{9a}$-$C_{5a}$, $C_{10a}$-$C_{4a}$, $C_8$-$C_7$, $C_4$-$N_3$ | $v_{61}$ | 1406 | $N_3$-H, $C_{9a}$-$C_{5a}$, $C_6$-H, $C_4$-$N_3$, $N_5$-$C_{4a}$, $C_{11}$-$H_3$ |
| 1416 | $v_{62}$ | 1418 | $C_{11,7a}$-$H_3$, $C_6$-H, $N_3$-H, $C_4$-$N_3$, $C_7$-$C_6$, $N_{10}$-$C_{10a}$ | $v_{64}$ | 1440 | $N_5$-$C_{4a}$, $C_{7a}$-$H_3$, $C_{4a}$-$C_4$, $C_{10a}$-$N_1$, $N_3$-H, $C_2$-$N_3$, $C_8$-$C_7$ |
| | $v_{65}$ | 1445 | $C_{8a,11,7a}$-$H_3$, $C_9$-H, $C_{10a}$-$N_1$, $C_{10a}$-$C_{4a}$, $C_{9a}$-$C_{5a}$ | $v_{65}$ | 1443 | $C_{7a,8a,11}$-$H_3$, $C_{10a}$-$N_1$, $C_{10a}$-$C_{4a}$, $C_8$-$C_9$ |
| 1498 | $v_{71}$ | 1537 | $C_8$-$C_7$, $C_9$-$C_{9a}$, $C_{9a}$-$C_{5a}$, $C_6$-$C_{5a}$, $C_{6,9}$-H, $C_{7a}$-$H_3$, $N_5$-$C_{4a}$, $C_{10a}$-$C_{4a}$ | $v_{71}$ | 1519 | $C_8$-$C_7$, $C_9$-$C_{9a}$, $C_{9a}$-$C_{5a}$, $C_6$-$C_{5a}$, $C_{6,9}$-H, $C_{7a}$-$H_3$, $N_5$-$C_{4a}$, $C_{10a}$-$C_{4a}$ |
| 1570 | $v_{73}$ | 1645 | $C_8$-$C_9$, $C_6$-$C_{5a}$, $C_{6,9}$-H, $C_7$-$C_6$, $N_{10}$-$C_{9a}$, $N_5$-$C_{4a}$, $C_{10a}$-$N_1$ | $v_{73}$ | 1626 | $C_8$-$C_9$, $C_6$-$C_{5a}$, $C_{6,9}$-H, $C_7$-$C_6$, $N_{10}$-$C_{9a}$, $N_5$-$C_{4a}$, $C_{10a}$-$N_1$ |
| | $v_{74}$ | 1711 | $C_2$-$O_{2'}$, $N_3$-H, $C_4$-$O_{4'}$ (as), $C_{4a}$-$C_{10a}$, $C_4$-$N_3$ | $v_{74}$ | 1709 | $C_2$-$O_{2'}$, $N_3$-H, $C_4$-$O_{4'}$ (as), $C_{4a}$-$C_{10a}$, $C_4$-$N_3$ |

| | | | | | | |
|---|---|---|---|---|---|---|
| 1626 | $\nu_{75}$ | 1739 | $C_4$-$O_{4'}$, $N_3$-H, $C_2$-$O_{2'}$, $C_4$-$C_{4a}$ (s) | $\nu_{75}$ | 1733 | $C_4$-$O_{4'}$, $N_3$-H, $C_2$-$O_{2'}$ (s), $C_{4a}$-$C_{10a}$ |
| **Exp. RR FMN $S_1$** | **BHHLYP/cc-pVDZ** | | | **BHHLYP/aug-cc-pVDZ** | | |
| | **$\nu_\#$** | **$S_1$offR** | **Assignment** | **$\nu_\#$** | **$S_1$offR** | **Assignment** |
| 1200 | $\nu_{46}$ | 1144 | $C_{11}$-$H_3$, $N_3$-H, $C_4$-$C_{4a}$, $C_{9a}$-$C_{5a}$, $C_4$-$N_3$, $C_{9a}$-$N_{10}$, $C_8$-$C_7$ | $\nu_{46}$ | 1146 | $C_{11}$-$H_3$, $N_3$-H, $C_{4a}$-$C_{10a}$, $N_{10}$-$C_{10a}$, $C_{9a}$-$N_{10}$, $C_4$-$N_3$, $C_{8a}$-$H_3$, $C_8$-$C_7$ |
| | - | - | - | - | - | - |
| 1250 | $\nu_{50}$ | 1264 | $C_6$-H, $C_{11,7a}$-$H_3$, $C_9$-$C_{9a}$, $C_7$-$C_6$, $C_{4a}$-$C_4$ | $\nu_{50}$ | 1265 | $C_6$-H, $C_7$-$C_6$, $C_9$-$C_{9a}$, $N_5$-$C_{4a}$, $C_2$-$N_3$, $C_{10a}$-$C_{4a}$ |
| 1338 | $\nu_{54}$ | 1381 | $C_{5a}$-$N_5$, $C_{9a}$-$C_{5a}$, $C_{10a}$-$C_{4a}$, $C_9$-$C_{9a}$, $N_3$-H, $C_9$-H, $N_{10}$-$C_{9a}$ | $\nu_{54}$ | 1374 | $C_{5a}$-$N_5$, $C_9$-$C_{9a}$, $C_{10a}$-$C_{4a}$, $N_3$-H, $C_9$-H, $C_{9a}$-$N_{10}$ |
| | - | - | - | - | - | - |
| 1381 | - | - | - | $\nu_{56}$ | 1438 | $C_{7a,11}$-$H_3$, $C_6$-H, $C_7$-$C_6$, $C_6$-$C_{5a}$, $C_{9a}$-$N_{10}$, $N_{10}$-$C_{10a}$, $C_{10a}$-$N_1$, $C_{5a}$-$N_5$, $C_4$-$N_3$ |
| | $\nu_{57}$ | 1442 | $C_{11}$-$H_3$, $C_6$-H, $C_7$-$C_6$, $C_{9a}$-$N_{10}$, $C_{10a}$-$N_1$, $C_4$-$N_3$ | $\nu_{57}$ | 1442 | $C_{7a,8a}$-$H_3$, $C_8$-$C_7$, $C_9$-$C_{9a}$, $C_6$-$C_{5a}$ |
| | $\nu_{58}$ | 1448 | $C_8$-$H_3$, $C_6$-$C_{5a}$, $C_{9a}$-$C_{5a}$ | $\nu_{58}$ | 1449 | $N_3$-H, $C_{8a,7a}$-$H_3$, $C_4$-$N_3$, $C_6$-$C_{5a}$, $C_{10a}$-$N_1$ |
| | - | - | - | - | - | - |
| 1416 | $\nu_{60}$ | 1477 | $C_{11}$-$H_3$, $C_{7a,8a}$-$H_3$, $C_7$-$C_6$, $N_{10}$-$C_{10a}$ | $\nu_{60}$ | 1482 | $C_{11,7a,8a}$-$H_3$, $C_{9a}$-$N_{10}$, $N_{10}$-$C_{10a}$, $N_3$-H, $N_1$-$C_2$ |
| | $\nu_{64}$ | 1504 | $C_{8a}$-$H_3$, $C_{10a}$-$C_{4a}$, $C_{9a}$-$C_{5a}$, $N_3$-H | $\nu_{63}$ | 1507 | $C_6$-H, $C_{7a}$-$H_3$, $C_{9a}$-$C_{5a}$, $C_8$-$C_7$, $C_{10a}$-$C_{4a}$, $C_{8a,11}$-$H_3$, $C_{10a}$-$N_1$ |
| 1498 | $\nu_{72}$ | 1632 | ($\nu_{71}$, $\nu_{70}$) $C_{10a}$-$N_1$, $N_5$-$C_{4a}$, $C_4$-$N_3$, $C_8$-$C_7$, $C_9$-H, $N_3$-H | $\nu_{72}$ | 1630 | ($\nu_{70}$, $\nu_{71}$) $C_{10a}$-$N_1$, $C_{4a}$-$N_3$, $N_5$-$C_{4a}$, $C_8$-$C_9$, $N_3$-H, $C_9$-H |
| 1570 | $\nu_{73}$ | 1696 | $C_8$-$C_9$, $C_6$-$C_{5a}$, $C_{9a}$-$N_{10}$, $C_7$-$C_6$, $C_9$-H, $C_{10a}$-$N_1$, $N_5$-$C_{4a}$ | $\nu_{73}$ | 1693 | $C_8$-$C_9$, $C_6$-$C_{5a}$, $C_{9a}$-$N_{10}$, $C_7$-$C_6$, $C_9$-H, $C_{10a}$-$N_1$, $N_5$-$C_{4a}$ |
| | $\nu_{74}$ | 1801 | $C_4$-$O_{4'}$, $N_3$-H, $C_2$-$O_{2'}$ (as) | $\nu_{74}$ | 1742 | $N_3$-H, $C_2$-$O_{2'}$, $C_4$-$O_{4'}$(as) |
| 1626 | $\nu_{75}$ | 1822 | $C_2$-$O_{2'}$, $C_4$-$O_{4'}$, $N_3$-H, $N_{10}$-$C_{10a}$ (s) | $\nu_{75}$ | 1765 | $C_2$-$O_{2'}$, $C_4$-$O_{4'}$, $N_3$-H, $C_{10a}$-$N_1$ (s) |
| **Exp. RR FMN $S_1$** | **BLYP/cc-pVDZ** | | | **BMK/cc-pVDZ** | | |
| | **$\nu_\#$** | **$S_1$offR** | **Assignment** | **$\nu_\#$** | **$S_1$offR** | **Assignment** |
| 1200 | $\nu_{51}$ | 1168 | $C_2$-$N_3$, $C_{10a}$-$C_{4a}$, $C_{6,9}$-H, $N_{10}$-$C_{4a}$, $C_9$-$C_{9a}$, $C_{9a}$-$C_{5a}$ | $\nu_{50}$ | 1229 | $C_6$-H, $C_{11}$-$H_3$, $C_4$-$N_3$, $N_{10}$-$C_{10a}$, $C_9$-$C_{9a}$, $C_{4a}$-$C_4$, $C_7$-$C_6$, $C_{9a}$-$C_{5a}$ |
| | $\nu_{52}$ | 1219 | $C_{6,9}$-H, $C_8$-$C_7$, $C_6$-$C_{5a}$, $C_4$-$N_3$, $N_3$-H | - | - | - |
| 1250 | $\nu_{53}$ | 1238 | $C_{6,9}$-H, $N_3$-H, $C_{11}$-$H_3$, $N_{10}$-$C_{10a}$, $N_5$-$C_{4a}$, $C_8$-$C_9$ | $\nu_{51}$ | 1267 | $N_3$-H, $C_2$-$N_3$, $C_{4a}$-$C_4$, $C_4$-$N_3$, $N_5$-$C_{4a}$, $C_6$-H, $C_7$-$C_6$ |
| 1338 | $\nu_{54}$ | 1258 | $C_{6,9}$-H, $C_{9a}$-$C_{5a}$, $C_{10a}$-$C_{4a}$, $N_5$-$C_{4a}$ | $\nu_{53}$ | 1321 | $C_9$-H, $C_8$-$C_7$, $C_6$-$C_{5a}$, $C_8$-$C_9$, $C_{5a}$-$N_5$, $C_{7a,8a,11}$-$H_3$ |
| | $\nu_{55}$ | 1284 | $C_{11}$-$H_3$, $N_3$-H, $N_{10}$-$C_{10a}$, $C_{9a}$-$N_{10}$, $C_7$-$C_6$, $C_{9a}$-$C_{5a}$ | - | - | - |
| 1381 | $\nu_{56}$ | 1311 | $C_{5a}$-$C_{9a}$, $C_{4a}$-$C_4$, $C_8$-$C_9$, $N_1$-$C_2$, $C_{9a}$-$N_{10}$, $N_5$-$C_{4a}$ | $\nu_{56}$ | 1391 | $C_{11,8a,7a}$-$H_3$, $N_3$-H, $C_8$-$C_7$, $C_{5a}$-$N_5$, $C_7$-$C_6$, $C_{9a}$-$C_{5a}$, $C_{10a}$-$N_1$ |
| | $\nu_{59}$ | 1340 | $C_{7a}$-$H_3$, $N_3$-H, $N_5$-$C_{4a}$, $C_4$-$N_3$ | $\nu_{57}$ | 1395 | $C_{8a,7a,11}$-$H_3$, $N_3$-H, $C_9$-H, $C_{5a}$-$N_5$, $C_{9a}$-$N_{10}$, $C_{10a}$-$C_{4a}$, $C_{10a}$-$N_1$ |
| | $\nu_{60}$ | 1343 | $N_3$-H, $C_{9a}$-$C_{5a}$, $C_6$-H, $C_{8a}$-$H_3$, $C_8$-$C_7$, $C_4$-$N_3$, $N_5$-$C_{4a}$ | $\nu_{58}$ | 1401 | $C_{8a,7a}$-$H_3$, $N_3$-H, $C_6$-$C_{5a}$, $C_8$-$C_7$, $C_8$-$C_9$ |
| | - | - | - | $\nu_{59}$ | 1406 | $N_3$-H, $C_{11}$-$H_3$, $C_6$-H, $C_{10a}$-$C_{4a}$, $C_2$-$N_3$, $C_{9a}$-$C_{5a}$, $C_4$-$N_3$ |
| 1416 | $\nu_{62}$ | 1366 | $C_{11,8a}$-$H_3$, $C_{4a}$-$C_4$, $N_1$-$C_2$ | $\nu_{61}$ | 1437 | $N_3$-H, $C_{7a,8a,11}$-$H_3$, $C_{9a}$-$N_{10}$, $N_5$-$C_{4a}$, $N_1$-$C_2$, $C_8$-$C_7$ |
| | $\nu_{65}$ | 1397 | $C_{11,7a,8a}$-$H_3$, $N_{10}$-$C_{10a}$, $C_9$-$C_{9a}$ | $\nu_{65}$ | 1465 | $C_{7a,8a}$-$H_3$, $N_3$-H, $C_{5a}$-$N_5$, $C_{4a}$-$C_4$, $N_1$-$C_2$, $C_2$-$N_3$ |

| | | | | | | |
|---|---|---|---|---|---|---|
| 1498 | $\nu_{71}$ | 1475 | $C_{6,9}-H, C_8-C_7, C_9-C_{9a}, C_6-C_{5a}$ | $\nu_{71}$ | 1544 | ($\nu_{72}$) $C_8-C_7, C_{9a}-C_{5a}, C_{7a,11}-H_3, C_9-H, C_{9a}-N_{10}, C_{10a}-C_{4a}, C_{10a}-N_1$ |
| 1570 | $\nu_{73}$ | 1554 | $C_8-C_9, C_6-C_{5a}, C_2-O_{2'}, C_{6,9}-H, N_3-H$ | $\nu_{73}$ | 1648 | $C_9-H, C_8-C_9, C_6-C_{5a}, C_{9a}-N_{10}, C_7-C_6, C_{10a}-N_1, N_5-C_{4a}$ |
| | $\nu_{74}$ | 1568 | $C_2-O_{2'}, N_3-H, C_{10a}-C_{4a}, C_6-C_{5a}, C_8-C_9, C_9-H$ ~(as) | $\nu_{74}$ | 1786 | $C_2-O_{2'}, C_4-O_{4'}, N_3-H$ (as) |
| 1626 | $\nu_{75}$ | 1643 | $C_4-O_{4'}, N_3-H, C_{4a}-C_4$ ~(s) | $\nu_{75}$ | 1810 | $C_2-O_{2'}, C_4-O_{4'}$ (s) |
| **Exp. RR FMN $S_1$** | **BMK/aug-cc-pVDZ** | | | **BP86/cc-pVDZ** | | |
| | **$\nu_\#$** | **$S_1$offR** | **Assignment** | **$\nu_\#$** | **$S_1$offR** | **Assignment** |
| 1200 | $\nu_{50}$ | 1227 | $C_6-H, C_7-C_6, C_9-C_{9a}, C_{10a}-C_{4a}, C_{11,7a,8a}-H_3$ | $\nu_{48}$ | 1118 | $C_6-H, C_7-C_6, C_{5a}-N_5, N_5-C_{4a}, C_{10a}-N_1, C_4-N_3$ |
| | - | - | - | $\nu_{51}$ | 1184 | $C_2-N_3, C_{4a}-C_4, C_{9,6}-H, N_3-H$ |
| 1250 | $\nu_{51}$ | 1260 | $C_2-N_3, C_4-N_3, N_3-H, C_{6,9}-H, C_{4a}-C_4, N_1-C_2, C_7-C_6$ | $\nu_{53}$ | 1241 | $C_{6,9}-H, N_3-H, C_{11}-H_3, C_{10a}-N_1, N_5-C_{4a}$ |
| 1338 | $\nu_{54}$ | 1333 | $C_8-C_7, C_{9a}-C_{5a}, C_{5a}-N_5, N_3-H, N_{10}-C_{10a}, C_{10a}-C_{4a}, C_4-N_3, N_1-C_2$ | - | - | - |
| | $\nu_{55}$ | 1378 | $C_{7a}-H_3, C_9-H, C_8-C_7, C_6-C_{5a}, N_3-H, C_4-N_3$ | $\nu_{55}$ | 1297 | $N_3-H, N_{10}-C_{10a}, C_7-C_6, C_6-C_{5a}, C_4-N_3, N_1-C_2$ |
| 1381 | $\nu_{56}$ | 1390 | $N_3-H, C_{7a,11,8a}-H_3, C_{5a}-N_5, C_{10a}-N_1, C_4-N_3, C_7-C_6, C_{9a}-N_{10}$ | $\nu_{57}$ | 1326 | $N_3-H, C_{11,7a}-H_3, C_6-C_{5a}, C_{9a}-N_{10}, C_2-N_1$ |
| | $\nu_{57}$ | 1396 | $C_{7a,8a}-H_3, N_3-H, C_6-C_{5a}, C_8-C_7$ | - | - | - |
| | $\nu_{58}$ | 1401 | $N_3-H, C_{8a}-H_3, C_{10a}-C_{4a}, C_{9a}-C_{5a}, C_2-N_3$ | $\nu_{60}$ | 1346 | $N_3-H, C_{11}-H_3, C_6-H, N_5-C_{4a}, N_{10}-C_{10a}$ |
| | $\nu_{59}$ | 1406 | $C_{8a}-H_3, N_3-H, C_6-H, C_4-N_3, C_{10a}-C_{4a}$ | $\nu_{61}$ | 1356 | $C_8-C_7, C_{9a}-C_{5a}, C_9-H, C_{8a,11}-H_3, N_3-H$ |
| 1416 | $\nu_{61}$ | 1441 | $C_{11}-H_3, C_6-H, N_3-H, C_{9a}-C_{5a}, C_{10a}-C_{4a}, N_{10}-C_{10a}, N_5-C_{4a}$ | $\nu_{64}$ | 1391 | $C_{7a,8a,11}-H_3, N_5-C_{4a}, C_{4a}-C_4$ |
| | $\nu_{64}$ | 1463 | $N_5-C_{4a}, C_{8a,7a}-H_3, N_3-H, C_{10a}-N_1, C_2-N_3$ | $\nu_{65}$ | 1391 | $C_{8a,7a,11}-H_3, C_{10a}-N_1, C_{5a}-N_5$ |
| 1498 | $\nu_{71}$ | 1534 | ($\nu_{72}$) $C_{9a}-N_{10}, C_7-C_6, C_{11,7a,8a}-H_3, C_8-C_9, C_{10a}-N_1, C_4-N_3, N_5-C_{4a}$ | $\nu_{71}$ | 1487 | $C_8-C_7, C_6-C_{5a}, C_9-C_{9a}, C_{6,9}-H, C_{7a}-H_3$ |
| 1570 | $\nu_{73}$ | 1638 | $C_8-C_9, C_6-C_{5a}, C_9-H, C_{9a}-N_{10}, C_7-C_6, C_{10a}-N_1, N_5-C_{4a}$ | $\nu_{73}$ | 1572 | $C_8-C_9, C_6-C_{5a}, C_{6,9}-H, C_2-O_{2'}, C_{9a}-N_{10}, N_3-H, C_{10a}-N_1$ |
| | $\nu_{74}$ | 1726 | $C_2-O_{2'}, C_4-O_{4'}, N_3-H$ (as) | $\nu_{74}$ | 1589 | $C_2-O_{2'}, N_3-H, C_8-C_9, N_{10}-C_{10a}, C_{4a}-C_4, C_{9a}-C_{5a}$ ~(as) |
| 1626 | $\nu_{75}$ | 1753 | $C_2-O_{2'}, C_4-O_{4'}$ (s) | $\nu_{75}$ | 1665 | $C_4-O_{4'}, N_3-H, C_{4a}-C_4$ ~(s) |
| **Exp. RR FMN $S_1$** | **BPBE/cc-pVDZ** | | | **CAM-B3LYP/cc-pVDZ** | | |
| | **$\nu_\#$** | **$S_1$offR** | **Assignment** | **$\nu_\#$** | **$S_1$offR** | **Assignment** |
| 1200 | $\nu_{48}$ | 1126 | $C_6-H, C_8-C_7, C_{7a}-H_3, C_{5a}-N_5, C_{9a}-N_{10}, C_{11}-H_3$ | $\nu_{48}$ | 1181 | $C_6-H, C_{7a}-H_3, C_8-C_7, C_{5a}-N_5, N_{10}-C_{10a}$ |
| | $\nu_{51}$ | 1191 | $C_2-N_3, C_{10a}-C_{4a}, C_{6,9}-H, N_3-H, C_{5a}-N_5$ | $\nu_{49}$ | 1216 | $C_{6,9}-H, N_3-H, C_4-N_3, N_5-C_{4a}, C_8-C_7$ |
| 1250 | $\nu_{53}$ | 1248 | $C_{6,9}-H, N_3-H, C_{11}-H_3, C_{10a}-N_1, C_8-C_7, N_4-C_{4a}$ | $\nu_{50}$ | 1234 | $C_6-H, C_{11}-H_3, C_{4a}-C_4, N_{10}-C_{10a}, C_8-C_7$ |
| 1338 | $\nu_{55}$ | 1303 | $N_{10}-C_{10a}, N_1-C_2, N_3-H, C_{11}-H_3, C_7-C_6, C_{5a}-N_5$ | $\nu_{53}$ | 1319 | $N_{10}-C_{10a}, C_{11}-H_3, C_{9a}-C_{5a}, N_5-C_{4a}, C_9-H, C_7-C_6$ |
| | $\nu_{57}$ | 1333 | $N_3-H, C_{11}-H_3, C_6-H, C_{7a}-H_3, C_{9a}-N_{10}, C_6-C_{5a}$ | $\nu_{54}$ | 1348 | $C_{9a}-C_{5a}, C_{10a}-C_{4a}, C_{5a}-N_5, N_3-H, N_{10}-C_{10a}, C_9-H, N_1-C_2$ |
| 1381 | - | - | - | $\nu_{57}$ | 1402 | $N_3-H, C_{11}-H_3, C_4-N_3, C_{10a}-N_1, C_{5a}-N_5, C_{9a}-C_{5a}$ |
| | $\nu_{58}$ | 1340 | $N_3-H. C_{8a}-H_3, N_1-C_2, N_3-C_4, C_8-C_9, C_6-C_{5a}$ | $\nu_{58}$ | 1407 | $C_{8a,7a}-H_3, C_{9a}-C_{5a}, C_8-C_7, C_6-C_{5a}$ |
| | $\nu_{60}$ | 1353 | $N_3-H, C_{11}-H_3, C_6-H, N_5-C_{4a}, C_{10a}-C_{4a}$ | $\nu_{59}$ | 1409 | $N_3-H, C_{11,7a}-H_3, C_6-H, C_4-N_3, C_{9a}-N_{10}$ |
| | $\nu_{61}$ | 1363 | $C_{6,9}-H, C_7-C_8, C_{9a}-C_{5a}, N_3-H, C_{11}-H_3, C_{8a}-H_3$ | $\nu_{60}$ | 1430 | $C_{11,7a,8a}-H_3, N_{10}-C_{10a}, C_{10a}-N_1, C_{5a}-N_5$ |
| 1416 | $\nu_{64}$ | 1397 | $C_{11}-H_3, C_{8a}-H_3, C_{10a}-C_{4a}, C_8-C_9$ | $\nu_{64}$ | 1459 | $C_{8a}-H_3, C_9-H, C_{10a}-C_{4a}, C_2-N_3, C_{9a}-C_{5a}$ |

| | | | | | | |
|---|---|---|---|---|---|---|
| | - | - | - | $\nu_{68}$ | 1483 | $C_{7a,8a,11}-H_3$, $C_{10a}-C_{4a}$, $C_7-C_6$, $N_3-H$ |
| 1498 | $\nu_{71}$ | 1494 | $C_9-C_{9a}$, $C_6-C_{5a}$, $C_8-C_7$, $C_{6,9}-H$, $C_{7a}-H_3$ | $\nu_{72}$ | 1585 | ($\nu_{71}$) $C_{10a}-N_1$, $N_{5a}-C_4$, $C_{9a}-N_{10}$, $C_4-N_3$, $C_7-C_6$, $C_8-C_9$, $C_{9a}-N_{10}$ |
| 1570 | $\nu_{73}$ | 1579 | $C_2-O_{2'}$, $C_{6,9}-H$, $N_3-H$, $C_8-C_9$, $C_6-C_{5a}$, $N_1-C_{10a}$, $N_5-C_{4a}$ | $\nu_{73}$ | 1661 | $C_8-C_9$, $C_6-C_{5a}$, $C_9-H$, $C_{9a}-N_{10}$, $N_5-C_{4a}$, $C_{10a}-N_1$ |
| | $\nu_{74}$ | 1596 | $C_2-O_{2'}$, $N_3-H$, $C_8-C_9$, $C_6-C_{5a}$, $N_{10}-C_{10a}$, $C_{4a}-C_4$, $C_9-H$ ~(as) | $\nu_{74}$ | 1754 | $C_4-O_{4'}$, $N_3-H$, $C_2-O_{2'}$ (as) |
| 1626 | $\nu_{75}$ | 1671 | $C_4-O_{4'}$, $N_3-H$ ~(s) | $\nu_{75}$ | 1775 | $C_4-O_{4'}$, $C_2-O_{2'}$(s) |
| **Exp. RR FMN $S_1$** | **HCTH/407/cc-pVDZ** | | | **HISSbPBE/cc-pVDZ** | | |
| | **$\nu_\#$** | **$S_1$offR** | **Assignment** | **$\nu_\#$** | **$S_1$offR** | **Assignment** |
| 1200 | $\nu_{48}$ | 1150 | $C_6-H$, $C_{8a}-H_3$, $C_{10a}-N_1$, $N_5-C_{4a}$, $C_6-C_{5a}$, $C_7-C_6$, $C_{5a}-N_5$ | $\nu_{51}$ | 1310 | $C_9-H$, $N_3-H$, $C_4-N_3$, $C_3-C_2$, $C_{9a}-C_{5a}$, $C_8-C_7$ |
| | $\nu_{51}$ | 1216 | $C_2-N_3$, $C_{10a}-C_{4a}$, $C_{9,6}-H$, $C_{9a}-C_{5a}$, $N_{10}-C_{10a}$ | $\nu_{52}$ | 1310 | $C_2-N_3$, $N_3-H$, $C_{6,9}-H$, $C_{10a}-C_{4a}$, $N_{10}-C_{10a}$, $C_8-C_7$ |
| 1250 | $\nu_{53}$ | 1274 | $C_{6,9}-H$, $C_{11}-H_3$, $N_3-H$, $C_{10a}-N_1$ | $\nu_{53}$ | 1362 | $N_{10}-C_{10a}$, $N_5-C_{4a}$, $C_{9a}-C_{5a}$, $C_7-C_6$, $C_{11}-H_3$, $C_8-C_7$, $C_2-N_3$ |
| 1338 | $\nu_{55}$ | 1325 | $N_3-H$, $N_{10a}-C_{10a}$, $C_{5a}-N_5$, $C_{11}-H_3$, $C_7-C_6$, $C_{9a}-C_{5a}$, $C_{10a}-N_1$ | $\nu_{54}$ | 1380 | $C_{5a}-N_5$, $C_9-C_{9a}$, $C_{10a}-C_{4a}$, $C_2-N_3$, $C_{7a}-H_3$, $C_{6,9}-H$, $N_3-H$ |
| | $\nu_{56}$ | 1342 | $C_{7a}-H_3$, $C_9-H$, $C_8-C_7$ | $\nu_{55}$ | 1399 | $C_{7a}-H_3$, $C_8-C_7$, $N_{10}-C_{10a}$, N3-H |
| 1381 | $\nu_{57}$ | 1360 | $C_{8a,11}-H_3$, $N_3-H$, $C_{9a}-N_{10}$ | $\nu_{57}$ | 1427 | $N_3-H$, $C_{5a}-N_5$, $C_{10a}-N_1$, $C_4-N_3$, $C_8-C_7$, $C_{7a,8a,11}-H_3$ |
| | $\nu_{58}$ | 1360 | $C_{8a}-H_3$, $N_3-H$, $C_6-H$, $C_{9a}-C_9$, $C_7-C_6$ | $\nu_{58}$ | 1430 | $C_{11,7a}-H_3$, $N_3-H$, $C_7-C_6$, $C_6-C_{5a}$, $C_{10a}-C_{4a}$, $N_3-C_2$ |
| | $\nu_{59}$ | 1370 | $N_3-H$, $C_{8a}-H_3$, $N_1-C_2$, $C_{9a}-N_{10}$, $C_4-N_3$, $N_5-C_{4a}$ | $\nu_{59}$ | 1440 | $C_{11,8a,7a}-H_3$, $C_8-C_7$, $C_6-C_{5a}$, $C_9-C_{9a}$, $N_3-H$ |
| | $\nu_{61}$ | 1388 | $N_3-H$, $C_6-H$, $C_{9a}-C_{5a}$, $C_8-C_7$, $C_{7a,8a,11}-H_3$, $C_{10a}-C_{4a}$, $C_4-N_3$ | $\nu_{60}$ | 1457 | $C_{7a,11}-H_3$, $N_{10}-C1_{0a}$, $C_8-C_7$, $C_4-N_3$, $N_3-H$ |
| 1416 | $\nu_{65}$ | 1422 | $C_{8a,11}-H_3$, $C_9-H$, $N_{10}-C_{10a}$, $N_1-C_2$, $C_8-C_9$ | $\nu_{63}$ | 1474 | $C_{5a}-C_{9a}$, $C_6-H$, $N_3-H$, $C_4-N_3$, $C_{10a}-N_1$ |
| | - | - | - | $\nu_{65}$ | 1492 | $C_{8a,11}-H_3$, $N_1-C_2$, $C_{4a}-C_4$, $C_{9a}-C_{5a}$ |
| 1498 | $\nu_{71}$ | 1538 | $C_8-C_7$, $C_6-C_{5a}$, $C_9-C_{9a}$, $C_{6,9}-H$ | $\nu_{70}$ | 1580 | $C_8-C_7$, $C_9-C_{9a}$, $C_{9a}-C_{5a}$, $C_6-C_{5a}$, $C_{6,9}-H$, $C_{7a}-H_3$, $N_1-C_{10a}$ |
| 1570 | $\nu_{73}$ | 1613 | $C_8-C_9$, $C_6-C_{5a}$, $C_{6,9}-H$, $C_9-C_{9a}$, $C_7-C_6$, $N_3-H$, $C_2=O_{2'}$, $C_{10a}-N_1$, $C_4-N_3$ | $\nu_{73}$ | 1702 | $C_8-C_9$, $C_6-C_{5a}$, $C_{6,9}-H$, $C_7-C_6$, $N_{10}-C_{9a}$, $N_5-C_{4a}$, $C_{10a}-N_1$ |
| | $\nu_{74}$ | 1640 | $C_2-O_{2'}$, $N_3-H$, $C_{10a}-C_{4a}$, $C_4-N_3$, $C_8-C_9$ | $\nu_{74}$ | 1794 | $C_2-O_{2'}$, $N_3-H$, $C_4-O_{4'}$ (as), $C_{4a}-C_{10a}$, $C_4-N_3$ |
| 1626 | $\nu_{75}$ | 1711 | $C_2-O_{2'}$, $N_3-H$, $C_{4a}-C_4$ ~(s) | $\nu_{75}$ | 1813 | $C_4-O_{4'}$, $C_2-O_{2'}$ (s), $N_3-H$, $C_{4a}-C_{10a}$ |
| **Exp. RR FMN $S_1$** | **HSEH1PBE/cc-pVDZ** | | | **LC-OPBE/cc-pVDZ** | | |
| | **$\nu_\#$** | **$S_1$offR** | **Assignment** | **$\nu_\#$** | **$S_1$offR** | **Assignment** |
| 1200 | $\nu_{49}$ | 1201 | $C_{6,9}-H$, $N_3-H$, $C_{8a}-H_3$, $C_{5a}-N_5$, $C_7-C_6$ | - | - | - |
| | $\nu_{50}$ | 1215 | $C_{6,9}-H$, $N_3-H$, $C_{11}-H_3$, $C_4-N_3$, $C_{10a}-N_1$, $N_{10}-C_{10a}$, $C_{5a}-N_5$ | $\nu_{50}$ | 1266 | $C_{6,9}-H$, $C_{11}-H_3$, $C_{4a}-C_4$, $C_{9a}-C_{5a}$ |
| 1250 | $\nu_{51}$ | 1257 | $C_9-H$, $N_3-H$, $C_{4a}-C_4$, $N_1-C_2$, $N_5-C_{4a}$, $C_{9a}-C_{5a}$ | $\nu_{51}$ | 1308 | $C_9-H$, $N_3-H$, $C_{11}-H_3$, $N_{10}-C_{10a}$, $C_4-C_{4a}$ |
| 1338 | - | - | - | $\nu_{53}$ | 1382 | $C_{7a,8a}-H_3$, $C_{6,9}-H$, $C_{9a}-C_{5a}$, $C_8-C_7$, $N_3-H$ |
| | $\nu_{56}$ | 1369 | $C_{7a}-H_3$, $N_3-H$, $C_6-H$, $C_{5a}-N_5$, $C_{10a}-N_1$ | - | - | - |
| 1381 | $\nu_{58}$ | 1389 | $C_{11}-H_3$, $C_6-H$, $N_3-H$, $C_{7a}-H_3$, $C_7-C_6$, $C_{9a}-N_{10}$, $C_{10a}-C_{4a}$ | $\nu_{59}$ | 1439 | $C_{7a,8a}-H_3$, $N_3-H$, $C_{11}-H_3$, $C_6-H$ |
| | $\nu_{59}$ | 1402 | $C_{11}-H_3$, $C_{8a}-H_3$, $C_9-H$, $N_3-H$, $C_8-C_9$, $C_6-C_{5a}$, $C_4-N_3$ | $\nu_{61}$ | 1459 | $C_{7a,8a}-H_3$, $C_{11}-H_3$, $C_9-H$, $C_{9a}-C_{5a}$, $C_{11}-N_{10}$ |
| | $\nu_{61}$ | 1422 | $C_{11}-H_3$, $C_{8a}-H_3$, $N_3-H$, $C_{9a}-C_{5a}$, $C_{9a}-N_{10}$, $N_1-C_2$, $N_5-C_{4a}$ | $\nu_{64}$ | 1474 | $N_3-H$, $C_{11}-H_3$, $C_4-N_3$, $C_9-C_{9a}$, $C_8-C_7$, $C_{6,9}-H$ |

| | | | | | | |
|---|---|---|---|---|---|---|
| | $\nu_{62}$ | 1423 | $N_3$-H, $C_6$-H, $C_{7a}$-$H_3$, $C_9$-$C_{9a}$, $C_{9a}$-$C_{5a}$, $C_4$-$N_3$ | $\nu_{66}$ | 1538 | $C_{7a}$-$H_3$, $C_7$-$C_6$, $C_8$-$C_9$, $C_{9a}$-$N_{10}$, $C_{5a}$-$N_5$, $C_{4a}$-$C_4$, $C_{10a}$-$N_1$, $C_2$-$N_3$ |
| 1416 | - | - | - | $\nu_{67}$ | 1557 | $C_{4a}$-$C_{10a}$, $C_2$-$N_3$, $C_9$-$C_{9a}$, $C_7$-$C_6$, $N_3$-H, $N_3$-$C_4$ |
| | $\nu_{65}$ | 1449 | $C_{8a}$-$H_3$, $C_{11}$-$H_3$, $N_1$-$C_2$, $C_{10a}$-$C_{4a}$ | $\nu_{68}$ | 1594 | $N_5$-$C_{4a}$, $C_{9a}$-$N_{10}$, $C_6$-$C_{5a}$, $C_{10a}$-$N_1$, $C_8$-$C_9$, $C_{6,9}$-H, $C_{11}$-$H_3$ |
| 1498 | $\nu_{71}$ | 1543 | $C_8$-$C_7$, $C_{9a}$-$C_{5a}$, $C_9$-H, $C_{7a}$-$H_3$, $N_5$-$C_{4a}$, $N_{10a}$-$C_{4a}$ | $\nu_{71}$ | 1701 | $C_7$-$C_6$, $C_{9a}$-$N_{10}$, $C_{9a}$-$C_{5a}$, $C_{6,9}$-H, $C_{10a}$-$C_{4a}$ |
| 1570 | $\nu_{73}$ | 1653 | $C_8$-$C_9$, $C_6$-$C_{5a}$, $C_{9a}$-$N_{10}$, $C_{6,9}$-H, $C_{10a}$-$N_1$, $N_5$-$C_{4a}$, $C_2$-$O_{2'}$ | $\nu_{73}$ | 1788 | $C_{10a}$-$N_1$, $N_5$-$C_{4a}$, $C_4$-$N_3$, $C_{5a}$-$N_5$, $C_{9a}$-$N_{10}$, $C_8$-$C_9$, $N_3$-H |
| | $\nu_{74}$ | 1728 | $C_2$-$O_{2'}$, $N_3$-H, $C_4$-$O_{4'}$ (as) | $\nu_{74}$ | 1865 | $C_2$-$O_{2'}$, $C_4$-$O_{4'}$ (as), $N_3$-H, $C_{4a}$-$C_{10a}$ |
| 1626 | $\nu_{75}$ | 1754 | $C_4$-$O_{4'}$, $N_3$-H, $C_2$-$O_{2'}$ (s) | $\nu_{75}$ | 1901 | $C_4$-$O_{4'}$, $C_2$-$O_{2'}$ (s), $N_3$-H, $C_{4a}$-$N_5$, $C_{10a}$-$N_1$ |
| **Exp. RR FMN $S_1$** | **LC-wHPBE/cc-pVDZ** | | | **LSDA/cc-pVDZ** | | |
| | **$\nu_\#$** | **$S_1$offR** | **Assignment** | **$\nu_\#$** | **$S_1$offR** | **Assignment** |
| 1200 | $\nu_{54}$ | 1369 | $N_3$-H, $C_{9a}$-$C_{5a}$, $C_{5a}$-$N_5$, $C_{10a}$-$C_{4a}$, $C_{7a}$-$H_3$, $C_{6,9}$-H | $\nu_{49}$ | 1160 | $C_{8a,7a}$-$H_3$, $C_8$-$C_7$, $C_7$-$C_6$, $C_{6,9}$-H, $C_{10a}$-$N_1$, $C_{9a}$-$N_{10}$ |
| | $\nu_{55}$ | 1383 | $C_{7a,8a}$-$H_3$, $C_9$-H, $C_{9a}$-$C_{5a}$, | - | - | - |
| 1250 | $\nu_{60}$ | 1437 | $C_{7a,8a}$-$H_3$, $C_{11}$-$H_3$, $C_4$-$N_3$, $C_{5a}$-$N_5$ | $\nu_{53}$ | 1266 | $C_{10a}$-$N_1$, $N_3$-H, $C_6$-H, $C_{11,8a}$-$H_3$, $C_6$-$C_{5a}$ |
| 1338 | - | - | - | $\nu_{57}$ | 1322 | $C_{7a,11}$-$H_3$, $N_3$-H, $N_{10}$-$C_{10a}$, $C_2$-$N_3$, $C_{6a}$-$N_5$ |
| | $\nu_{64}$ | 1469 | $C_{11,8a}$-$H_3$, $C_9$-H, $N_1$-$C_2$, $C_{9a}$-$C_{5a}$, $C_4$-$N_3$, $C_8$-$C_7$ | $\nu_{58}$ | 1324 | $C_{11}$-$H_3$, $N_3$-H, $C_{9a}$-$N_{10}$ |
| 1381 | - | - | - | - | - | - |
| | $\nu_{65}$ | 1469 | $C_{11}$-$H_3$, $C_{8a}$-$H_3$, $C_9$-H, $N_1$-$C_2$ | $\nu_{65}$ | 1390 | $C_{8a,11}$-$H_3$, $N_1$-$C_2$, $C_4$-$N_3$, $C_9$-$C_{9a}$, $C_8$-$C_7$, $N_{10}$-$C_{10a}$ |
| | $\nu_{66}$ | 1477 | $C_{7a}$-$H_3$, $C_{11}$-$H_3$, $C_{5a}$-$N_5$, $N_5$-$C_{4a}$, $C_4$-$N_3$, $C_{10a}$-$N_1$ | $\nu_{66}$ | 1402 | $C_8$-$C_7$, $C_{7a,11}$-$H_3$, $C_{9a}$-$C_{5a}$, $C_{10a}$-$C_{4a}$, $C_6$-$C_{5a}$, $C_9$-$C_{9a}$, $N_1$-$C_2$ |
| | $\nu_{67}$ | 1499 | $C_{10a}$-$C_{4a}$, $C_9$-$C_{9a}$, $C_{4a}$-$C_4$, $N_1$-$C_2$, $C_{11}$-$H_3$, $C_{5a}$-$N_5$ | - | - | - |
| 1416 | - | - | - | $\nu_{67}$ | 1413 | $C_{9a}$-$C_{5a}$, $C_{9a}$-$N_{10}$, $C_{11,7a,8a}$-$H_3$, $C_8$-$C_7$, $C_{4a}$-$C_4$, $N_1$-$C_2$ |
| | $\nu_{70}$ | 1579 | $C_6$-$C_{5a}$, $C_{5a}$-$N_5$, $C_8$-$C_7$, $C_{10a}$-$N_1$, $C_{6,9}$-H | - | - | - |
| 1498 | $\nu_{71}$ | 1629 | $C_7$-$C_6$, $C_{9a}$-$C_{5a}$, $C_{9a}$-$N_{10}$, $C_{6,9}$-H, $N_5$-$C_{4a}$, $C_{10a}$-$C_{4a}$ | $\nu_{71}$ | 1529 | $C_9$-$C_{9a}$, $C_8$-$C_7$, $C_6$-$C_{5a}$, $C_{9,6}$-H |
| 1570 | $\nu_{72}$ | 1675 | $N_5$-$C_{4a}$, $C_{10a}$-$N_1$, $C_6$-$C_{5a}$, $C_8$-$C_9$, $C_4$-$N_3$, $C_{6,9}$-H | $\nu_{73}$ | 1623 | $C_8$-$C_9$, $C_6$-$C_{5a}$, $C_{9,6}$-H, $C_9$-$C_{9a}$, $C_7$-$C_6$, $C_{10}$-$N_1$, $N_4$-$C_{4a}$, $C_2$-$O_{2'}$ |
| | $\nu_{73}$ | 1695 | $C_8$-$C_9$, $C_{10a}$-$N_1$, $N_5$-$C_{4a}$, $C_6$-$C_{5a}$, $C_9$-H | $\nu_{74}$ | 1652 | $C_2$-$O_{2'}$, $N_3$-H, $N_{10}$-$C_{10a}$, $C_{4a}$-$C_4$ |
| | $\nu_{74}$ | 1790 | $C_4$-$O_{4'}$, $C_2$-$O_{2'}$, $N_3$-H (as) | - | - | - |
| 1626 | $\nu_{75}$ | 1821 | $C_2$-$O_{2'}$, $C_4$-$O_{4'}$ (s) | $\nu_{75}$ | 1730 | $C_4$-$O_{4'}$, $N_3$-H, $C_{4a}$-$C_4$ ~(s) |
| **Exp. RR FMN $S_1$** | **LSDA/aug-cc-pVDZ** | | | **M05-2X/aug-cc-pVDZ** | | |
| | **$\nu_\#$** | **$S_1$offR** | **Assignment** | **$\nu_\#$** | **$S_1$offR** | **Assignment** |
| 1200 | - | - | - | $\nu_{50}$ | 1239 | $C_6$-H, $C_{11}$-$H_3$, $C_9$-$C_{9a}$, $C_7$-$C_6$, $C_2$-$N_3$, $C_{10a}$-$C_{4a}$ |
| | $\nu_{52}$ | 1259 | $N_3$-H, $C_2$-$O_{2'}$, $C_4$-$N_3$, $C_{10a}$-$N_1$, $C_6$-$C_{5a}$ | - | - | - |
| 1250 | $\nu_{53}$ | 1282 | $N_3$-H, $C_2$-$O_{2'}$, $C_{4a}$-$C_4$, $C_4$-$N_3$, $C_{11}$-$H_3$, $N_{10}$-$C_{10a}$, $C_6$-$C_{5a}$, $C_8$-$C_7$ | $\nu_{51}$ | 1266 | $N_3$-H, $C_4$-$N_3$, $N_3$-H, $N_1$-$C_2$, $C_{6,9}$-H |
| 1338 | $\nu_{54}$ | 1305 | $N_5$-$C_{4a}$, $N_{10}$-$C_{11}$, $C_{9a}$-$C_{5a}$, $C_{7a,8a}$-$H_3$, $C_6$-H, $C_{4a}$-$C_4$, $C_8$-$C_9$ | $\nu_{53}$ | 1327 | $C_{9a}$-$C_{5a}$, $N_{10}$-$C_{10a}$, $N_4$-$C_{4a}$, $C_{11}$-$H_3$, $C_8$-$C_7$ |
| | - | - | - | $\nu_{54}$ | 1346 | $N_3$-H, $C_{5a}$-$N_5$, $C_9$-$C_{9a}$, $N_{10}$-$C_{10a}$, $C_9$-H, $C_{4a}$-$C_{10a}$ |
| 1381 | - | - | - | $\nu_{57}$ | 1410 | $N_3$-H, $C_{11}$-$H_3$, $C_4$-$N_3$, $C_{10a}$-$N_1$ |

| | | | | | | |
|---|---|---|---|---|---|---|
| | $\nu_{58}$ | 1355 | $C_{11,7a}$-$H_3$, $N_3$-H, $C_{4a}$-$C_4$, $C_{10a}$-$C_{4a}$, $C_{9a}$-$C_{5a}$, $C_2$-$N_3$ | $\nu_{58}$ | 1412 | $N_3$-H, $C_6$-H, $C_{11}$-$H_3$, $C_7$-$C_6$, $C_{9a}$-$N_{10}$ |
| | $\nu_{62}$ | 1377 | $C_{7a,8a,11}$-$H_3$, $N_3$-H, $C_{11}$-$N_{10}$, $C_8$-$C_7$, $N_{10}$-$C_{10a}$ | - | - | - |
| | $\nu_{63}$ | 1383 | $C_8$-$C_7$, $C_9$-$C_{9a}$, $C_{5a}$-$N_5$, $C_{7a,8a}$-$H_3$, $N_3$-H, $C_2$-$N_3$ | - | - | - |
| 1416 | $\nu_{67}$ | 1426 | $N_3$-H, $C_2$-$N_3$, $C_{11,8a}$-$H_3$, $C_8$-$C_9$, $C_6$-$C_{5a}$, $C_{9a}$-$C_{5a}$, $N_{10}$-$C_{10a}$ | $\nu_{60}$ | 1451 | $C_{11,7a,8a}$-$H_3$, $C_{9a}$-$N_{10}$, $C_{10a}$-$C_{4a}$ |
| | $\nu_{68}$ | 1467 | $N_3$-H, $C_2$-$N_3$, $C_{4a}$-$C_{10a}$, $C_8$-$C_9$, $C_7$-$C_6$, $C_{6,9}$-H, $C_{8a}$-$H_3$ | $\nu_{63}$ | 1476 | $C_{7a,8a,11}$-$H_3$, $C_9$-H, $C_9$-$C_{9a}$, $C_{10a}$-$C_{4a}$, $N_1$-$C_2$ |
| 1498 | $\nu_{71}$ | 1547 | $C_9$-$C_{9a}$, $C_{10a}$-$N_1$, $N_{10a}$-$C_{10a}$, $C_8$-$C_7$, $C_{11}$-$H_3$, $C_9$-H, $C_6$-$C_{5a}$ | $\nu_{72}$ | 1594 | $C_{10a}$-$N_1$, $N_5$-$C_{4a}$, $C_4$-$N_3$, $C_8$-$C_9$, $C_{9a}$-$N_{10}$, $C_9$-H |
| 1570 | $\nu_{72}$ | 1577 | $C_8$-$C_7$, $C_9$-$C_{9a}$, $C_{9a}$-$C_{5a}$, $C_6$-$C_{5a}$, $C_{10a}$-$C_{4a}$, $C_{6,9}$-H, $C_{7a}$-$H_3$, $N_1$-$C_{10a}$ | $\nu_{73}$ | 1662 | $C_8$-$C_9$, $C_6$-$C_{5a}$, $C_9$-H, $C_{9a}$-$N_{10}$, $C_{10a}$-$N_1$, $N_5$-$C_{4a}$ |
| | $\nu_{73}$ | 1611 | $N_1$-$C_2$, $C_4$-$O_{4'}$, $N_3$-H, $C_{10a}$-$C_{4a}$ | $\nu_{74}$ | 1706 | $C_4$-$O_{4'}$, $C_2$-$O_{2'}$, $N_3$-H (as) |
| 1626 | $\nu_{75}$ | 1702 | $C_4$-$O_{4'}$, $N_1$-$C_2$, $C_2$-$O_{2'}$ (s), $N_3$-H, $C_{4a}$-$C_4$ | $\nu_{75}$ | 1731 | $C_2$-$O_{2'}$, $C_4$-$O_{4'}$ (s) |
| **Exp. RR FMN $S_1$** | **M06/cc-pVDZ** | | | **M06/aug-cc-pVDZ** | | |
| | **$\nu_\#$** | **$S_1$offR** | **Assignment** | **$\nu_\#$** | **$S_1$offR** | **Assignment** |
| 1200 | $\nu_{49}$ | 1183 | $C_{6,9}$-H, $N_3$-H, $C_4$-$N_3$, $C_{5a}$-$N_5$, $C_8$-$C_7$, $C_2$-$N_3$ | $\nu_{49}$ | 1188 | $C_{6,9}$-H, $N_3$-H, $C_4$-$N_3$, $C_{11}$-$H_3$, $C_{5a}$-$N_5$ |
| | $\nu_{50}$ | 1198 | $C_{6,9}$-H, $C_9$-$C_{9a}$, $C_{4a}$-$C_4$, $N_{10}$-$C_{10a}$, $C_{10a}$-$N_1$, $C_2$-$N_3$, $C_7$-$C_6$ | $\nu_{50}$ | 1203 | $C_{6,9}$H, $C_2$-$C_3$, $C_{10a}$-$C_{4a}$, $C_9$-$C_{9a}$, $C_{11}$-$H_3$ |
| 1250 | $\nu_{51}$ | 1234 | $N_3$-H, $C_4$-$N_3$, $C_{4a}$-$C_4$, $N_1$-$C_2$, $C_9$-H, $N_5$-$C_{4a}$, $C_{9a}$-$C_{5a}$ | $\nu_{51}$ | 1235 | $N_3$-H, $C_2$-$N_3$, $N_3$-$C_4$, $C_{4a}$-$C_4$, $C_{6,9}$-H, $C_{5a}$-$N_5$, $C_7$-$C_6$ |
| 1338 | $\nu_{52}$ | 1250 | $C_9$-H, $C_8$-$C_7$, $N_{10}$-$C_{10a}$, $C_{10a}$-$C_{4a}$, $N_1$-$C_2$, $C_9$-$C_{9a}$ | - | - | - |
| | $\nu_{56}$ | 1357 | $C_{8a,11}$-$H_3$, $N_3$-H, $C_{5a}$-$N_5$, $C_{10a}$-$N_1$, $C_4$-$N_3$, $C_{9a}$-$N_{10}$, $C_8$-$C_7$ | $\nu_{56}$ | 1360 | $N_3$-H, $C_{11}$-$H_3$, $C_{10a}$-$N_1$, $C_{5a}$-$N_5$, $N_3$-$C_4$, $C_{9a}$-$N_{10}$ |
| 1381 | $\nu_{58}$ | 1367 | $N_3$-H, $C_6$-H, $C_{10a}$-$C_{4a}$, $C_{11}$-$H_3$, $C_{9a}$-$C_{5a}$, $C_7$-$C_6$, $N_3$-$C_2$ | $\nu_{57}$ | 1367 | $N_3$-H, $C_6$-H, $C_{7a,11}$-$H_3$, $C_7$-$C_6$, $C_{10a}$-$C_{4a}$, $C_{9a}$-$C_{5a}$ |
| | $\nu_{59}$ | 1380 | $C_{8a,7a,11}$-$H_3$, $C_6$-$C_{5a}$, $C_8$-$C_7$, $C_{6,9}$-H | $\nu_{58}$ | 1375 | $C_{8a}$-$H_3$, $N_3$-H, $C_4$-$N_3$, $C_{9a}$-$C_{5a}$, $C_7$-$C_8$ |
| | $\nu_{63}$ | 1410 | $N_3$-H, $C_{9a}$-$C_{5a}$, $C_6$-H, $C_{10a}$-$C_{4a}$, $C_4$-$N_3$ | $\nu_{59}$ | 1385 | $C_{7a,8a,11}$-$H_3$, $N_3$-H, $C_6$-$C_{5a}$, $C_8$-$C_9$, $N_{10}$-$C_{10a}$ |
| | $\nu_{66}$ | 1425 | $C_{8a,11}$-$H_3$, $C_{10a}$-$C_{4a}$, $C_4$-$N_3$, $C_9$-H, $N_1$-$C_2$, $C_8$-$C_7$ | - | - | - |
| 1416 | $\nu_{68}$ | 1450 | $N_{10}$-$C_{10a}$, $N_5$-$C_4$, $C_{11}$-$H_3$, $C_{10a}$-$N_1$, $C_9$-$C_{9a}$, $C_8$-$C_7$ | $\nu_{66}$ | 1433 | $C_{8a}$-$H_3$, $C_9$-H, $C_{11}$-$H_3$, $N_1$-$C_2$, $C_4$-$N_3$, $C_6$-$C_{5a}$ |
| | $\nu_{69}$ | 1460 | $N_5$-$C_{4a}$, $C_8$-$C_7$, $C_{10a}$-$N_1$, $C_{7a,8a}$-$H_3$, $C_{9a}$-$C_{5a}$, $C_2$-$N_3$, $N_3$-H | $\nu_{69}$ | 1457 | $C_{8a,11}$-$H_3$, $N_5$-$C_{4a}$, $N_{10}$-$C_{10a}$, $N_1$-$C_2$, $C_4$-$N_3$ |
| 1498 | $\nu_{71}$ | 1526 | $C_8$-$C_7$, $C_{9a}$-$C_{5a}$, $C_{10a}$-$C_{4a}$, $C_{6,9}$-H, $C_{7a}$-$H_3$ | $\nu_{71}$ | 1518 | $C_8$-$C_7$, $C_{9a}$-$C_{5a}$, $C_{10a}$-$C_{4a}$, $C_{6,9}$-H, $C_{7a}$-$H_3$ |
| 1570 | $\nu_{73}$ | 1646 | $C_8$-$C_9$, $C_6$-$C_{5a}$, $C_{6,9}$-H, $C_{10a}$-$N_1$, $C_{4a}$-$C_4$ | $\nu_{73}$ | 1644 | $C_8$-$C_9$, $C_6$-$C_{5a}$, $C_{6,9}$-H, $C_{9a}$-$N_{10}$, $N_4$-$C_{4a}$, $C_2$-$O_{2'}$, $C_{10a}$-$N_1$ |
| | $\nu_{74}$ | 1747 | $C_2$-$O_{2'}$, $C_4$-$O_{4'}$ (as), $N_3$-H, $C_{4a}$-$C_4$, $C_4$-$N_3$ | $\nu_{74}$ | 1696 | $C_2$-$O_{2'}$, $N_3$-H, $C_4$-$O_{4'}$ (as) |
| 1626 | $\nu_{75}$ | 1770 | $C_2$-$O_{2'}$, $C_4$-$O_{4'}$ (s), $N_3$-H, $C_{4a}$-$C_4$ | $\nu_{75}$ | 1717 | $C_4$-$O_{4'}$, $C_2$-$O_{2'}$ (s) |
| **Exp. RR FMN $S_1$** | **M06-HF/cc-pVDZ** | | | **M06L/cc-pVDZ** | | |
| | **$\nu_\#$** | **$S_1$offR** | **Assignment** | **$\nu_\#$** | **$S_1$offR** | **Assignment** |
| 1200 | $\nu_{49}$ | 1183 | $C_{6,9}$-H, $N_3$-H, $C_{11,8a}$-$H_3$, $C_4$-$N_3$, $C_8$-$C_7$, $N_{10}$-$C_{10a}$ | $\nu_{51}$ | 1255 | $C_9$-H, $C_2$-$N_3$, $C_{10a}$-$C_{4a}$, $C_4$-$N_3$, $N_3$-H, $C_6$-$C_{5a}$, $C_8$-$C_9$ |
| | $\nu_{50}$ | 1198 | $C_{6,9}$-H, $C_4$-$N_3$, $C_{4a}$-$C_4$, $C_{10a}$-$N_1$, $C_{9a}$-$N_{10}$ | $\nu_{52}$ | 1266 | $C_{6,9}$-H, $C_2$-$O_{2'}$, $N_3$-H, $C_8$-$C_7$, $C_{9a}$-$N_{10}$ |
| 1250 | $\nu_{51}$ | 1234 | $N_3$-H, $C_9$-H, $C_2$-$N_3$, $C_{4a}$-$C_4$, $N_5$-$C_{4a}$, $C_7$-$C_6$ | $\nu_{53}$ | 1301 | $C_{6,9}$-H, $C_2$-$O_{2'}$, $C_{4a}$-$C_4$, $N_3$-H, $N_{10}$-$C_{10a}$, $C_9$-$C_{9a}$, $C_8$-$C_7$ |
| 1338 | $\nu_{54}$ | 1317 | $C_{9a}$-$C_{5a}$, $C_{5a}$-$N_5$, $C_{10a}$-$C_{4a}$, $C_6$-H, $C_{7a}$-$H_3$, $N_{10}$-$C_{10a}$ | $\nu_{55}$ | 1347 | $C_4$-$C_{4a}$, $C_{10a}$-$N_1$, $C_{9a}$-$N_{10}$, $C_{5a}$-$N_5$, $N_3$-H, $C_8$-$C_7$, $C_{11,8a,7a}$-$H_3$ |

| | | | | | | |
|---|---|---|---|---|---|---|
| | $v_{56}$ | 1357 | $C_{11,8a}$-$H_3$, $N_3$-H, $C_{5a}$-$N_5$, $C_{10a}$-$N_1$ | $v_{56}$ | 1365 | $C_{5a}$-$N_5$, $C_8$-$C_7$, $C_{9a}$-$C_{5a}$, $C_{4a}$-$C_4$, $N_3$-H, $C_9$-H, $C_{7a,8a}$-$H_3$ |
| 1381 | $v_{57}$ | 1363 | $C_{8a,11}$-$H_3$, $C_{10a}$-$N_1$, $C_{5a}$-$N_5$ | $v_{61}$ | 1424 | $C_{11}$-$H_3$, $N_3$-H, $N_{10}$-$C_{10a}$, $C_2$-$N_3$, $C_7$-$C_6$, $C_8$-$C_9$ |
| | $v_{58}$ | 1367 | $N_3$-H, $C_6$-H, $C_{11}$-$H_3$, $C_{10a}$-$C_{4a}$, $C_4$-$N_3$, $C_7$-$C_6$ | $v_{63}$ | 1432 | $C_{7a,8a}$-$H_3$, $C_{5a}$-$N_5$, $C_7$-$C_6$ |
| | $v_{59}$ | 1380 | $C_{7a,8a,11}$-$H_3$, $C_6$-$C_{5a}$, $C_{10a}$-$N_1$ | - | - | - |
| | - | - | - | $v_{65}$ | 1446 | $C_{11,7a,8a}$-$H_3$, $C_{9a}$-$C_{5a}$, $C_{6,9}$-H |
| 1416 | $v_{63}$ | 1410 | $N_3$-H, $C_{5a}$-$C_{9a}$, $C_6$-H, $C_{10a}$-$C_{4a}$, $C_4$-$N_3$ | $v_{67}$ | 1458 | $C_{8a,11}$-$H_3$, $N_3$-H, $C_{9a}$-$N_{10}$, $C_6$-$C_{5a}$, $C_8$-$C_9$, $C_{10a}$-$N_1$ |
| | $v_{66}$ | 1425 | $C_{8a,11}$-$H_3$, $C_9$-H, $C_{10a}$-$C_{4a}$, $N_1$-$C_2$, $C_{4a}$-$C_4$ | $v_{68}$ | 1490 | $N_3$-H, $C_{10a}$-$C_{4a}$, $C_2$-$C_3$, $C_8$-$C_9$, $C_7$-$C_6$, $C_{10a}$-$C_{4a}$, $C_{6,9}$-H, $C_{11,7a,8a}$-$H_3$ |
| 1498 | $v_{71}$ | 1526 | $C_8$-$C_7$, $C_{9a}$-$C_{5a}$, $C_{10a}$-$C_{4a}$, $C_{9,6}$-H, $C_{7a}$-$H_3$ | $v_{71}$ | 1531 | $C_8$-$C_7$, $C_9$-$C_{9a}$, $C_6$-$C_{5a}$, $C_{6,9}$-H, $C_{10a}$-$N_1$, $N_5$-$C_{4a}$, $C_{8a}$-$H_3$ |
| 1570 | $v_{73}$ | 1646 | $C_8$-$C_9$, $C_6$-$C_{5a}$, $C_{9,6}$-H, $C_{10a}$-$N_1$, $N_5$-$C_{4a}$ | $v_{73}$ | 1626 | $N_1$-$C_2$, $N_3$-H, $C_4$-$O_{4'}$, $C_4$-$N_3$, $C_{9a}$-$C_{5a}$, $C_8$-$C_7$ |
| | $v_{74}$ | 1747 | $C_2$-$O_{2'}$, $N_3$-H, $C_4$-$O_{4'}$ (as) | $v_{74}$ | 1661 | $C_8$-$C_9$, $C_6$-$C_{5a}$, $C_7$-$C_6$, $C_{6,9}$-H, $C_9$-$C_{9a}$, $C_{10a}$-$C_{4a}$ |
| 1626 | $v_{75}$ | 1770 | $C_4$-$O_{4'}$, $C_2$-$O_{2'}$, $N_3$-H (**s**) | $v_{75}$ | 1761 | $C_4$-$O_{4'}$, $C_2$-$O_{2'}$ (**s**), $N_3$-H, $C_{4a}$-$C_4$, $C_{10a}$-$N_1$ |
| **Exp. RR FMN $S_1$** | **M06L/aug-cc-pVDZ** | | | **M11L/cc-pVDZ** | | |
| | **v#** | **$S_1$offR** | **Assignment** | **v#** | **$S_1$offR** | **Assignment** |
| 1200 | $v_{48}$ | 1174 | $C_6$-H, $C_{7a,8a}$-$H_3$, $C_8$-$C_7$, $C_8$-$C_9$, $N_{10}$-$C_{10a}$, $N_5$-$C_{4a}$ | - | - | - |
| | $v_{49}$ | 1180 | $C_6$-H, $C_{5a}$-$N_5$, $C_7$-$C_6$, $C_{7a,11}$-$H_3$, $N_3$-H | $v_{49}$ | 1178 | $C_{8a,7a}$-$H_3$, $C_6$-H, $C_8$-$C_7$, $C_7$-$C_6$, $N_3$-H |
| 1250 | $v_{51}$ | 1226 | $N_3$-H, $C_2$-$N_3$, $C_{9,6}$-H, $C_{4a}$-$C_4$, $N_5$-$C_{4a}$, $C_{5a}$-$C_{9a}$, $N_{10}$-$C_{10a}$ | $v_{52}$ | 1269 | $N_3$-H, $C_2$-$N_3$, $C_{4a}$-$C_4$, $N_5$-$C_{4a}$, $C_9$-H, $C_{5a}$-$N_5$ |
| 1338 | $v_{54}$ | 1308 | $C_{6,9}$-H, $C_{9a}$-$C_{5a}$, $C_{10a}$-$C_{4a}$, $C_{5a}$-$N_5$, $C_{10a}$-$N_1$, $C_{7a}$-$H_3$ | $v_{53}$ | 1306 | $C_6$-H, $C_{11,8a}$-$H_3$, $N_3$-H, $C_{10a}$-$N_1$, $N_5$-$C_{4a}$, $C_8$-$C_7$, $C_2$-$N_3$, $C_6$-$C_{5a}$ |
| | $v_{56}$ | 1363 | $C_{7a}$-$H_3$, $C_9$-H, $C_{11}$-$H_3$, $C_{9a}$-$C_{5a}$ | $v_{55}$ | 1339 | $C_{7a}$-$H_3$, $C_6$-H, $C_{9a}$-$C_{5a}$, $C_8$-$C_7$, $C_{10a}$-$N_1$ |
| 1381 | $v_{57}$ | 1370 | $N_3$-H, $C_6$-H, $C_{11,7a}$-$H_3$, $C_{9a}$-$N_{10}$, $C_6$-$C_{5a}$, $C_7$-$C_6$, $N_5$-$C_{4a}$ | $v_{56}$ | 1357 | $C_{7a,11}$-$H_3$, $N_3$-H, $C_6$-$C_{5a}$, $C_{10a}$-$C_{4a}$, $C_{10a}$-$N_1$, $C_{9a}$-$N_{10}$ |
| | $v_{58}$ | 1382 | $C_{8a}$-$H_3$, $C_{11}$-$H_3$, $N_1$-$C_2$, $C_{9a}$-$N_{10}$ | $v_{58}$ | 1375 | $C_{11}$-$H_3$, $C_6$-H, $N_3$-H, $C_7$-$C_6$, $N_{10}$-$C_{10a}$, $C_{9a}$-$C_{5a}$ |
| | $v_{59}$ | 1389 | $C_8a$-$H_3$, $C_8$-$C_9$, $C_6$-$C_{5a}$, $C_{9a}$-$N_{10}$, $N_5$-$C_{4a}$, $C_{10a}$-$N_1$, $C_4$-$N_3$ | $v_{60}$ | 1387 | $C_{11,8a,7a}$-$H_3$, $N_{10}$-$C_{10}a$, $C_6$-$C_{5a}$, $C_{9a}$-$C_{5a}$ |
| | $v_{61}$ | 1403 | $N_3$-H, $C_6$-H, $C_4$-$N_3$, $C_{9a}$-$C_{5a}$, $C_{11}$-$H_3$ | $v_{63}$ | 1414 | $C_{11,8a,7a}$-$H_3$, $C_8$-$C_7$, $C_{9a}$-$C_{5a}$, $C_4$-$N_3$ |
| 1416 | $v_{65}$ | 1438 | $C_{7a,8a,11}$-$H_3$, $C_{10a}$-$N_1$, $N_5$-$C_{4a}$ | $v_{66}$ | 1439 | $C_{11,8a,7a}$-$H_3$, $C_9$-H, $N_1$-$C_2$, $N_5$-$C_{4a}$, $C_8$-$C_7$, $C_{9a}$-$C_{5a}$ |
| | $v_{66}$ | 1440 | $C_{7a,8a}$-$H_3$, $N_5$-$C_{4a}$, $C_4$-$N_3$, $C_{10a}$-$C_4$ | $v_{69}$ | 1486 | $C_{11}$-$H_3$, $C_{4a}$-$C_4$, $N_5$-$C_{4a}$, $N_1$-$C_2$, $N_{10}$-$C_{4a}$, $C_8$-$C_7$ |
| 1498 | $v_{71}$ | 1535 | $C_8$-$C_7$, $C_{9a}$-$C_{5a}$, $C_{6,9}$-H, $C_{7a}$-$H_3$, $C_{10a}$-$C_{4a}$ | $v_{71}$ | 1543 | $C_8$-$C_7$, $C_{9a}$-$C_{5a}$, $C_{6,9}$-H, $C_9$-$C_{9a}$, $C_6$-$C_{5a}$, $C_{10a}$-$C_{4a}$, $C_{7a}$-$H_3$ |
| 1570 | $v_{73}$ | 1635 | $C_2$-$O_{2'}$, $N_3$-H, $C_{6,9}$-H, $C_8$-$C_9$, $C_6$-$C_{5a}$, $C_4$-$O_{4'}$, $N_5$-$C_{4a}$ (as) | $v_{73}$ | 1664 | $C_8$-$C_9$, $C_6$-$C_{5a}$, $C_{6,9}$-H, $C_{9a}$-$N_{10}$, $C_7$-$C_6$, $N_5$-$C_{4a}$, $C_{10a}$-$N_1$ |
| | $v_{74}$ | 1652 | $C_2$-$O_{2'}$, $C_8$-$C_9$, $C_6$-$C_{5a}$, $N_3$-H, $C_{10a}$-$N_1$, $N_5$-$C_{4a}$, $C_{9a}$-$N_{10}$ | $v_{74}$ | 1750 | $C_2$-$O_{2'}$, $N_3$-H, $C_{10a}$-$N_1$, $C_4$-$O_{4'}$ (as) |
| 1626 | $v_{75}$ | 1695 | $C_4$-$O_{4'}$, $N_3$-H, $C_{4a}$-$C_4$, $C_6$-$C_{5a}$, $C_2$-$O_{2'}$ (**s**) | $v_{75}$ | 1789 | $C_4$-$O_{4'}$, $N_3$-H, $C_{4a}$-$C_4$, $C_2$-$O_{2'}$ (**s**) |
| **Exp. RR FMN $S_1$** | **MN15/cc-pVDZ** | | | **MN15/aug-cc-pVDZ** | | |
| | **v#** | **$S_1$offR** | **Assignment** | **v#** | **$S_1$offR** | **Assignment** |
| 1200 | $v_{49}$ | 1196 | $C_{6,9}$-H, $N_3$-H, $C_{8a}$-$H_3$, $C_4$-$N_3$, $C_7$-$C_6$, $C_{5a}$-$N_5$ | - | - | - |
| | $v_{50}$ | 1211 | $C_{6,9}$-H, $C_{11}$-$H_3$, $C_9$-$C_{9a}$, $C_{4a}$-$C_4$, $C_7$-$C_6$ | $v_{50}$ | 1214 | $C_6$-H, $C_{11}$-$H_3$, $C_9$-H, $C_9$-$C_{9a}$, $C_{4a}$-$C_4$, $C_7$-$C_6$ |

| | | | | | | |
|---|---|---|---|---|---|---|
| 1250 | $\nu_{52}$ | 1264 | $N_3$-H, $C_{6,9}$-H, $C_2$-$N_3$, $C_{4a}$-$C_4$, $N_{10}$-$C_{10a}$ | $\nu_{51}$ | 1258 | $C_9$-H, $N_3$-H, $C_4$-$N_3$, $N_1$-$C_2$ |
| 1338 | $\nu_{53}$ | 1313 | $N_{10}$-$C_{10a}$, $C_{11}$-$H_3$, $N_5$-$C_{4a}$, $C_6$-$C_{5a}$, $C_2$-$N_3$, $C_{8a}$-$H_3$, $C_8$-$C_7$ | $\nu_{53}$ | 1308 | $N_{10}$-$C_{10a}$, $C_{5a}$-$N_5$, $N_5$-$C_{4a}$, $C_{9a}$-$C_{5a}$, $C_7$-$C_6$, $C_{11}$-$H_3$ |
| | $\nu_{55}$ | 1351 | $C_{7a}$-$H_3$, $C_9$-H, $N_3$-H, $C_8$-$C_7$, $C_{10a}$-$C_{4a}$, $C_2$-$N_3$ | - | - | - |
| 1381 | $\nu_{58}$ | 1384 | $C_{11,7a}$-$H_3$, $C_6$-H, $C_6$-$C_{5a}$, $C_7$-$C_6$, $C_{10a}$-$N_1$ | $\nu_{58}$ | 1386 | $C_{11,7a}$-$H_3$, $C_6$-H, $C_7$-$C_6$, $C_6$-$C_{5a}$, $C_{5a}$-$N_5$, $C_{10a}$-$N_1$ |
| | $\nu_{59}$ | 1391 | $C_{11,8a,7a}$-$H_3$, $C_8$-$C_7$, $C_{9a}$-$C_{5a}$, $C_6$-$C_{5a}$ | $\nu_{59}$ | 1393 | $C_{7a,8a,11}$-$H_3$, $C_8$-$C_7$, $C_6$-$C_{5a}$, $C_{9a}$-$C_{5a}$ |
| | $\nu_{61}$ | 1405 | $C_{7a,11,8a}$-$H_3$, $N_{10}$-$C_{10a}$, $C_{10a}$-$N_1$, $N_3$-H, $C_4$-$N_3$ | $\nu_{64}$ | 1438 | $C_{7a,8a}$-$H_3$, $C_6$-H, $C_{9a}$-$C_{5a}$, $C_{5a}$-$N_5$, $C_{10a}$-$C_{4a}$ |
| | - | - | - | $\nu_{65}$ | 1446 | $C_{8a,11}$-$H_3$, $C_9$-H, $N_1$-$C_2$, $C_{10a}$-$C_{4a}$, $C_6$-$C_{5a}$, $C_9$-$C_{9a}$ |
| 1416 | $\nu_{67}$ | 1457 | $C_6$-H, $C_7$-$C_6$, $C_{7a,11}$-$H_3$, $C_6$-$C_{5a}$, $C_{9a}$-$C_{5a}$, $C_{10a}$-$C_{4a}$, $C_7$-$C_6$, $N_5$-$C_{4a}$ | $\nu_{67}$ | 1456 | $C_{7a}$-$H_3$, $C_6$-H, $C_7$-$C_6$, $C_6$-$C_{5a}$, $N_5$-$C_{4a}$, $C_{10a}$-$C_{4a}$ |
| | $\nu_{68}$ | 1470 | $C_{11}$-$H_3$, $C_{9a}$-$N_{10}$, $C_{10a}$-$N_1$, $N_{10}$-$C_{10a}$, $N_5$-$C_{4a}$ | $\nu_{68}$ | 1470 | $N_5$-$C_{4a}$, $N_{10a}$-$N_1$, $C_{5a}$-$N_5$, $C_{11}$-$H_3$, $N_3$-H, $C_4$-$N_3$, $C_{9a}$-$C_{5a}$ |
| 1498 | $\nu_{71}$ | 1547 | ($\nu_{70}$, $\nu_{72}$) $C_7$-$C_6$, $C_{9a}$-$N_{10}$, $C_8$-$C_9$, $C_{10a}$-$N_1$, $C_{7a,11}$-$H_3$, $N_5$-$C_{4a}$, $C_4$-$N_3$ | $\nu_{70}$ | 1530 | ($\nu_{72}$) $C_8$-$C_7$, $C_{9a}$-$C_{5a}$, $C_{6,9}$-H, $C_6$-$C_{5a}$, $C_9$-$C_{9a}$, $C_{7a}$-$H_3$, $N_5$-$C_{4a}$ |
| 1570 | $\nu_{73}$ | 1650 | $C_8$-$C_9$, $C_6$-$C_{5a}$, $C_{6,9}$-H, $C_7$-$C_6$, $C_{9a}$-$N_{10}$, $C_{10a}$-$N_1$, $N_5$-$C_{4a}$ | $\nu_{73}$ | 1641 | $C_8$-$C_9$, $C_6$-$C_{5a}$, $C_{6,9}$-H, $C_7$-$C_6$, $C_{9a}$-$N_{10}$, $C_{10a}$-$N_1$, $N_5$-$C_{4a}$ |
| | $\nu_{74}$ | 1757 | $C_2$-$O_{2'}$, $C_4$-$O_{4'}$, $N_3$-H (as) | $\nu_{74}$ | 1698 | $C_2$-$O_{2'}$, $C_4$-$O_{4'}$, $N_3$-H (as) |
| 1626 | $\nu_{75}$ | 1777 | $C_4$-$O_{4'}$, $C_2$-$O_{2'}$, $N_{10}$-$C_{10a}$ (s) | $\nu_{75}$ | 1722 | $C_4$-$O_{4'}$, $C_2$-$O_{2'}$, $N_{10}$-$C_{10a}$ (s) |
| **Exp. RR FMN $S_1$** | **mPW1PW91/cc-pVDZ** | | | **mPWLYP/cc-pVDZ** | | |
| | **$\nu_{\#}$** | **$S_1$offR** | **Assignment** | **$\nu_{\#}$** | **$S_1$offR** | **Assignment** |
| 1200 | $\nu_{49}$ | 1209 | $C_{6,9}$-H, $N_3$-H, $C_{5a}$-$N_5$, $C_7$-$C_6$, $C_4$-$N_3$ | $\nu_{49}$ | 1125 | $C_6$-H, $C_{7a,8a}$-$H_3$, $C_8$-$C_7$, $C_{5a}$-$N_5$, $N_{10}$-$C_{10a}$ |
| | $\nu_{50}$ | 1222 | $C_{6,9}$-H, $C_{11}$-$H_3$, $C_4$-$N_3$, $N_{10}$-$C_{10a}$, $C_{10a}$-$N_1$, $N_5$-$C_{4a}$, $C_9$-$C_{9a}$ | $\nu_{50}$ | 1137 | $C_{6,9}$-H, $C_{11}$-$H_3$, $C_4$-$N_3$, $N_3$-H, $C_{5a}$-$N_5$ |
| 1250 | $\nu_{51}$ | 1262 | $C_2$-$N_3$, $N_3$-H, $C_{4a}$-$C_4$, $C_9$-H, $N_1$-$C_2$, $C_7$-$C_6$, $C_{5a}$-$N_5$ | $\nu_{55}$ | 1282 | $N_{10}$-$C_{10a}$, $C_{11}$-$H_3$, $N_3$-H, $C_{9a}$-$C_{5a}$, $C_7$-$C_6$, $C_{10a}$-$N_1$, $C_{4a}$-$C_4$, $C_{10a}$-$C_{4a}$ |
| 1338 | - | - | - | - | - | - |
| | $\nu_{56}$ | 1377 | $C_{7a}$-$H_3$, $N_3$-H, $C_{5a}$-$N_5$, $C_{10a}$-$N_1$ | $\nu_{60}$ | 1341 | $N_3$-H, $C_{8a,7a}$-$H_3$, $C_6$-H, $C_{9a}$-$C_{5a}$, $C_8$-$C_7$, $C_2$-$N_3$ |
| 1381 | $\nu_{58}$ | 1396 | $N_3$-H, $C_{11,7a}$-$H_3$, $C_6$-H, $C_7$-$C_6$, $C_{9a}$-$N_{10}$, $C_{10a}$-$C_{4a}$ | - | - | - |
| | $\nu_{59}$ | 1408 | $C_{8a,11}$-$H_3$, $C_8$-$C_7$, $N_3$-H, $C_{9a}$-$C_{5a}$, $C_4$-$N_3$, $C_6$-$C_{5a}$ | $\nu_{62}$ | 1365 | $C_{11}$-$H_3$, $C_{4a}$-$C_4$, $N_1$-$C_2$, $C_4$-$N_3$, $C_{8a}$-$H_3$ |
| | $\nu_{61}$ | 1428 | $C_{11}$-$H_3$, $N_3$-H, $N_1$-$C_2$, $C_{9a}$-$N_{10}$, $C_{10a}$-$C_{4a}$, $C_{5a}$-$N_5$ | $\nu_{63}$ | 1383 | $C_{7a,11}$-$H_3$, $N_5$-$C_{4a}$, $C_{10a}$-$C_{4a}$, $C_7$-$C_6$ |
| | $\nu_{62}$ | 1432 | $N_3$-H, $C_6$-H, $C_4$-$N_3$, $C_{9a}$-$C_{5a}$, $C_{11}$-$H_3$, $C_{10a}$-$C_{4a}$ | - | - | - |
| 1416 | $\nu_{65}$ | 1456 | $C_{8a}$-$H_3$, $C_9$-H, $N_1$-$C_2$, $C_{10a}$-$C_{4a}$ | $\nu_{67}$ | 1415 | $C_{7a,11,8a}$-$H_3$, $C_{9a}$-$C_{5a}$, $C_{10a}$-$C_{4a}$ |
| | - | - | - | - | - | - |
| 1498 | $\nu_{71}$ | 1549 | $C_{9a}$-$N_{10}$, $C_{10a}$-$N_1$, $C_{11}$-$H_3$, $C_{5a}$-$N_5$, $C_8$-$C_9$, $C_{7a,8a}$-$H_3$ | $\nu_{71}$ | 1474 | $C_8$-$C_7$, $C_{6,9}$-H, $C_9$-$C_{9a}$, $C_6$-$C_{5a}$, $C_{9a}$-$C_{5a}$, $C_{7a,8a}$-$H_3$ |
| 1570 | $\nu_{73}$ | 1661 | $C_8$-$C_9$, $C_6$-$C_{5a}$, $C_{9,6}$-H, $C_9$-$C_{9a}$, $C_7$-$C_6$, $C_{10a}$-$N_1$, $N_5$-$C_{4a}$ | $\nu_{73}$ | 1553 | $C_2$-$O_{2'}$, $C_8$-$C_9$, $C_6$-$C_{5a}$, $C_{9,6}$-H, $N_3$-H, $C_7$-$C_6$, $C_9$-$C_{9a}$ |
| | $\nu_{74}$ | 1736 | $C_2$-$O_{2'}$, $N_3$-H, $C_4$-$O_{4'}$, $C_4$-$N_3$ (as) | $\nu_{74}$ | 1568 | $C_2$-$O_{2'}$, $N_3$-H, $C_8$-$C_9$, $C_6$-$C_{5a}$, $N_{10}$-$C_{10a}$, $C_{4a}$-$C_4$ |
| 1626 | $\nu_{75}$ | 1758 | $C_4$-$O_{4'}$, $C_2$-$O_{2'}$, $N_3$-H (s) | $\nu_{75}$ | 1644 | $C_4$-$O_{4'}$, $N_3$-H, $C_{4a}$-$C_4$ |
| **Exp. RR FMN $S_1$** | **O3LYP/cc-pVDZ** | | | **OLYP/cc-pVDZ** | | |
| | **$\nu_{\#}$** | **$S_1$offR** | **Assignment** | **$\nu_{\#}$** | **$S_1$offR** | **Assignment** |

| | | | | | | |
|---|---|---|---|---|---|---|
| 1200 | $\nu_{49}$ | 1183 | $C_6$-H, $N_5$-$C_{5a}$, $C_6$-$C_7$, $N_1$-$C_{10a}$ | $\nu_{48}$ | 1146 | $C_6$-H, $N_1$-$C_{10a}$, $C_7$-$C_6$, $N_5$-$C_{4a}$, $N_3$-$C_4$, $C_{7a,8a}$-$H_3$ |
| | - | - | - | $\nu_{50}$ | 1175 | $N_3$-H, $C_{6,9}$-H, $C_{11}$-$H_3$, $N_3$-$C_4$, $N_1$-$C_2$ |
| 1250 | $\nu_{51}$ | 1228 | $C_2$-$N_3$, $C_{4a}$-$C_4$, $N_3$-H, $C_{6,9}$-H, $C_9$-$C_{9a}$ | $\nu_{51}$ | 1215 | $N_3$-H, $N_3$-$C_2$, $C_{10a}$-$C_{4a}$, $C_{4a}$-$C_4$, $C_{6,9}$-H |
| 1338 | $\nu_{53}$ | 1290 | $C_{11}$-$H_3$, $C_{6,9}$-H, $N_{10}$-$C_{10a}$, $N_5$-$C_{4a}$, $C_8$-$C_7$ | $\nu_{53}$ | 1270 | $C_{6,9}$-H, $N_3$-H, $C_{11}$-$H_3$, $N_1$-$C_{10a}$, $N_5$-$C_{4a}$, $C_2$-$N_3$ |
| | - | - | - | - | - | - |
| 1381 | $\nu_{57}$ | 1373 | $C_{11}$-$H_3$, $N_3$-H, $C_6$-H, $C_{7a}$-$H_3$, $C_7$-$C_6$, $C_{10a}$-$C_{4a}$, $C_{9a}$-$N_{10}$ | $\nu_{57}$ | - | - |
| | $\nu_{58}$ | 1376 | $C_{8a}$-$H_3$, $C_8$-$C_7$, $C_{9a}$-$C_{5a}$, $N_3$-H | $\nu_{58}$ | - | - |
| | $\nu_{59}$ | 1385 | $C_{8a}$-$H_3$, $N_3$-H, $N_6$-$C_{5a}$, $N_3$-$C_4$, $N_1$-$C_2$, $C_8$-$C_9$ | $\nu_{59}$ | - | - |
| | $\nu_{61}$ | 1402 | $N_3$-H, $C_6$-H, $C_{9a}$-$C_{5a}$, $N_3$-$C_4$, $C_{10a}$-$C_{4a}$ | $\nu_{61}$ | 1387 | $N_3$-H, $C_{6,9}$-H, $C_{9a}$-$C_{5a}$, $C_8$-$C_7$, $C_{7a,8a}$-$H_3$ |
| 1416 | $\nu_{64}$ | 1437 | $C_{8a}$-$H_3$, $C_9$-H, $C_{11}$-$H_3$, $C_4$-$C_{4a}$, $N_1$-$C_2$ | $\nu_{64}$ | 1419 | $C_{7a}$-$H_3$, $C_{11}$-$H_3$, $C_{8a}$-$H_3$, $C_9$-H |
| | - | - | - | $\nu_{66}$ | 1420 | $C_{7a}$-$H_3$, $N_5$-$C_{4a}$, $C_{8a}$-$H_3$ |
| 1498 | $\nu_{71}$ | 1526 | $C_{6,9}$-H, $C_8$-$C_7$, $C_{5a}$-$C_{9a}$, $C_{10a}$-$C_{4a}$, $C_{7a}$-$H_3$ | $\nu_{71}$ | 1519 | $C_{6,9}$-H, $C_{9a}$-$C_{9a}$, $C_6$-$C_{5a}$, $C_8$-$C_7$ |
| 1570 | $\nu_{73}$ | 1628 | $C_{6,9}$-H, $C_8$-$C_9$, $C_{5a}$-$C_6$, $C_{10a}$-$N_1$ | $\nu_{73}$ | 1605 | $C_8$-$C_9$, $C_6$-$C_{5a}$, $C_{6,9}$-H, $C_2$-$O_{2'}$, $N_3$-H, $C_{10a}$-$N_1$ |
| | $\nu_{74}$ | 1684 | $C_2$-$O_{2'}$, $N_3$-H, $C_4$-$C_{4a}$ ~(as) | $\nu_{74}$ | 1624 | $C_2$-$O_{2'}$, $N_3$-H, $N_{10}$-$C_{10a}$, $C_8$-$C_9$, $C_6$-$C_{5a}$, $C_{4a}$-$C_4$ ~(as) |
| 1626 | $\nu_{75}$ | 1725 | $C_2$-$O_{2'}$, $N_3$-H, $C_4$-$O_{4'}$ ~(s) | $\nu_{75}$ | 1697 | $C_4$-$O_{4'}$, $N_3$-H ~(s) |
| **Exp. RR FMN $S_1$** | **OPBE/cc-pVDZ** | | | **PBE1PBE/cc-pVDZ** | | |
| | **$\nu_\#$** | **$S_1$offR** | **Assignment** | **$\nu_\#$** | **$S_1$offR** | **Assignment** |
| 1200 | $\nu_{49}$ | 1167 | $C_6$-H, $C_{7a,8a}$-$H_3$, $N_1$-$C_{10a}$, $C_7$-$C_6$, $N_5$-$C_{4a}$, $N_5$-$C_{5a}$ | $\nu_{49}$ | 1205 | $C_{6,9}$-H, $N_3$-H, $C_{5a}$-$N_5$, $C_4$-$N_3$, $C_8$-$C_7$ |
| | - | - | - | $\nu_{50}$ | 1219 | $C_{6,9}$-H, $C_{11}$-$H_3$, $C_9$-$C_{9a}$, $C_{4a}$-$C_4$, $C_4$-$N_3$, $C_{10a}$-$N_1$ |
| 1250 | $\nu_{51}$ | 1237 | $C_{6,9}$-H, $N_3$-H, $N_3$-$C_2$, $N_5$-$C_{4a}$, $C_4$-$C_{4a}$, $N_5$-$C_{5a}$ | $\nu_{51}$ | 1262 | $N_3$-H, $C_4$-$N_3$. $C_9$-H, $C_{4a}$-$C_4$, $N_1$-$C_2$, $C_{9a}$-$C_{5a}$ |
| 1338 | $\nu_{53}$ | 1282 | $C_6$-H, $N_3$-H, $C_{11}$-$H_3$, $C_{7a,8a}$-$H_3$, $N_1$-$C_{10a}$, $N_5$-$C_{4a}$, $C_6$-$C_{5a}$ | $\nu_{53}$ | 1322 | $N_{10}$-$C_{10a}$, $C_6$-$C_{5a}$, $C_{11}$-$H_3$, $N_5$-$C_{4a}$, $C_8$-$C_7$, $C_2$-$N_3$, $C_{8a}$-$H_3$ |
| | - | - | - | $\nu_{56}$ | 1373 | $N_3$-H, $C_{5a}$-$N_5$, $C_{10a}$-$N_1$, $C_{7a}$-$H_3$, $C_{11}$-$H_3$ |
| 1381 | - | - | - | - | - | - |
| | $\nu_{58}$ | 1359 | $C_{11}$-$H_3$, $N_3$-H, $C_{8a}$-$H_3$, $C_6$-H, $N_{10}$-$C_{9a}$ | $\nu_{58}$ | 1392 | $N_3$-H, $C_{11}$-$H_3$, $C_6$-H, $C_{7a}$-$H_3$, $C_7$-$C_6$ |
| | $\nu_{62}$ | 1394 | $C_{7a,8a}$-$H_3$, $C_{11}$-$H_3$, $C_6$-H, $N_3$-$C_4$, $N_3$-H, $C_{9a}$-$C_{5a}$, $C_{10a}$-$C_{4a}$ | $\nu_{59}$ | 1404 | $C_{11}$-$H_3$, $C_{8a}$-$H_3$, $C_8$-$C_7$, $N_3$-H, $C_6$-$C_{5a}$ |
| | $\nu_{63}$ | 1408 | $C_{7a,8a}$-$H_3$, $C_{11}$-$H_3$, $C_8$-$C_7$, $C_{9a}$-$C_{5a}$, $C_{10a}$-$C_{4a}$, $C_6$-H | - | - | - |
| 1416 | - | - | - | - | - | - |
| | $\nu_{66}$ | 1422 | $C_{7a}$-$H_3$, $C_{11}$-$H_3$, $C_{9a}$-$C_{5a}$, $C_{10a}$-$C_{4a}$, $N_1$-$C_2$ | $\nu_{65}$ | 1451 | $C_{8a}$-$H_3$, $C_9$-H, $N_1$-$C_2$, $C_{11}$-$H_3$ |
| 1498 | $\nu_{71}$ | 1535 | $C_{6,9}$-H, $C_{9a}$-$C_{9a}$, $C_6$-$C_{5a}$, $C_8$-$C_7$ | $\nu_{71}$ | 1548 | $C_8$-$C_7$, $C_{9a}$-$C_{5a}$, $C_{6,9}$-H, $C_{7a}$-$H_3$ |
| 1570 | $\nu_{73}$ | 1626 | $C_8$-$C_9$, $C_6$-$C_{5a}$, $C_{6,9}$-H, $C_2$-$O_{2'}$, $N_3$-H, $C_{10a}$-$N_1$ | $\nu_{73}$ | 1659 | $C_8$-$C_9$, $C_6$-$C_{5a}$, $C_{6,9}$-H, $N_5$-$C_{4a}$ |
| | $\nu_{74}$ | 1652 | $C_2$-$O_{2'}$, $N_3$-H, $N_{10}$-$C_{10a}$, $C_8$-$C_9$, $C_6$-$C_{5a}$, $C_{4a}$-$C_4$ ~(as) | $\nu_{74}$ | 1738 | $C_2$-$O_{2'}$, $N_3$-H, $C_4$-$O_{4'}$ (as) |
| 1626 | $\nu_{75}$ | 1725 | $C_4$-$O_{4'}$, $N_3$-H ~(s) | $\nu_{75}$ | 1759 | $C_4$-$O_{4'}$, $C_2$-$O_{2'}$, $N_3$-H (s) |
| **Exp. RR FMN $S_1$** | **PW6B95D3/cc-pVDZ** | | | **revTPSSh/aug-cc-pVDZ** | | |
| | **$\nu_\#$** | **$S_1$offR** | **Assignment** | **$\nu_\#$** | **$S_1$offR** | **Assignment** |
| 1200 | $\nu_{49}$ | 1208 | $C_6$-H, $N_3$-H, $C_8$-$C_7$, $C_{5a}$-$N_5$, $C_7$-$C_6$, $C_{5a}$-$N_5$, $C_4$-$N_3$ | $\nu_{51}$ | 1354 | $C_2$-$N_3$, $C_{10a}$-$C_{4a}$, $N_3$-H, $C_{6,9}$-H, $C_4$-$N_3$, $C_{9a}$-$C_{5a}$ |

| | | | | | | |
|---|---|---|---|---|---|---|
| | $\nu_{50}$ | 1222 | $C_{6,9}$-H, $C_{4a}$-$C_4$, $N_5$-$C_{4a}$, $N_{10}$-$C_{10a}$, $C_{11}$-$H_3$, $C_9$-$C_{9a}$, $C_{11}$-$H_3$, $C_{10a}$-$N_1$ | - | - | - |
| 1250 | $\nu_{51}$ | 1257 | $C_2$-$N_3$, $N_3$-H, $C_{4a}$-$C_4$, $C_{5a}$-$N_5$, $C_9$-H, $N_1$-$C_2$, $C_{9a}$-$C_{5a}$, $C_7$-$C_6$ | $\nu_{52}$ | 1382 | $C_9$-H, $C_8$-$C_7$, $C_{5a}$-$N_5$, $C_6$-H, $C_{7a}$-$H_3$, $C_{10a}$-$C_{4a}$ |
| 1338 | $\nu_{54}$ | 1339 | $C_{5a}$-$N_5$, $C_9$-$C_{9a}$, $N_{10}$-$C_{10a}$, $C_{6,9}$-H, $C_8$-$C_7$, $C_2$-$N_3$ | - | - | - |
| | $\nu_{56}$ | 1378 | $C_{7a}$-$H_3$, $N_3$-H, $C_{10a}$-$N_1$, $C_{5a}$-$N_5$, $C_8$-$C_7$ | $\nu_{56}$ | 1491 | $N_3$-H, $C_6$-H, $C_6$-$C_{5a}$, $C_{9a}$-$N_{10}$, $C_7$-$C_6$, $C_8$-$C_9$, $C_{10a}$-$C_{4a}$, $C_4$-$N_3$ |
| 1381 | $\nu_{58}$ | 1396 | $N_3$-H, $C_{11,7a}$-$H_3$, $C_6$-H, $C_7$-$C_6$, $C_{9a}$-$N_{10}$, $C_{5a}$-$N_5$, $C_{10a}$-$C_{4a}$ | $\nu_{57}$ | | |
| | $\nu_{59}$ | 1405 | $C_{11,7a,8a}$-$H_3$, $C_8$-$C_7$, $C_{9a}$-$C_{5a}$, $N_3$-H, $C_6$-$C_{5a}$ | $\nu_{58}$ | 1516 | $C_{7a,8a}$-$H_3$, $C_8$-$C_7$, $C_{5a}$-$C_6$, $C_2$-$N_3$, $N_3$-$C_4$ |
| | $\nu_{60}$ | 1425 | $C_{11,7a,8a}$-$H_3$, $C_{9a}$-$N_{10}$, $N_5$-$C_{4a}$, $C_{10a}$-$N_1$, $C_8$-$C_7$, $N_3$-H | $\nu_{59}$ | 1522 | $N_3$-H, $C_6$-H, $C_{9a}$-$C_{5a}$, $C_{4a}$-$C_{10a}$, $C_9$-$C_{9a}$, $C_{10a}$-$C_{4a}$, $C_4$-$N_3$ |
| | $\nu_{62}$ | 1432 | $N_3$-H, $C_{9a}$-$C_{5a}$, $C_{7a,11}$-$H_3$, $C_6$-H, $C_{10a}$-$N_1$, $C_7$-$C_6$, $C_4$-$N_3$ | $\nu_{60}$ | 1526 | $C_{7a}$-$H_3$, $C_{9a}$-$C_{5a}$, $C_{10a}$-$C_{4a}$, $C_2$-$N_3$, $N_3$-H |
| 1416 | $\nu_{65}$ | 1455 | $C_{8a,11}$-$H_3$, $C_2$-$N_1$, $C_{10a}$-$C_4$ | $\nu_{63}$ | 1574 | $N_5$-$C_{4a}$, $C_{10a}$-$C_{4a}$, $C_4$-$N_3$, $C_{10a}$-$N_1$, $C_{11,7a}$-$H_3$, $C_8$-$C_7$ |
| | $\nu_{68}$ | 1475 | $N_5$-$C_{4a}$, $C_{11}$-$H_3$, $C_{10a}$-$C_{4a}$, $C_7$-$C_6$, $C_2$-$N_3$, $C_{7a}$-$H_3$ | - | - | - |
| 1498 | $\nu_{71}$ | 1549 | $C_8$-$C_7$, $C_9$-$C_{9a}$, $C_{9a}$-$C_{5a}$, $C_6$-$C_{5a}$, $C_{6,9}$-H, $C_{7a}$-$H_3$, $C_{10a}$-$C_{4a}$ | $\nu_{71}$ | 1653 | $C_8$-$C_7$, $C_{9a}$-$C_{5a}$, $C_{6,9}$-H, $C_{7a}$-$H_3$, $C_6$-$C_{5a}$, $C_9$-$C_{9a}$, $C_{10a}$-$C_{4a}$ |
| 1570 | $\nu_{73}$ | 1659 | $C_8$-$C_9$, $C_6$-$C_{5a}$, $C_{6,9}$-H, $C_7$-$C_6$, $N_{10}$-$C_{9a}$, $N_5$-$C_{4a}$, $C_{10a}$-$N_1$ | $\nu_{73}$ | 1770 | $C_8$-$C_9$, $C_6$-$C_{5a}$, $C_{6,9}$-H, $C_7$-$C_6$, $C_9$-$C_{9a}$, $C_2$-$O_{2'}$, $C_4$-$O_{4'}$, $C_{10a}$-$N_1$ (as) |
| | $\nu_{74}$ | 1737 | $C_2$-$O_{2'}$, $N_3$-H, $C_4$-$O_{4'}$ (as), $C_{4a}$-$C_{10a}$ | $\nu_{74}$ | 1797 | $C_2$-$O_{2'}$, $N_3$-H, $C_2$-$N_3$, $C_6$-$C_{5a}$, $C_8$-$C_9$ |
| 1626 | $\nu_{75}$ | 1757 | $C_4$-$O_{4'}$, $C_2$-$O_{2'}$ (s), $N_3$-H, $C_{4a}$-$C_{10a}$ | $\nu_{75}$ | 1817 | $C_4$-$O_{4'}$, $N_3$-H, $C_{4a}$-$C_4$, $C_6$-$C_{5a}$, $C_2$-$O_{2'}$ (s), $C_8$-$C_9$ |
| **Exp. RR FMN $S_1$** | **revTPSS/aug-cc-pVDZ** | | | **SOGGA11/cc-pVDZ** | | |
| | **ν#** | **$S_1$offR** | **Assignment** | **ν#** | **$S_1$offR** | **Assignment** |
| 1200 | $\nu_{51}$ | 1199 | $C_2$-$N_3$, $C_{10}$-$C_{4a}$, $N_3$-H, $C_{6,9}$-H, $C_9$-$C_{9a}$, $C_{9a}$-$C_{5a}$, $N_5$-$C_4$, $N_1$-$C_2$ | $\nu_{49}$ | 1229 | $C_{6,9}$-H, $N_3$-H, $C_{5a}$-$N_5$, $C_4$-$N_3$, $C_7$-$C_6$, $C_8$-$C_9$ |
| | - | - | - | $\nu_{50}$ | 1240 | $C_{6,9}$-H, $C_{11}$-$H_3$, $C_{4a}$-$C_4$, $C_9$-$C_{9a}$, $N_{10}$-$C_{10a}$ |
| 1250 | $\nu_{53}$ | 1265 | $C_6$-H, $C_{11}$-$H_3$, $C_{10a}$-$C_{4a}$, $C_{9a}$-$N_{10}$, $N_3$-H, $C_{9a}$-$C_{5a}$, $C_8$-$C_7$ | $\nu_{51}$ | 1286 | $N_3$-H, $C_6$-H, $C_2$-$N_3$, $C_{4a}$-$C_4$, $N_1$-$C_2$, $N_5$-$C_{4a}$, $C_{5a}$-$C_{9a}$ |
| 1338 | - | - | - | $\nu_{53}$ | 1340 | $N_{10}$-$C_{10a}$, $N_5$-$C_{4a}$, $C_{9a}$-$C_{5a}$, $C_8$-$C_7$, $N_5$-$C_{4a}$, $C_{11}$-$H_3$, $C_2$-$N_3$ |
| | $\nu_{56}$ | 1340 | $N_3$-H, $C_6$-H, $C_{11}$-$H_3$, $C_{9a}$-$N_{10}$, $C_7$-$C_6$, $C_8$-$C_9$, $C_6$-$C_{5a}$, $N_1$-$C_2$ | $\nu_{54}$ | 1361 | $C_{5a}$-$N_5$, $C_{10a}$-$C_{4a}$, $N_3$-H, $C_9$-$C_{9a}$, $N_{10}$-$C_{10a}$, $C_2$-$N_3$, $C_{6,9}$-H, $C_{7a}$-$H_3$ |
| 1381 | $\nu_{57}$ | 1356 | $N_3$-H, $C_{10a}$-$N_1$, $C_6$-H, $N_{11}$-$H_3$, $N_1$-$C_2$, $C_{9a}$-$C_{5a}$, $N_5$-$C_{4a}$, $C_{9a}$-$N_{10}$ | $\nu_{57}$ | 1414 | $C_{11,7a}$-$H_3$, $C_{9a}$-$N_{10}$, $C_6$-$C_{5a}$, $C_{10a}$-$N_1$, $C_7$-$C_6$ |
| | $\nu_{58}$ | 1366 | $N_3$-H, $C_{10a}$-$N_{10}$, $N_5$-$C_{4a}$, $C_{10a}$-$N_1$, $C_7$-$C_6$, $C_6$-H, $C_8$-$C_7$, $C_{7a}H_3$ | $\nu_{58}$ | 1422 | $C_{11,7a,8a}$-$H_3$, $C_8$-$C_7$, $C_6$-$C_{5a}$, $C_{9a}$-$C_{5a}$, $N_3$-H |
| | $\nu_{59}$ | 1371 | $N_3$-H, $C_6$-H, $C_9$-$C_{9a}$, $C_{9a}$-$C_{5a}$, $C_4$-$N_3$, $C_8$-$C_7$, $C_{10a}$-$C_{4a}$ | $\nu_{59}$ | 1427 | $N_3$-H, $C_4$-$N_3$, $C_{10a}$-$C_{4a}$, $C_6$-H, $C_{7a}$-$H_3$, $C_{9a}$-$C_{5a}$ |
| | $\nu_{60}$ | 1377 | $C_{7a,8a}$-$H_3$, $C_8$-$C_7$, $C_{9a}$-$C_{5a}$, $N_5$-$C_{4a}$ | $\nu_{60}$ | 1445 | $C_{11,7a,8a}$-$H_3$, $N_{10}$-$C_{10a}$, $C_{10a}$-$C_{4a}$, $N_5$-$C_{5a}$ |
| 1416 | $\nu_{63}$ | 1420 | $C_{11,7a}$-$H_3$, $C_7$-$C_6$, $C_{5a}$-$N_5$, $C_9$-$C_{9a}$, $N_{10}$-$C_{10a}$, $N_5$-$C_{4a}$ | $\nu_{65}$ | 1477 | $C_{7a,11}$-$H_3$, $N_1$-$C_2$, $C_{10a}$-$C_{4a}$, $C_6$-$C_{5a}$ |
| | - | - | - | $\nu_{68}$ | 1498 | $N_5$-$C_{4a}$, $C_{10a}$-$N_1$, $C_{9a}$-$N_{10}$, $C_{11}$-$H_3$, $N_3$-H, $C_2$-$N_3$, $C_{7a,8a}$-$H_3$ |
| 1498 | $\nu_{71}$ | 1506 | $C_8$-$C_7$, $C_{9a}$-$C_{5a}$, $C_{6,9}$-H, $C_{7a}$-$H_3$, $C_6$-$C_{5a}$, $C_9$-$C_{9a}$, $C_{10a}$-$C_{4a}$ | $\nu_{70}$ | 1561 | $C_8$-$C_7$, $C_9$-$C_{9a}$, $C_{9a}$-$C_{5a}$, $C_6$-$C_{5a}$, $C_{10a}$-$N_1$, $C_{6,9}$-H, $C_{7a}$-$H_3$ |
| 1570 | $\nu_{73}$ | 1569 | $C_2$-$O_{2'}$, $N_3$-H, $C_2$-$N_3$, $C_6$-$C_{5a}$, $C_8$-$C_9$ | $\nu_{73}$ | 1679 | $C_8$-$C_9$, $C_6$-$C_{5a}$, $C_{6,9}$-H, $C_7$-$C_6$, $N_{10}$-$C_{9a}$, $N_5$-$C_{4a}$, $C_{10a}$-$N_1$ |

| | | | | | | |
|---|---|---|---|---|---|---|
| | $\nu_{74}$ | 1594 | (s) $C_8$-$C_9$, $C_6$-$C_{5a}$, $C_{6,9}$-H, $C_7$-$C_6$, $C_9$-$C_{9a}$, $C_2$-$O_{2'}$, $C_4$-$O_{4'}$, $C_{10a}$-$N_1$ | $\nu_{74}$ | 1777 | $C_2$-$O_{2'}$, $N_3$-H, $C_4$-$O_{4'}$ (as), $C_{4a}$-$C_{10a}$ |
| 1626 | $\nu_{75}$ | 1627 | $C_4$-$O_{4'}$, $N_3$-H, $C_{4a}$-$C_4$, $C_6$-$C_{5a}$, $C_8$-$C_9$ | $\nu_{75}$ | 1794 | $C_4$-$O_{4'}$, $C_2$-$O_{2'}$ (s), $C_{4a}$-$C_{10a}$ |
| **Exp. RR FMN $S_1$** | **SOGGA11x/cc-pVDZ** | | | **tHCTHhyb/cc-pVDZ** | | |
| | **$\nu_{\#}$** | **$S_1$offR** | **Assignment** | **$\nu_{\#}$** | **$S_1$offR** | **Assignment** |
| 1200 | $\nu_{49}$ | 1229 | $C_{6,9}$-H, $N_3$-H, $C_{5a}$-$N_5$, $C_4$-$N_3$, $C_7$-$C_6$, $C_8$-$C_9$ | $\nu_{49}$ | 1177 | $C_6$-H, $C_{5a}$-$N_5$, $C_{11}$-$H_3$, $N_3$-H, $C_8$-$C_7$ |
| | $\nu_{50}$ | 1240 | $C_{6,9}$-H, $C_{11}$-$H_3$, $C_{4a}$-$C_4$, $C_9$-$C_{9a}$, $N_{10}$-$C_{10a}$ | $\nu_{50}$ | 1187 | $C_9$-H, $C_4$-$N_3$, $N_3$-H, $C_{11}$-$H_3$, $C_{4a}$-$C_4$, $C_{10a}$-$N_1$, $C_{9a}$-$N_{10}$ |
| 1250 | $\nu_{51}$ | 1286 | $N_3$-H, $C_6$-H, $C_2$-$N_3$, $C_{4a}$-$C_4$, $N_1$-$C_2$, $N_5$-$C_{4a}$, $C_{5a}$-$C_{9a}$ | $\nu_{51}$ | 1220 | $C_2$-$N_3$, $N_3$-H, $C_9$-H, $C_{4a}$-$C_4$, $N_5$-$C_{4a}$, $C_{9a}$-$C_{5a}$ |
| 1338 | $\nu_{53}$ | 1340 | $N_{10}$-$C_{10a}$, $N_5$-$C_{4a}$, $C_{9a}$-$C_{5a}$, $C_8$-$C_7$, $N_5$-$C_{4a}$, $C_{11}$-$H_3$, $C_2$-$N_3$ | - | - | - |
| | $\nu_{54}$ | 1361 | $C_{5a}$-$N_5$, $C_{10a}$-$C_{4a}$, $N_3$-H, $C_9$-$C_{9a}$, $N_{10}$-$C_{10a}$, $C_2$-$N_3$, $C_{6,9}$-H, $C_{7a}$-$H_3$ | $\nu_{56}$ | 1355 | $C_{7a}$-$H_3$, $C_8$-$C_7$, $C_9$-H, $C_{11}$-$H_3$ |
| 1381 | $\nu_{57}$ | 1414 | $C_{11,7a}$-$H_3$, $C_{9a}$-$N_{10}$, $C_6$-$C_{5a}$, $C_{10a}$-$N_1$, $C_7$-$C_6$ | $\nu_{57}$ | 1367 | $N_3$-H, $C_6$-H, $C_{11,7a}$-$H_3$, $C_6$-$C_{5a}$, $C_{9a}$-$C_{5a}$, $C_{9a}$-$N_{10}$, $C_{10a}$-$C_{4a}$ |
| | $\nu_{58}$ | 1422 | $C_{11,7a,8a}$-$H_3$, $C_8$-$C_7$, $C_6$-$C_{5a}$, $C_{9a}$-$C_{5a}$, $N_3$-H | $\nu_{58}$ | 1371 | $C_{8a}$-$H_3$, $C_8$-$C_9$, $C_{9a}$-$C_{5a}$ |
| | $\nu_{59}$ | 1427 | $N_3$-H, $C_4$-$N_3$, $C_{10a}$-$C_{4a}$, $C_6$-H, $C_{7a}$-$H_3$, $C_{9a}$-$C_{5a}$ | $\nu_{59}$ | 1379 | $C_{8a}$-$H_3$, $N_3$-H, $C_{9a}$-$C_{5a}$, $C_6$-$C_{5a}$, $C_8$-$C_7$, $N_1$-$C_2$ |
| | $\nu_{60}$ | 1445 | $C_{11,7a,8a}$-$H_3$, $N_{10}$-$C_{10a}$, $C_{10a}$-$C_{4a}$, $N_5$-$C_{5a}$ | $\nu_{61}$ | 1394 | $N_3$-H, $N_3$-$C_4$, $C_6$-H, $C_{9a}$-$C_{5a}$, $C_{10a}$-$C_{4a}$ |
| 1416 | $\nu_{65}$ | 1477 | $C_{7a,11}$-$H_3$, $N_1$-$C_2$, $C_{10a}$-$C_{4a}$, $C_6$-$C_{5a}$ | $\nu_{64}$ | 1432 | $N_5$-$C_{4a}$, $C_{7a}$-$H_3$, $C_{10a}$-$C_{4a}$, $C_{10a}$-$N_1$ |
| | $\nu_{68}$ | 1498 | $N_5$-$C_{4a}$, $C_{10a}$-$N_1$, $C_{9a}$-$N_{10}$, $C_{11}$-$H_3$, $N_3$-H, $C_2$-$N_3$, $C_{7a,8a}$-$H_3$ | $\nu_{65}$ | 1433 | $C_{8a,11,7a}$-$H_3$, $C_9$-H, $N_{10}$-$C_{10a}$, $N_1$-$C_2$, $C_4$-$N_3$, $N_3$-H |
| 1498 | $\nu_{70}$ | 1561 | $C_8$-$C_7$, $C_9$-$C_{9a}$, $C_{9a}$-$C_{5a}$, $C_6$-$C_{5a}$, $C_{10a}$-$N_1$, $C_{6,9}$-H, $C_{7a}$-$H_3$ | $\nu_{71}$ | 1515 | $C_8$-$C_7$, $C_{9a}$-$C_{5a}$, $C_9$-H, $C_{7a}$-$H_3$, $C_9$-$C_{9a}$, $C_6$-$C_{5a}$, $C_{10a}$-$C_{4a}$ |
| 1570 | $\nu_{73}$ | 1679 | $C_8$-$C_9$, $C_6$-$C_{5a}$, $C_{6,9}$-H, $C_7$-$C_6$, $N_{10}$-$C_{9a}$, $N_5$-$C_{4a}$, $C_{10a}$-$N_1$ | $\nu_{73}$ | 1618 | $C_8$-$C_9$, $C_6$-$C_{5a}$, $C_{6,9}$-H, $C_9$-$C_{9a}$, $C_7$-$C_6$, $C_{10a}$-$N_1$, $C_{4a}$-$N_5$ |
| | $\nu_{74}$ | 1777 | $C_2$-$O_{2'}$, $N_3$-H, $C_4$-$O_{4'}$ (as), $C_{4a}$-$C_{10a}$ | $\nu_{74}$ | 1687 | $C_2$-$O_{2'}$, $N_3$-H, $C_{10a}$-$N_1$, $N_3$-$C_4$ |
| 1626 | $\nu_{75}$ | 1794 | $C_4$-$O_{4'}$, $C_2$-$O_{2'}$ (s), $C_{4a}$-$C_{10a}$ | $\nu_{75}$ | 1721 | $C_4$-$O_{4'}$, $N_3$-H, $C_{4a}$-$C_4$, $C_2$-$O_{2'}$ (s) |
| **Exp. RR FMN $S_1$** | **TPSSh/cc-pVDZ** | | | **TPSSTPSS/cc-pVDZ** | | |
| | **$\nu_{\#}$** | **$S_1$offR** | **Assignment** | **$\nu_{\#}$** | **$S_1$offR** | **Assignment** |
| 1200 | $\nu_{49}$ | 1173 | $C_6$-H, $C_{10a}$-$N_1$, $C_4$-$N_3$, $C_{5a}$-$N_5$, $C_7$-$C_6$, $N_5$-$C_{4a}$ | $\nu_{51}$ | 1221 | $C_9$-H, $C_2$-$N_3$, $C_{10a}$-$C_{4a}$, $C_4$-$N_3$, $C_6$-$C_{5a}$, $C_9$-$C_{9a}$ |
| | $\nu_{50}$ | 1184 | $C_{6,9}$-H, $C_{11}$-$H_3$, $C_4$-$N_3$, $N_3$-H, $C_8$-$C_7$, $C_{10a}$-$C_{4a}$ | $\nu_{52}$ | 1246 | $C_2$-$O_{2'}$, $N_3$-H, $C_{6,9}$-H, $C_9$-$N_{10}$, $C_8$-$C_9$, $N_1$-$C_2$ |
| 1250 | $\nu_{51}$ | 1220 | $C_2$-$N_3$, $C_{10a}$-$C_{4a}$, $N_3$-H, $C_{6,9}$-H, $C_4$-$N_3$, $C_{9a}$-$C_{5a}$ | $\nu_{54}$ | 1276 | $N_3$-H, $C_2$-$O_{2'}$, $C_{6,9}$-H, $C_{4a}$-$C_4$, $N_{10}$-$C_{10a}$, $C_{5a}$-$C_{9a}$, $C_{11}$-$H_3$ |
| 1338 | - | - | - | - | - | - |
| | $\nu_{56}$ | 1369 | $N_3$-H, $C_6$-H, $C_{11}$-$H_3$, $C_{9a}$-$N_{10}$, $C_7$-$C_6$, $C_8$-$C_9$, $C_{10a}$-$C_{4a}$, $C_4$-$N_3$ | $\nu_{56}$ | 1330 | $C_{5a}$-$N_5$, $C_9$-$C_{9a}$, $C_8$-$C_7$, $N_3$-H, $C_9$-H, $N_{10}$-$C_{10a}$ |
| 1381 | $\nu_{57}$ | 1375 | $N_3$-H, $C_{7a}$-$H_3$, $C_8$-$C_7$, $C_{9a}$-$C_{5a}$, $C_4$-$N_3$, $C_{6,9}$-H | $\nu_{57}$ | 1353 | $N_3$-H, $C_{10a}$-$N_1$, $C_6$-H, $N_{11}$-$H_3$, $N_1$-$C_2$, $C_{9a}$-$C_{5a}$, $N_5$-$C_{4a}$, $C_{9a}$-$N_{10}$ |
| | $\nu_{58}$ | 1383 | $C_{7a,11}$-$H_3$, $C_{10a}$-$N_1$, $C_{9a}$-$N_{10}$, $C_2$-$N_3$, $N_5$-$C_{4a}$ | $\nu_{58}$ | 1367 | $C_8$-$C_9$, $C_6$-$C_{5a}$, $C_{9a}$-$C_{5a}$, $C_{11}$-$H_3$, $N_3$-H, $C_{10a}$-$C_{4a}$ |
| | $\nu_{60}$ | 1396 | $N_3$-H, $C_{8a}$-$H_3$, $C_6$-H, $C_{9a}$-$C_{5a}$, $N_4$-$C_{4a}$, $C_4$-$N_3$ | - | - | - |
| | $\nu_{61}$ | 1401 | $C_{8a}$-$H_3$, $C_{9a}$-$C_{5a}$, $N_3$-H, $C_6$-H | $\nu_{61}$ | 1412 | $N_3$-H, $C_2$-$N_3$, $C_{11}$-$H_3$, $N_5$-$C_{4a}$ |
| 1416 | $\nu_{63}$ | 1435 | $C_{7a,11}$-$H_3$, $N_5$-$C_{4a}$, $C_8$-$C_7$, $C_{10a}$-$C_{4a}$, $C_4$-$N_3$ | - | - | - |

| | | | | | | |
|---|---|---|---|---|---|---|
| | - | - | - | $\nu_{64}$ | 1446 | $N_3$-H, $C_{7a,8a,11}$-$H_3$, $C_7$-$C_6$, $C_8$-$C_9$, $C_{10a}$-$C_{4a}$, $C_2$-$N_3$ |
| 1498 | $\nu_{71}$ | 1526 | $C_8$-$C_7$, $C_{9a}$-$C_{5a}$, $C_{9,6}$-H, $C_9$-$C_{9a}$, $C_6$-$C_{5a}$, $C_{7a,8a}$-$H_3$ | $\nu_{71}$ | 1518 | $C_8$-$C_7$, $C_9$-$C_{9a}$, $C_6$-$C_{5a}$, $C_{6,9}$-H, $N_{10}$-$C_{10a}$, $N_5$-$C_{4a}$, $C_{10a}$-$N_1$, $C_{11}$-$H_3$ |
| 1570 | $\nu_{73}$ | 1621 | $C_{6,9}$-H, $C_8$-$C_9$, $C_6$-$C_{5a}$, $C_7$-$C_6$, $C_9$-$C_{9a}$, $C_{10a}$-$N_1$, $C_2$-$O_{2'}$ | $\nu_{73}$ | 1582 | $C_2$-$O_{2'}$, $N_1$-$C_2$, $N_3$-H, $C_2$-$N_3$, $C_4$-$O_{4'}$, $C_6$-$C_{5a}$, $C_8$-$C_9$ |
| | $\nu_{74}$ | 1658 | $C_2$-$O_{2'}$, $N_3$-H $C_{10a}$-$N_1$ | $\nu_{74}$ | 1614 | $C_8$-$C_9$, $C_6$-$C_{5a}$, $C_{6,9}$-H, $C_7$-$C_6$, $C_9$-$C_{9a}$, $C_{10a}$-$N_1$ (as) |
| 1626 | $\nu_{75}$ | 1706 | $C_4$-$O_{4'}$, $N_3$-H, $C_{4a}$-$C_4$, $C_2$-$O_{2'}$ (s) | $\nu_{75}$ | 1695 | $C_4$-$O_{4'}$, $C_{4a}$-$C_4$, $C_2$-$O_{2'}$ (s), $N_1$-$C_2$, $N_3$-H |
| **Exp. RR FMN $S_1$** | **TPSSTPSS/aug-cc-pVDZ** | | | **VSXC/cc-pVDZ** | | |
| | **ν#** | **$S_1$offR** | **Assignment** | **ν#** | **$S_1$offR** | **Assignment** |
| 1200 | $\nu_{51}$ | 1191 | $C_2$-$N_3$, $C_{10}$-$C_{4a}$, $N_3$-H, $C_{6,9}$-H, $C_9$-$C_{9a}$, $C_{9a}$-$C_{5a}$, $N_5$-$C_4$, $N_1$-$C_2$ | $\nu_{51}$ | 1244 | $C_9$-H, $C_{4a}$-$C_4$, $N_3$-H, $C_6$-$C_{5a}$, $C_2$-$N_3$, $C_{9a}$-$C_{5a}$, $C_{11}$-$H_3$ |
| | - | - | - | $\nu_{52}$ | 1259 | $N_3$-H, $C_2$-$O_{2'}$, $C_{6,9}$-H, $C_{9a}$-$N_{10}$, $C_{10a}$-$N_1$ |
| 1250 | $\nu_{53}$ | 1261 | $C_6$-H, $C_{11}$-$H_3$, $C_{10a}$-$C_{4a}$, $C_{9a}$-$N_{10}$, $N_3$-H, $C_{9a}$-$C_{5a}$, $C_8$-$C_7$ | $\nu_{53}$ | 1282 | $N_5$-$C_{4a}$, $C_{6,9}$-H, $C_{9a}$-$C_{5a}$, $C_{10a}$-$N_1$, $C_8$-$C_7$ |
| 1338 | - | - | - | $\nu_{55}$ | 1328 | $C_{10a}$-$N_1$, $C_{9a}$-$N_{10}$, $C_{4a}$-$C_4$, $C_{5a}$-$N_5$, $C_8$-$C_7$, $C_{11}$-$H_3$, $N_3$-H |
| | $\nu_{56}$ | 1336 | $C_6$-H, $N_3$-H, $C_{11}$-$H_3$, $C_{9a}$-$N_{10}$, $C_7$-$C_6$, $C_8$-$C_9$, $C_6$-$C_{5a}$, $N_1$-$C_2$ | $\nu_{56}$ | 1342 | $C_{5a}$-$N_5$, $N_3$-H, $C_9$-H, $C_{7a}$-$H_3$, $C_9$-$C_{9a}$, $C_{4a}$-$C_4$, $C_8$-$C_7$ |
| 1381 | $\nu_{57}$ | 1351 | $N_3$-H, $C_{10a}$-$N_1$, $C_6$-H, $N_{11}$-$H_3$, $N_1$-$C_2$, $C_{9a}$-$C_{5a}$, $N_5$-$C_{4a}$, $C_{9a}$-$N_{10}$ | - | - | - |
| | $\nu_{59}$ | 1367 | $N_3$-H, $C_6$-H, $C_9$-$C_{9a}$, $C_{9a}$-$C_{5a}$, $C_4$-$N_3$, $C_8$-$C_7$, $C_{10a}$-$C_{4a}$ | $\nu_{58}$ | 1361 | $C_{9a}$-$C_{5a}$, $N_{10}$-$C_{10a}$, $C_{11,7a,8a}$-$H_3$, $N_5$-$C_{4a}$, $N_3$-H |
| | $\nu_{60}$ | 1371 | $C_{7a,8a}$-$H_3$, $C_8$-$C_7$, $C_{9a}$-$C_{5a}$, $N_5$-$C_{4a}$ | $\nu_{59}$ | 1366 | $C_{11,7a,8a}$-$H_3$, $N_3$-H, $C_{10a}$-$C_{4a}$, $C_{9a}$-$C_{5a}$ |
| | - | - | - | $\nu_{63}$ | 1417 | $C_{11,7a}$-$H_3$, $C_{9a}$-$C_{5a}$, $N_3$-H, $C_{10a}$-$N_1$ |
| 1416 | $\nu_{63}$ | 1414 | $C_{11,7a}$-$H_3$, $C_7$-$C_6$, $C_{5a}$-$N_5$, $C_9$-$C_{9a}$, $N_{10}$-$C_{10a}$, $N_5$-$C_{4a}$ | $\nu_{66}$ | 1442 | $C_{11,7a,8a}$-$H_3$, $N_3$-H, $C_2$-$N_3$, $N_{10}$-$C_{10a}$ |
| | - | - | - | $\nu_{68}$ | 1479 | $N_3$-H, $C_2$-$N_3$, $C_{10a}$-$C_{4a}$, $C_{6,9}$-H, $C_8$-$C_9$, $C_{11}$-$H_3$ |
| 1498 | $\nu_{71}$ | 1501 | $C_8$-$C_7$, $C_{9a}$-$C_{5a}$, $C_{6,9}$-H, $C_{7a}$-$H_3$, $C_6$-$C_{5a}$, $C_9$-$C_{9a}$, $C_{10a}$-$C_{4a}$ | $\nu_{71}$ | 1536 | $C_8$-$C_7$, $C_9$-$C_{9a}$, $C_{9a}$-$C_{5a}$, $C_6$-$C_{5a}$, $C_{10a}$-$N_1$, $C_{6,9}$-H, $C_{7a}$-$H_3$ |
| 1570 | $\nu_{73}$ | 1566 | $C_4$-$O_{4'}$, $N_3$-H, $C_2$-$N_3$, $C_6$-$C_{5a}$, $C_8$-$C_9$ | $\nu_{73}$ | 1612 | $C_8$-$C_9$, $C_6$-$C_{5a}$, $C_{6,9}$-H, $C_7$-$C_6$, $N_{10}$-$C_{9a}$, $N_5$-$C_{4a}$, $C_{10a}$-$N_1$ |
| | $\nu_{74}$ | 1590 | (s) $C_8$-$C_9$, $C_6$-$C_{5a}$, $C_{6,9}$-H, $C_7$-$C_6$, $C_9$-$C_{9a}$, $C_2$-$O_{2'}$, $C_4$-$O_{4'}$, $C_{10a}$-$N_1$ | $\nu_{74}$ | 1635 | $C_2$-$O_{2'}$, $N_3$-H, $C_4$-$O_{4'}$ (as), $C_{4a}$-$C_{10a}$ |
| 1626 | $\nu_{75}$ | 1623 | $C_2$-$O_{2'}$, $N_3$-H, $C_{4a}$-$C_4$, $C_6$-$C_{5a}$, $C_8$-$C_9$ | $\nu_{75}$ | 1725 | $C_4$-$O_{4'}$, $C_2$-$O_{2'}$ (s), $C_{4a}$-$C_{10a}$, $C_{10a}$-$N_1$ |
| **Exp. RR FMN $S_1$** | **VSXC/aug-cc-pVDZ** | | | **wB97XD/cc-pVDZ** | | |
| | **ν#** | **$S_1$offR** | **Assignment** | **ν#** | **$S_1$offR** | **Assignment** |
| 1200 | $\nu_{49}$ | 1150 | $C_6$-H, $C_{11}$-$H_3$, $C_{5a}$-$N_5$, $C_{10a}$-$N_1$, $C_7$-$C_6$, $C_{4a}$-$C_4$ | $\nu_{50}$ | 1232 | $C_6$-H, $C_{11}$-$H_3$, $C_9$-$C_{9a}$, $C_{10a}$-$C_{4a}$, $C_{4a}$-$C_4$ |
| | $\nu_{51}$ | 1204 | $C_2$-$N_3$, $N_3$-H, $C_{10a}$-$C_{4a}$, $C_{6,9}$-H, $C_{9a}$-$C_{5a}$, $N_5$-$C_{4a}$ | - | - | - |
| 1250 | $\nu_{53}$ | 1271 | $C_9$-H, $C_{11}$-$H_3$, $N_3$-H, $C_{10a}$-$N_1$, $C_{9a}$-$C_{5a}$, $C_{10a}$-$C_{4a}$, $N_5$-$C_{4a}$ | $\nu_{53}$ | 1319 | $C_{11}$-$H_3$, $N_{10}$-$C_{10a}$, $C_{9a}$-$C_{5a}$, $N_5$-$C_{4a}$, $C_8$-$C_7$, $C_4$-$N_3$, $C_9$-$C_{9a}$ |
| 1338 | - | - | - | $\nu_{54}$ | 1344 | $C_{5a}$-$N_1$, $C_9$-$C_{9a}$, $C_{10a}$-$C_{4a}$, $N_{10}$-$C_{10a}$, $N_3$-H, $C_{7a}$-$H_3$, $N_3$-$C_2$ |
| | $\nu_{56}$ | 1344 | $N_3$-H, $C_{7a,11}$-$H_3$, $C_6$-$C_{5a}$, $C_6$-H, $C_{9a}$-$N_{10}$, $C_2$-$N_3$ | - | - | - |

| | | | | | | |
|---|---|---|---|---|---|---|
| 1381 | $\nu_{58}$ | 1363 | $C_{8a,11}-H_3$, $C_{9a}-N_{10}$, $C_{10a}-C_{4a}$, $N_1-C_2$, $C_4-N_3$, $C_{5a}-N_5$ | - | - | - |
| | $\nu_{59}$ | 1365 | $C_{8a}-H_3$, $C_9-H$, $N_5-C_{4a}$, $C_{9a}-C_{5a}$ | $\nu_{57}$ | 1398 | $C_{11}-H_3$, $N_3-H$, $C_{10a}-N_1$, $C_{5a}-N_5$, $C_7-C_6$, $C_{9a}-N_{10}$ |
| | $\nu_{60}$ | 1368 | $N_3-H$, $C_6-H$, $C_{9a}-C_{5a}$, $C_{10a}-C_{4a}$, $C_4-N_3$ | $\nu_{58}$ | 1404 | $C_{8a,7a}-H_3$, $C_6-C_{5a}$, $C_8-C_7$, $C_{9a}-C_{5a}$ |
| | $\nu_{61}$ | 1380 | $C_{11,8a,7a}-H_3$, $C_{9a}-C_{5a}$, $C_8-C_7$, $C_{10a}-N_1$ | $\nu_{60}$ | 1426 | $C_{11,7a}-H_3$, $C_{5a}-N_5$, $N_{10}-C_{10a}$, $C_2-N_3$ |
| 1416 | $\nu_{64}$ | 1415 | $C_{7a}-H_3$, $N_5-C_{4a}$, $C_4-N_3$, $C_{10a}-N_1$ | $\nu_{64}$ | 1459 | $C_{8a}-H_3$, $C_{10a}-C_{4a}$, $N_3-H$, $C_2-N_3$, $C_{9a}-C_{5a}$, $N_1-C_2$ |
| | $\nu_{65}$ | 1428 | $C_{11,8a}-H_3$, $C_9-H$, $C_{10a}-C_{4a}$, $N_1-C_2$ | $\nu_{68}$ | 1484 | $C_{11}-H_3$, $N_{10}-C_{10a}$, $N_5-C_{4a}$, $C_{7a,8a}-H_3$, $N_1-C_2$ |
| 1498 | $\nu_{71}$ | 1511 | $C_8-C_7$, $C_{9a}-C_{5a}$, $C_{6,9}-H$, $C_{7a,8a}-H_3$, $C_6-C_{5a}$, $C_9-C_{9a}$, $C_{10a}-C_{4a}$ | $\nu_{72}$ | 1585 | $C_{10a}-N_1$, $N_5-C_{4a}$, $C_8-C_9$, $C_7-C_6$, $C_{9a}-C_{5a}$, $C_{6,9}-H$, $N_3-H$, $C_4-N_3$ |
| 1570 | $\nu_{73}$ | 1587 | $C_2-O_{2'}$, $N_3-H$, $C_6-C_{5a}$, $C_{9,6}-H$ | $\nu_{73}$ | 1656 | $C_8-C_9$, $C_6-C_{5a}$, $C_{9a}-N_{10}$, $C_7-C_6$, $C_{9,6}-H$, $C_{10a}-N_1$, $C_{4a}-C_4$ |
| | $\nu_{74}$ | 1608 | (s) $C_2-O_{2'}$, $C_4-O_{4'}$, $C_8-C_9$, $C_6-C_{5a}$, $C_{6,9}-H$, $C_{9a}-N_{10}$, $C_{10a}-N_1$ | $\nu_{74}$ | 1759 | $C_4-O_{4'}$, $N_3-H$, $C_2-O_{2'}$ (as) |
| 1626 | $\nu_{75}$ | 1645 | $C_4-O_{4'}$, $N_3-H$, $C_{4a}-C_4$, $C_6-C_{5a}$, $C_8-C_9$ | $\nu_{75}$ | 1782 | $C_2-O_{2'}$, $C_4-O_{4'}$, $N_3-H$ (s) |

| Exp. RR FMN $S_1$ | X3LYP/cc-pVDZ | | |
|---|---|---|---|
| | $\nu_\#$ | $S_1$offR | **Assignment** |
| 1200 | $\nu_{51}$ | 1232 | $C_2-N_3$, $N_3-H$, $C_{10a}-C_{4a}$, $C_9-H$, $C_{9a}-C_{5a}$, $N_{10}-C_{11}$ |
| | - | - | - |
| 1250 | $\nu_{53}$ | 1298 | $N_{10}-C_{10a}$, $C_{11}-H_3$, $C_{6,9}-H$, $C_8-C_7$, $N_5-C_{4a}$, $N_3-C_2$, $C_{10a}-C_{4a}$ |
| 1338 | $\nu_{54}$ | 1321 | $C_{5a}-N_5$, $C_{6,9}-H$, $C_9-C_{9a}$, $N_{10}-C_{10a}$, $C_{4a}-C_4$, $N_3-C_2$, $C_8-C_7$ |
| | $\nu_{55}$ | 1345 | $N_3-H$, $N_{10}-C_{10a}$, $C_{4a}-C_{10a}$, $C_{5a}-C_{9a}$, $C_7-C_6$, $C_{11}-H_3$, $C_6-H$ |
| 1381 | $\nu_{57}$ | 1383 | $N_3-H$, $C_6-H$, $C_{11}-H_3$, $C_7-C_6$, $C_{9a}-N_{10}$, $C_6-C_{5a}$, $C_{10a}-C_{4a}$, $C_4-N_3$ |
| | $\nu_{59}$ | 1394 | $C_{8a}-H_3$, $C_6-C_{5a}$, $C_8-C_9$, $N_1-C_2$, $C_4-N_3$ |
| | $\nu_{60}$ | 1403 | $C_{10a}-N_1$, $N_5-C_{4a}$, $C_{9a}-N_{10}$, $N_1-C_2$, $N_3-H$, $N_2-C_3$, $C_9-C_{9a}$, $C_{11,7a,8a}-H_3$ |
| | $\nu_{61}$ | 1410 | $N_3-H$, $C_4-N_3$, $C_6-H$, $C_{9a}-C_{5a}$, $N_5-C_{4a}$, $C_{10a}-N_1$ |
| 1416 | $\nu_{64}$ | 1444 | $N_5-C_{4a}$, $C_{10a}-N_1$, $C_4-N_3$, $C_2-N_3$, $C_{7a,11}-H_3$, $C_8-C_7$ |
| | $\nu_{65}$ | 1447 | $C_{8a,11}-H_3$, $C_{10a}-C_{4a}$, $N_1-C_2$, $C_8-C_9$, $C_7-C_6$, $C_{4a}-C_4$ |
| 1498 | $\nu_{71}$ | 1527 | $C_8-C_7$, $C_9-C_{9a}$, $C_{9a}-C_{5a}$, $C_6-C_{5a}$, $C_{10a}-C_{4a}$, $C_{6,9}-H$, $C_{7a}-H_3$ |
| 1570 | $\nu_{73}$ | 1635 | $C_8-C_9$, $C_6-C_{5a}$, $C_{6,9}-H$, $C_7-C_6$, $N_{10}-C_{9a}$, $N_5-C_{4a}$, $C_{10a}-N_1$ |
| | $\nu_{74}$ | 1701 | $C_2-O_{2'}$, $N_3-H$, $C_4-O_{4'}$ (as), $C_{4a}-C_{10a}$, $C_4-N_3$ |
| 1626 | $\nu_{75}$ | 1726 | $C_4-O_{4'}$, $C_2-O_{2'}$ (s), $N_3-H$, $C_{4a}-C_4$ |
